# Simultaneous, real-time tracking of many neuromodulatory signals with Multiplexed Optical Recording of Sensors on a micro-Endoscope

**DOI:** 10.1101/2025.01.26.634931

**Authors:** Peter N. Kalugin, Paul A. Soden, Crystian I. Massengill, Oren Amsalem, Marta Porniece, Diana C. Guarino, David Tingley, Stephen X. Zhang, Jordan C. Benson, Madalon F. Hammell, David M. Tong, Charlotte D. Ausfahl, Tiara E. Lacey, Ya’el Courtney, Alexandra Hochstetler, Andrew Lutas, Huan Wang, Lan Geng, Guochuan Li, Bohan Li, Yulong Li, Maria K. Lehtinen, Mark L. Andermann

## Abstract

Dozens of extracellular molecules jointly impact a given neuron, yet we lack methods to simultaneously record many such signals in real time. We developed a probe to track ten or more neuropeptides and neuromodulators using spatial multiplexing of genetically encoded fluorescent sensors. Cultured cells expressing one sensor at a time are immobilized at the front of a gradient refractive index (GRIN) lens for 3D two-photon imaging *in vitro* and *in vivo.* The sensor identity and detection sensitivity of each cell are determined via robotic dipping of the probe into wells containing various ligands and concentrations. Using this probe, we detected stimulation-evoked release of multiple neuromodulators in acute brain slices. We also tracked endogenous and drug-evoked changes in cerebrospinal fluid composition in the awake mouse lateral ventricle, which triggered downstream activation of the choroid plexus epithelium. Our approach offers a first step towards quantitative, real-time, high-dimensional tracking of brain fluid composition.

## Introduction

Brain cells exist in a rich environment of neuromodulators, peptides, hormones, and other molecular signals derived from both local and distant sources. Changes in these extracellular signals on timescales ranging from milliseconds to hours underlie fundamental processes in brain function and physiology, including learning and motivated behavior, circadian and diurnal rhythms, development, inflammation and degeneration^1–6^. Dozens of such signals travel through brain interstitial fluid and cerebrospinal fluid (CSF) to engage receptors in neurons, glia, immune cells, and brain barriers such as the choroid plexus (ChP) epithelium^7–10^. Further, many neurons express a dozen or more receptors for these signals at once^11–15^. Recent functional studies have begun to unravel the complex interplay of parallel neuromodulatory signaling axes in the brain previously predicted by genomic studies^16^. Despite clear evidence of their dynamic and complementary effects on various tissues, rapid fluctuations in these signals have typically been studied for one signal at a time due to technical limitations^17,18^. Moreover, neuromodulatory ligands impact brain physiology not simply by their presence or absence but through their time-varying levels^19,20^. While pharmacological replacements of neurochemicals, such as vasopressin in autism spectrum disorder and L-DOPA in Parkinson’s disease, are compelling treatments, questions remain about their precise trafficking routes, quantitative pharmacodynamics in target brain regions, and the potential for more titrated dosing using closed-loop control^21,22^. Polypharmacy of prescription medications, both psychotropic and otherwise, continues to rise, with minimal mechanistic understanding of combinatorial drug effects on brain activity^23,24^. While a complete accounting of the coordinated dynamics of many molecular signals in brain fluids is a technically challenging goal, it could yield profound mechanistic insights into the nature of brain states, as well as new therapeutic opportunities.

Techniques to study the composition of the extracellular environment of brain cells include sampling and offline molecular analysis, electrochemistry and field-effect transistors, and optical methods, each with powerful capabilities but also significant limitations. For instance, microdialysis collection followed by mass spectrometry allows up to hundreds of molecules to be measured at the same time with exceptional sensitivity and quantitative precision^25^. However, microdialysis typically requires collection and analysis times of minutes (but see Wells *et al*., 2024^26^), thereby missing more rapid signaling and precluding real-time readout of concentrations, as well as risking analyte dilution and degradation in the time prior to and during *ex vivo* sample processing. Electrochemical methods such as fast-scan cyclic voltammetry provide more rapid readouts of signals *in situ*, but are challenging to expand beyond the small set of molecules with measurable redox activity and suffer from significant crosstalk between ligands of related chemical structure^27–32^. Aptamer-field-effect transistor devices are also capable of fast and sensitive molecular measurements but have so far been fragile and prone to biofouling, hindering their effective application in complex biological fluids and living systems^33,34^.

In recent years, optical methods such as reporter dyes and genetically encoded indicators have emerged as popular and versatile tools for tracking molecular concentrations in the brain^35–38^. Indicators for extracellular signals based on G protein-coupled receptors (GPCRs) and bacterial periplasmic binding proteins have now been developed for over two dozen neuromodulators and neuropeptides^18,39–46^. These sensors can detect concentrations in the nanomolar range and are able to distinguish signal transients relevant to brain circuit functions *in vivo* at subsecond resolution. Some such sensors have recently been demonstrated to possess measurable fluorescence lifetime signals, opening the possibility of chronic recordings over days and weeks^47^. Despite these major achievements, the utility of genetically encoded indicators faces two important challenges: (i) they do not readily provide a quantitative readout of analyte concentration, even in the case of fluorescence lifetime sensors, due to the uncertainty of extrapolating *in vitro* calibrations to sensors expressed in brain tissue, and (ii) most current sensors are green, and their broad fluorescence excitation and emission spectra limit the number of simultaneously measurable sensors to 2-3 even in the rare cases where spectrally distinct sensors are available^46,48^.

To begin to address these challenges, we developed Multiplexed Optical Recording of Sensors on a micro-Endoscope (MORSE) – an optical probe capable of simultaneously recording ten or more molecular signals in brain and body extracellular fluids with nanomolar sensitivity, time resolution of seconds and real-time readout. Our probe uses a stable, porous hydrogel containing a mixture of cultured cells expressing different green fluorescent neuromodulator or neuropeptide sensors one at a time. A thin (≤200 μm) layer of this hydrogel is immobilized at the front of a half-millimeter diameter gradient refractive index (GRIN) lens. This allows 3D imaging of all green fluorescent sensors *in vitro* or *in vivo* followed by estimation of sensor identity and quantitative calibration via post hoc robotic dipping of the probe into fluid samples containing various ligands and concentrations. Notably, this approach is scalable up to tens of sensors and amenable to two-photon imaging using standard microscopes. We first applied MORSE to simultaneously and quantitatively decode the concentrations of up to ten molecular signals in fluids *in vitro*, with successful unmixing of crosstalk at sensors that exhibit partial sensitivity to multiple ligands. Next, we used the probe to detect distinct quantitative temporal dynamics of release of multiple neuromodulators from different brain nuclei in acute brain slices. Finally, we deployed MORSE in the lateral ventricle of an awake mouse. There, we tracked the pharmacodynamics of three or more signals simultaneously during sequential peripheral ligand delivery and trafficking into the CSF. We additionally observed spontaneous fluctuations in CSF serotonin at a level sufficient to drive second messenger activity in the CSF-facing ChP epithelial cells. These findings demonstrate MORSE as a flexible approach for combining a broad range of genetically encoded fluorescent sensors for high-dimensional interfacing with complex chemical environments.

## Results

### MORSE probe development and characterization

Preparation of MORSE probes involved two fabrication stages. First, a standard half-millimeter diameter gradient refractive index (GRIN) lens was secured in a piece of polyimide tubing with RTV silicone elastomer. The silicone was shaped to form a curved well at the front of the lens extending from the edges of the tubing to a depth of ∼200 µm at the center (**Figure 1A, left**). Next, HEK-293T cells were transfected with plasmids encoding one of up to 16 different green fluorescent protein (GFP)-based GPCR activation-based (GRAB) sensors for different neuromodulators and neuropeptides^45,49–58^. We then mixed these “sensor” cells together in a hydrogel consisting of extracellular matrix components collagen I and Matrigel and applied this mixture to the front of the GRIN lens (**Figures 1A, right and 1B**). 3D two-photon imaging of the resultant hydrogel through the GRIN lens (30 planes/volume, ∼600×600×270 µm^3^ volume) demonstrated a dispersed population of 100-200 green cells spanning the full thickness of the gel (**Figure 1C, left**). The total active probe preparation time requires approximately 3 hours, and the process is scalable to many probes in parallel with minimal additional time.

**Figure 1.**
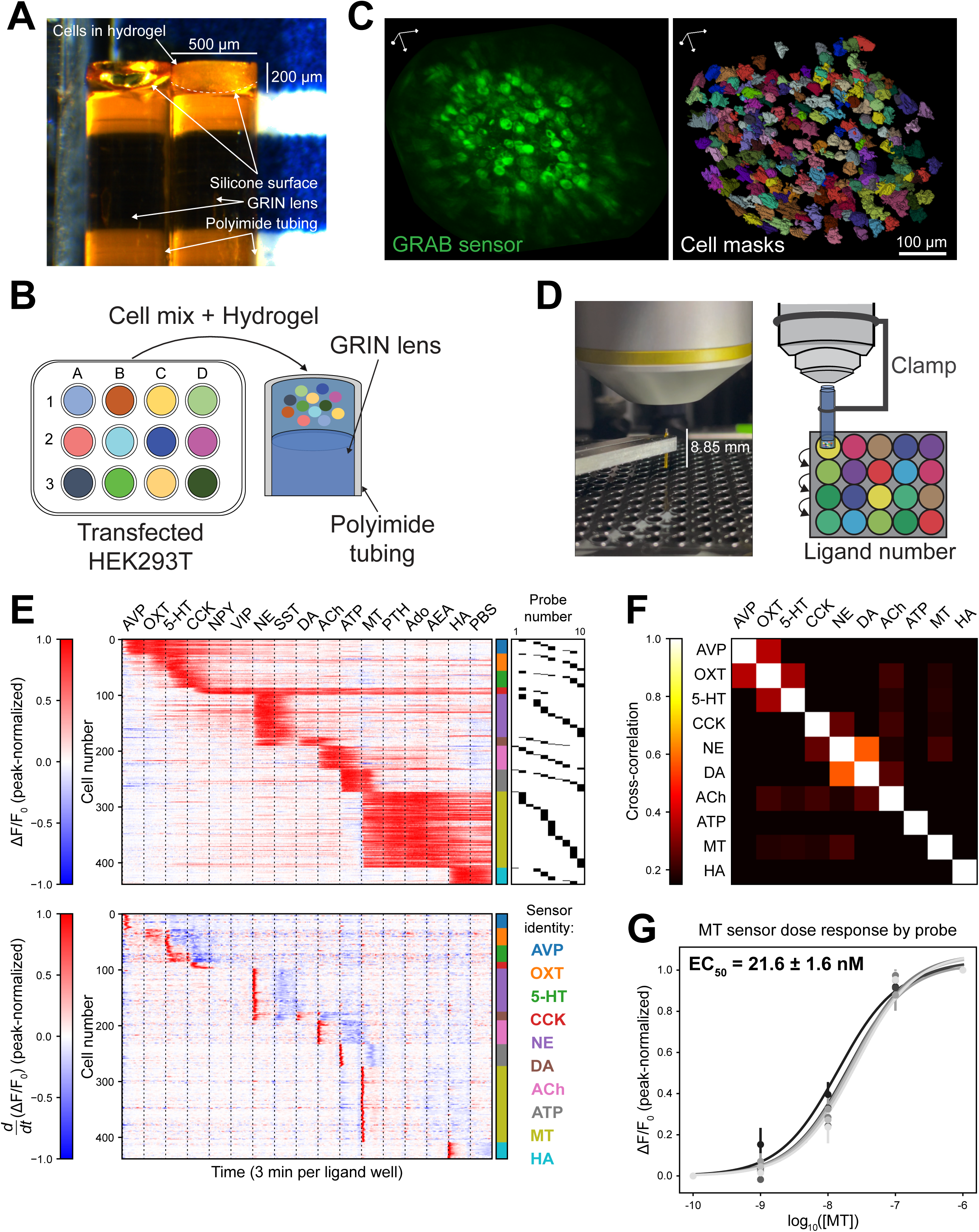
MORSE probe design. (**A**) Photo of MORSE probes before (left) and after (right) hydrogel deposition. (**B**) Cultured HEK-293T cells are transfected with one plasmid vector expressing a green GRAB sensor, so that each cell expresses only one type of sensor. Twenty-four hours later, cells expressing up to 16 individual sensor types are pooled, mixed into an optimized hydrogel matrix, and attached to the front of a half-millimeter diameter gradient refractive index (GRIN) lens outfitted with a receptacle built with polyimide tubing and RTV silicone. (**C**) **Left:** 3D rendering of a single two-photon imaging volume collected from a MORSE probe through the GRIN lens, demonstrating a dispersed population of HEK-293T cells expressing green sensor proteins. **Right:** single-cell masks calculated from this MORSE probe, demonstrating recovery of individual cell shapes in 3D. Scale bar: 100 μm. (**D**) Summary of the dipping characterization step. **Left:** a MORSE probe is secured on the microscope with a custom lens clamp above a 384-well plate carrying ∼20 μL wells of solutions carrying various ligands at different concentrations. **Right:** the MORSE probe is subsequently robotically dipped for several minutes at a time into one well of the plate after another, with 3D volumetric two-photon imaging performed throughout. Each well drives a different pattern of sensor cell activation based on the ligand composition in the well, allowing both sensor identities and dose-response relationships to be recovered from each sensor cell on the MORSE probe. (**E**) 442 single-cell ROIs were recovered from n=10 MORSE probes, each carrying 16 GRAB sensor cell types and dipped into high concentrations of their 16 respective ligands: vasopressin (AVP, 1 μM), oxytocin (OXT, 1 μM), serotonin (5-HT, 1 μM), cholecystokinin (CCK, 1 μM), neuropeptide Y (NPY, 1 μM), vasoactive intestinal peptide (VIP, 1 μM), norepinephrine (NE, 10 μM), somatostatin (SST, 1 μM), dopamine (DA, 1 μM), acetylcholine (ACh, 1 mM), adenosine triphosphate (ATP, 1 mM), melatonin (MT, 10 μM), parathyroid hormone (PTH, 1 μM), adenosine (Ado, 1 mM), anandamide (AEA, 1 mM), and histamine (HA, 1 mM), followed by a wash well of phosphate-buffered saline (PBS). Each probe was dipped into a 20 μL volume of each ligand for 3 minutes prior to movement into the next well. **Top:** heatmap depicting peak-normalized ΔF/F_0_ signals for each individual cell following background correction. The identity of the probe that each cell is derived from is depicted in the black raster plot on the far right, and the sensor identity of the cell is shown with the color bar corresponding to the sensor names in the legend on the right. **Bottom:** heatmap depicting peak-normalized time-derivatives of the single-cell ΔF/F_0_ signals shown in the top heatmap. The cells from all probes are listed in order of activation during the recording, demonstrating robust activation of up to 10/16 sensors by their respective ligands (median of 8 active sensors per probe, interquartile range 5-9; average of 6.1 cells per sensor per probe). (**F**) Cross-correlation of the mean time-derivative traces for each sensor group in **E** (**bottom**), which are depicted individually in **Fig. S3A**. Robust crosstalk is evident between the NE and DA sensors, along with weaker crosstalk between AVP and OXT sensors. Additional mild crosstalk between OXT/5-HT and CCK/NE sensor pairs likely reflects the slower wash-on and wash-off kinetics of the OXT and CCK sensors, and in the former case is not replicated in an independent set of probes in **Figs. S3B-D**. (**G**) Dose-response activity of GRAB melatonin sensor across n=7 MORSE probes (8-27 cells per probe, most abundant sensor across probes) plotted individually. In probes with robust cell numbers (≥8 cells/probe) responding to melatonin doses, the sensor demonstrated consistent EC_50_ values around 20 nM. Error bars: ±SEM across sensor cells in each probe, error interval: ±SEM across probes.

We developed optimized strategies for many steps in the probe fabrication process, which should also be of broad value to the design of other gel-brain interfaces. We chose to use extracellular matrix-based hydrogels to achieve optimal biocompatibility and porosity to allow passage of molecules from the brain^59^. To assess hydrogel stability, we first performed time-lapse 3D imaging of the hydrogel (∼600×600×270 µm^3^ volumes, 30 planes/volume, ∼0.5 volumes/s) while the probe was stationary in phosphate buffered saline (PBS). We found that the GRAB sensors exhibited a wide range of baseline fluorescence intensities, resulting in high variability of sensor cell brightnesses, so we first preprocessed the recordings by performing 3D cell segmentation of each volume using Cellpose2.0 (**Figure S1A**)^60–62^. This generated a probability map quantifying how likely each voxel was to be part of a fluorescent sensor cell at each timepoint, a measure largely independent of fluorescence brightness (**Figure S1B**). To quantify hydrogel deformation, we next performed nonrigid registration on these probability maps to a single reference timepoint using optical flow-based warping (**Figure S1C**)^63–66^. We found that the small hydrogel volume (∼100 nL) on the MORSE probes, when composed of pure Matrigel, was not sufficiently stable to withstand even static imaging on the GRIN lens and exhibited substantial global deformation as well as some local cell movement over the course of tens of minutes (**Figure S1D; Supplementary Video 1, left**). To reduce cell movement, we incubated the cells with colchicine, a microtubule polymerization inhibitor and chemotaxis blocker, prior to harvesting^67^. We also introduced collagen I, a more rigid extracellular matrix component, to the hydrogel mixture, which stabilized the probes and minimized gel deformation (**Figure S1D; Supplementary Video 1, right**)^68^.

To determine the identity, crosstalk and sensitivity of the sensor expressed in each cell, we next performed time-lapse 3D imaging during several hours of sequential robotic dipping of the probe into several dozen wells of a 384-well plate containing ∼20 µL samples of known solutions (**Figure 1D; Supplementary Video 2**). During this protocol, which involves frequent probe-liquid contact cycles, we observed translational hydrogel shifts of large magnitudes not seen in the stationary imaging case, at times resulting in complete gel detachment and likely reflecting insufficient gel adhesiveness to the RTV silicone coating (**Figure S1E, left; Supplementary Video 3, left**). These deformations and detachments were strongly reduced by incubating the silicone surface with potassium hydroxide (KOH) for surface activation prior to gel administration (**Figures S1E-G; Supplementary Video 3, middle and right**)^69,70^. We settled on 1 hour of cell incubation with 1 μM colchicine and 1 hour of surface treatment with 2.5% KOH followed by application of a 60% collagen I:40% Matrigel hydrogel formulation, which was sufficiently porous to allow rapid ligand diffusion (see below) while exhibiting optimal rigidity and adhesiveness, resulting in high stability probes with relatively little movement.

Despite these improvements, slow residual hydrogel deformation occurred across tens of minutes. To account for this, we began by applying the hydrogel deformations calculated as above to stabilize the raw fluorescence images (**Figure S2A**). In a subset of probes, we prepared hydrogels with 10% of the cells expressing cytosolic tdTomato, performed two-color imaging, and compared the results of image alignment using the ligand-sensitive green GRAB channel or the ligand-insensitive red channel (**Figure S2B**). We found that registration using Cellpose2.0 probability maps derived from the GRAB signals performed similarly to or better than the probability maps from the tdTomato signals in all classes of experiments that we performed (**Figure S2C; Supplementary Video 4**). This demonstrates segmentation probability map filtering as a generalizable approach for extracting structural information from two-photon imaging datasets with large dynamic range of cell-to-cell signal intensities^71^.

To extract signals from individual cells, we next applied Caiman, a constrained non-negative matrix factorization algorithm, to define 3D masks based on correlated local fluorescence fluctuations^72^. We performed semi-automated curation of the resulting masks using a custom GUI^73^. Many of the MORSE probes were imaged multiple times in several contexts and experiments, such as in biological samples followed by *ex vivo* calibration. In these cases, we also matched cells across these recordings. To account for 3D rotations and deformations between different recordings of the same probe, we aligned representative single volumes of each recording using manually defined landmarks (**Figure S2D**)^74,75^. These alignments were used to match overlapping cell masks between the two recordings (**Figure S2E**). With this approach, we were able to reliably track an average of 70-130 active sensor cells per probe across 2-8 sequential recordings in several different contexts lasting up to 8 hours in total (an initial 1-5 hr recording involving (i) insertion into a brain slice, (ii) *in vivo* in lateral ventricle CSF, or (iii) dipping into a set of *ex vivo* fluid samples of interest, followed by a dipping characterization to identify the sensor identities of each cell and then up to six additional dipping experiments into various sets of ligand dilutions and fluid samples of 30-90 min each) (**Figures 1C, right and S2F**).

We began by dipping 16-sensor MORSE probes sequentially into solutions containing near-saturating concentrations of each of 16 individual sensor ligands, one ligand at a time (**Table 1**). This resulted in rapid, robust, and consistent patterns of elevated fluorescence in subsets of cells in response to up to ten of the ligands in each probe (**Figures 1E and S3A; Supplementary Video 5**). We were unable to detect robust fluorescence responses to the other six ligands, likely owing to a combination of variations in sensor sensitivity and density of cells on the probe expressing each sensor. By performing hierarchical clustering on the rise time of the individual sensor cell signals, we were able to assign clear sensor identities across several hundred cells in ten representative MORSE probes (median of 8 active sensors per probe, interquartile range 5-9; average of 6.1 cells per sensor per probe) (**Figure 1E, color bar**). Significant crosstalk was detectable most prominently between dopamine and norepinephrine sensors, and to a lesser extent between vasopressin and oxytocin sensors, as reported previously (**Figures 1F and S3B-E,** top)^43,58^.

**Table 1.**
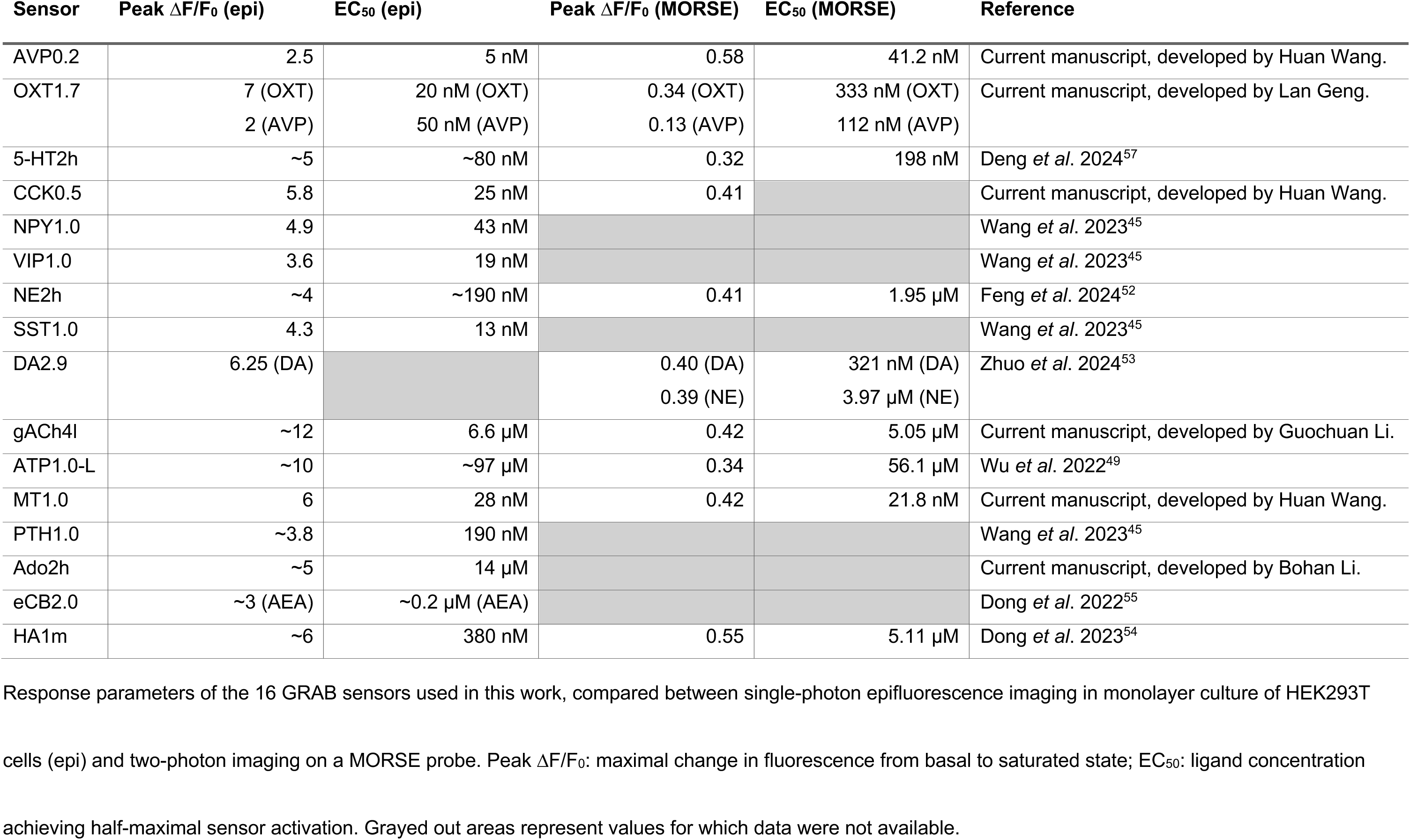
Summary of GRAB sensors used in this work.

| Sensor | Peak $\Delta F/F_0$ (epi) | EC<sub>50</sub> (epi) | Peak $\Delta F/F_0$ (MORSE) | EC<sub>50</sub> (MORSE) | Reference |
| --- | --- | --- | --- | --- | --- |
| AVP0.2 | 2.5 | 5 nM | 0.58 | 41.2 nM | Current manuscript, developed by Huan Wang. |
| OXT1.7 | 7 (OXT) | 20 nM (OXT) | 0.34 (OXT) | 333 nM (OXT) | Current manuscript, developed by Lan Geng. |
|  | 2 (AVP) | 50 nM (AVP) | 0.13 (AVP) | 112 nM (AVP) |  |
| 5-HT2h | ~5 | ~80 nM | 0.32 | 198 nM | Deng *et al.* 2024<sup>57</sup> |
| CCK0.5 | 5.8 | 25 nM | 0.41 |  | Current manuscript, developed by Huan Wang. |
| NPY1.0 | 4.9 | 43 nM |  |  | Wang *et al.* 2023<sup>45</sup> |
| VIP1.0 | 3.6 | 19 nM |  |  | Wang *et al.* 2023<sup>45</sup> |
| NE2h | ~4 | ~190 nM | 0.41 | 1.95 μM | Feng *et al.* 2024<sup>52</sup> |
| SST1.0 | 4.3 | 13 nM |  |  | Wang *et al.* 2023<sup>45</sup> |
| DA2.9 | 6.25 (DA) |  | 0.40 (DA) | 321 nM (DA) | Zhuo *et al.* 2024<sup>53</sup> |
| | | | 0.39 (NE) | 3.97 μM (NE) | |
| gACh4I | ~12 | 6.6 μM | 0.42 | 5.05 μM | Current manuscript, developed by Guochuan Li. |
| ATP1.0-L | ~10 | ~97 μM | 0.34 | 56.1 μM | Wu *et al.* 2022<sup>49</sup> |
| MT1.0 | 6 | 28 nM | 0.42 | 21.8 nM | Current manuscript, developed by Huan Wang. |
| PTH1.0 | ~3.8 | 190 nM |  |  | Wang *et al.* 2023<sup>45</sup> |
| Ado2h | ~5 | 14 μM | | | Current manuscript, developed by Bohan Li. |
| eCB2.0 | ~3 (AEA) | ~0.2 μM (AEA) | | | Dong *et al.* 2022<sup>55</sup> |
| HA1m | ~6 | 380 nM | 0.55 | 5.11 μM | Dong *et al.* 2023<sup>54</sup> |

Response parameters of the 16 GRAB sensors used in this work, compared between single-photon epifluorescence imaging in monolayer culture of HEK293T cells (epi) and two-photon imaging on a MORSE probe. Peak $\Delta F/F_0$ : maximal change in fluorescence from basal to saturated state; EC<sub>50</sub>: ligand concentration achieving half-maximal sensor activation. Grayed out areas represent values for which data were not available.

Once the sensor identity of each cell was defined in this way, we next performed dose-response calibrations by dipping the probe sequentially into wells containing mixtures of ligands, with increasing concentrations of each ligand in successive wells (**Supplementary Video 6**). This allowed us to assess probe sensitivity to all ligands in a reasonable amount of time. We found that EC_50_ values, the sensors’ half-maximal effective concentrations (EC_50_ is equivalent to a sensor’s apparent dissociation constant K_d_), could be reliably estimated for nine of the ten sensors for which robust responses were observed (**Figures 1G and S3F**). Depending on the sensor, EC_50_ values could be very sensitive (e.g. tens of nanomolar range). The exception was the CCK0.5 sensor, which was identified in small numbers of cells on only n=3/10 probes and demonstrated unusually long signal wash-off times following dipping, likely reflecting unfavorable ligand dissociation characteristics in this version of the protein^45^. Importantly, while each sensor exhibited a relatively broad range of peak response magnitudes across individual cells and probes, normalizing each cell’s signal to its maximal observed response and averaging that across many cells per sensor per probe produced remarkably consistent dose-response relationships (**Figures 1G and S3E-F**). This highlights an intrinsic benefit of the MORSE strategy, which minimized the influence of variability in baseline expression level, autofluorescence, optical aberrations of the GRIN lens, and other parameters across individual sensor cells on a probe by pooling signals from multiple cells on each probe to estimate the ligand concentration. Using these parameters, we were able to derive estimates of the quantitative concentrations of each ligand at the probe. Furthermore, by observing the time course of the response to each concentration as it reached a plateau, and assuming that ligand binding at the sensor is fast (τ on the order of 1 sec or less in all GRAB sensors reported so far), we could assess the diffusion time constant through the hydrogel layer, which depended on ligand identity and plateaued within 1 min for most ligands (**Figure S3E, bottom**)^45,49–58^.

In summary, these *in vitro* experiments establish the feasibility of MORSE for quantitatively tracking ten or more green fluorescent sensors at once – an order of magnitude more sensors than is possible with current methods. In combination with our customized 3D image analysis pipeline, MORSE provides a promising platform for scalable multiplexed ligand sensing in fluids. We proceeded to apply MORSE to study the mouse brain in live slices and *in vivo*.

### MORSE detects regionalized neuromodulator release in brain slices

We first tested MORSE’s ability to detect endogenous release of multiple signals in brain slices. For this application, we created probes using seven of the most robust sensors: norepinephrine (NE), dopamine (DA), serotonin (5-HT), acetylcholine (ACh), vasopressin (AVP), oxytocin (OXT), and histamine (HA). We gently pressed the probes gel-first against specific regions of the brain slice containing varying levels of neuromodulatory inputs. Then, we collected 3D two-photon images while performing bath applications of potassium chloride (KCl) to depolarize the neurons and evoke neuromodulator release, followed by bath applications of saturating doses of each ligand.

We began by recording in the dorsolateral geniculate nucleus (dLGN). This thalamic nucleus is locally enriched in serotonergic and noradrenergic synapses^76,77^. We therefore targeted the MORSE probes to the dLGN by visualizing serotonergic synapses in the dLGN after expressing a virus encoding tdTomato in serotonin neurons of the dorsal raphe nucleus (DRN; in *Pet1-Cre* mice) (**Figure 2A**). Upon bath application of 50 mM KCl, we detected activation of the GRAB sensors for NE, DA, and 5-HT (**Figures 2B-C, top; Supplementary Video 7**). We directly confirmed the identity of these sensors by subsequent bath application of the corresponding ligands on the slice. In a subset of experiments (n=4/9), we additionally performed a dipping characterization of the probe following the slice experiment (**Figure S4A; Supplementary Video 8**). Dipping the probe sequentially into wells containing NE, DA or 5-HT evoked robust responses in different subsets of several dozen cells matching the cells previously imaged on the slice. These data highlight our capacity to record from this probe for 1-2 hours in a brain slice followed by accurate alignment to data collected during 1.5 additional hours of dipping (**Figure 2B**). The dipping also identified many additional cells expressing sensors for AVP, OXT, and HA. Most of these cells did not have matching cell masks in the slice experiment, indicating that they did not respond to bath application of NE, DA, or 5-HT, and KCl did not elicit sufficient release of AVP, OXT, or HA from the slice to drive the respective sensors.

**Figure 2.**
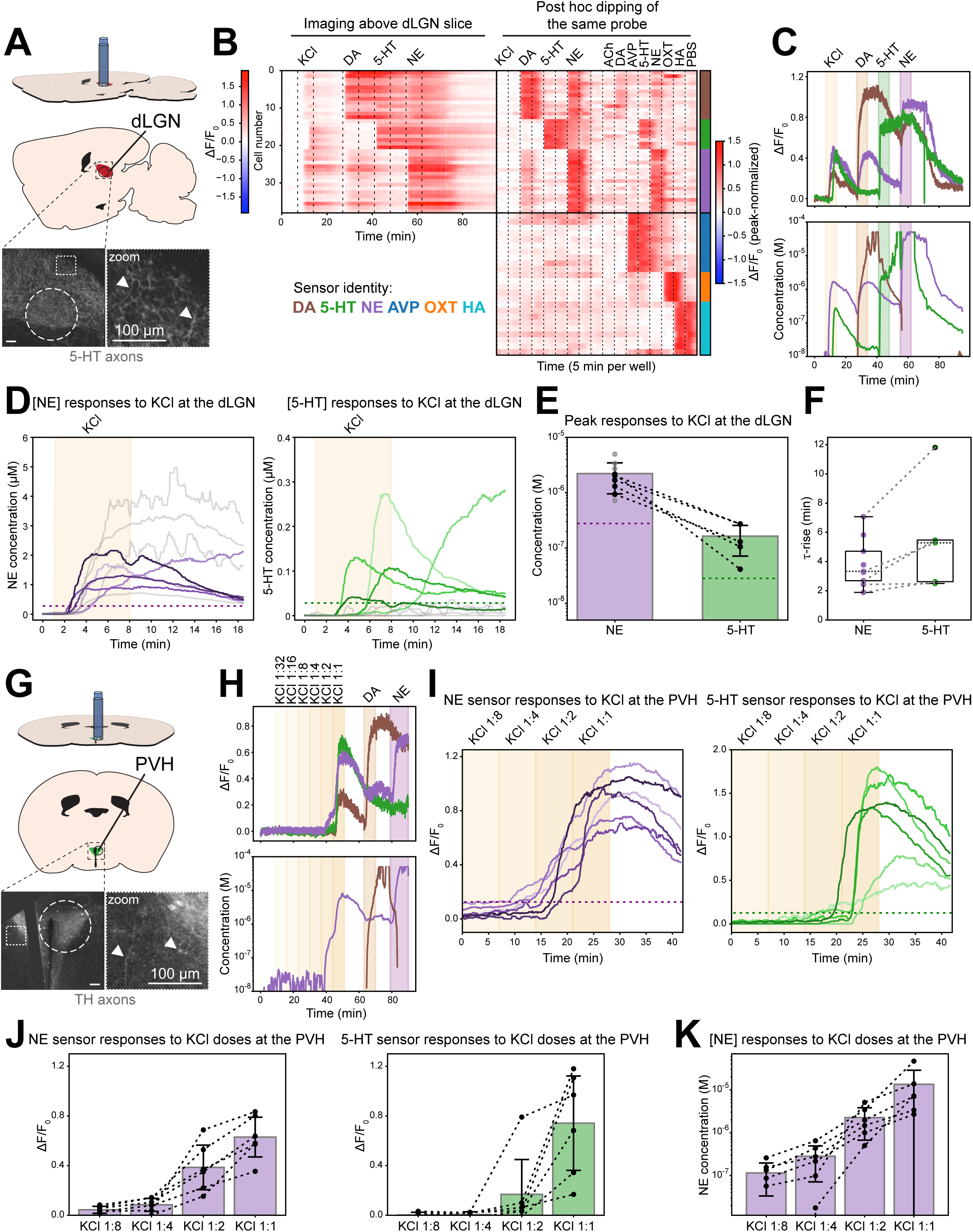
MORSE recordings from live brain slices. (**A**) Schematic of the dorsolateral geniculate nucleus (dLGN) slice recording setup. The dLGN was identified in sagittal slices of the mouse brain by observing tdTomato expression in serotonergic neurons from *Pet1-Cre* animals. **Inset, left:** two-photon image of tdTomato expression at an identified dLGN region, with the MORSE probe location on the slice indicated by the dashed circle. **Inset, right:** zoomed in region of the slice corresponding to the dotted square in **inset, left**, indicating individual serotonergic axons with white arrowheads. Scale bars: 100 μm. (**B**) Single-sensor cell traces from one seven-sensor (ACh, AVP, OXT, 5-HT, NE, DA, HA) MORSE probe recorded in the dLGN (left heatmap, ΔF/F_0_ depicted as per left color bar) and subsequently dipped into standard solutions (right heatmap, peak-normalized ΔF/F_0_ depicted as per right color bar). The slice was continuously perfused with warm aCSF, which was periodically replaced with aCSF containing KCl (50 mM), DA (50 μM), 5-HT (50 μM), and NE (50 μM) at the indicated times. During dipping, the probe was first dipped into these same solutions with interposed dips into wells containing aCSF in between them (left half of right heatmap) and subsequently dipped into standard solutions of the seven ligands corresponding to each sensor formulated in PBS (1 mM ACh, 1 μM DA, 1 μM AVP, 1 μM 5-HT, 10 μM NE, 1 μM OXT, 1 mM HA). The upper half of the right heatmap demonstrates the sensor cells with ROIs successfully matched between the slice recording and the dipping characterization, and the lower half shows additional cells that were inactive during the slice recording but responded to dipping. The sensor identity of each cell is shown with the color bar on the right as per legend. (**C**) **Top:** mean ΔF/F_0_ traces for each sensor type from the left heatmap in **B**, with ligand wash-on timings indicated by colored shading and sensors color-coded as in **B**. **Bottom:** estimated ligand concentration traces calculated from the ΔF/F_0_ traces above, color-coded in the same fashion. Notably, unmixing of NE and DA ligand contributions to the response of the DA sensor demonstrates that the KCl wash-on response of that sensor is solely due to NE release. There is also some DA ligand contribution to the NE signal at the time of 50 µM DA wash-on, but we could not unmix this effect because we only tested DA doses up to 1 µM in the dose-response calibrations of the probes. (**D**) NE (**left**) and 5-HT (**right**) concentrations measured by n=9 MORSE probes positioned over the dLGN during KCl administration. In n=4/9 recordings (gray traces in both plots), peak 5-HT responses were below 28.3 nM, corresponding to a peak-normalized ΔF/F_0_ signal of 0.125, which was arbitrarily set in these recordings to distinguish strong effects. The remaining n=5/9 recordings demonstrated strong release of both NE and 5-HT in response to KCl administration. These are plotted with matching hue depths for comparison (e.g. the darkest purple NE curve corresponds to the darkest green 5-HT curve). Colored dotted lines on both plots demonstrate the peak-normalized ΔF/F_0_ = 0.125 thresholds (corresponding to 0.278 µM for NE and 28.3 nM for 5-HT). (**E**) Quantification of the peak NE and 5-HT concentrations elicited by KCl administration in **D**, with values from each of the n=5/9 recordings demonstrating strong release of both ligands plotted in black and connected with dotted lines, and the remaining n=4/9 NE responses plotted in gray. Dotted lines on both colored bars correspond to the peak-normalized ΔF/F_0_ = 0.125 thresholds (0.278 µM for NE and 28.3 nM for 5-HT). Error bars: ±SEM across recordings. (**F**) Quantification of 1-rise onset timings of the NE and DA concentration time courses in **D**, with values from each of the n=5/9 recordings demonstrating strong release of both ligands connected by dotted lines. (**G**) Schematic of the paraventricular nucleus of the hypothalamus (PVH) slice recording setup. The PVH was identified in coronal slices of the mouse brain by observing YFP expression in monoaminergic neurons from *TH-Cre* animals. **Inset, left:** two-photon image of YFP expression at an identified PVH region, with the MORSE probe location on the slice indicated by the dashed circle. **Inset, right:** zoomed in region of the slice corresponding to the dotted square in **inset, left**, indicating individual monoaminergic axons with white arrowheads. Scale bars: 100 μm. (**H**) A seven-sensor (ACh, AVP, OXT, 5-HT, NE, DA, HA) MORSE probe was recorded in the PVH and subsequently dipped into various solutions for sensor identification. The slice was continuously perfused with warm aCSF, which was periodically replaced with aCSF containing a dilution range of KCl concentrations (1:32, 1:16, 1:8, 1:4, and 1:2 dilutions of a 50 mM stock of KCl in aCSF as well as the full stock concentration labeled 1:1), DA (50 μM), and NE (50 μM) at the indicated times. **Top:** mean ΔF/F_0_ traces for each sensor type from this probe recorded at the PVH, corresponding to the left heatmap in **Fig. S4E**, with ligand wash-on timings indicated by colored shading and sensors color-coded as in **B**. **Bottom:** estimated ligand concentration traces calculated from the ΔF/F_0_ traces above, color-coded in the same fashion. Notably, unmixing of NE and DA ligand contributions to the response of the DA sensor demonstrates that the KCl wash-on response of that sensor is solely due to NE release. There is also some DA ligand contribution to the NE signal at the time of 50 µM DA wash-on, but we could not unmix this effect because we only tested DA doses up to 1 µM in the dose-response calibrations of the probes. (**I**) NE (**left**) and 5-HT (**right**) sensor ΔF/F_0_ signals measured by n=6 MORSE probes positioned over the PVH during administration of escalating doses of KCl. All n=6/6 recordings demonstrated strong release of both NE and 5-HT in response to KCl administration, above a threshold peak-normalized ΔF/F_0_ signal of 0.125. These are plotted with matching hue depths for comparison (e.g. the darkest purple NE curve corresponds to the darkest green 5-HT curve). Colored dotted lines on both plots correspond to the peak-normalized ΔF/F_0_ = 0.125 thresholds. Notably, 5-HT sensor identity was confirmed by dipping in n=2/6 of these recordings and putative 5-HT sensor identity inferred in the others by a pattern of responding to KCl but neither DA nor NE wash-on. (**J**) Peak NE (**left**) and 5-HT (**right**) sensor ΔF/F_0_ signals elicited by escalating doses of KCl in **I**, with values corresponding to each recording connected by dotted lines across the KCl doses. Error bars: ±SEM across recordings. (**K**) Quantification of the peak NE concentrations elicited by escalating doses of KCl in **I**, with values corresponding to each recording connected by dotted lines across the KCl doses. Error bars: ±SEM across recordings.

Due to the known crosstalk of NE binding to many versions of both the GRAB-NE and GRAB-DA sensors, we assessed whether the signal in the DA sensor could be explained by the release of NE alone from the slice^43^. To do this, we modeled the DA sensor’s responses to both DA and NE ligands using dose-response calibration parameters, and obtained quantitative estimates of NE, DA, and 5-HT ligand concentrations at the probe over the course of each slice recording. This approach successfully recovered the saturating wash-on responses of all three ligands at 50 µM and, importantly, indicated that the responses to KCl at both the NE and DA sensors were likely due solely to release of NE, rather than a combination of DA and NE (**Figure 2C, bottom**).

Robust responses to KCl, which we defined as a peak-normalized ΔF/F_0_ sensor signal increase of at least 0.125, were detected for NE in n=9/9 slice experiments and for 5-HT in n=5/9 experiments (**Figure 2D**). KCl-evoked peak concentrations were estimated at around 2.21 µM in the case of NE and around 164 nM for 5-HT in the recordings with detectable 5-HT release (**Figure 2E**). We also found that the rise time of activation of the 5-HT sensor was consistently delayed relative to the rise time for the NE sensor by an average of 99 sec (**Figure 2F**). To further improve our quantitative estimates of NE and 5-HT release in response to KCl, we used the hydrogel diffusion time constants obtained for these sensors to perform deconvolution and correct for the smoothing of the signal time course caused by diffusion of signals through the gel (**Figures S3E, bottom and S4B**). This approach resulted in sharper and narrower peak responses for each ligand, suggesting that the true peak concentrations at the slice surface were up to 20% higher and occurred ∼35 sec sooner than those recorded at the sensors in the hydrogel (**Figures S4C-D**).

To test the ability of MORSE to detect regional differences in neuromodulation, we compared the above experiment in the dLGN to a similar experiment in the paraventricular nucleus of the hypothalamus (PVH). The PVH is known to be innervated by several monoaminergic projections and also contains a number of peptidergic populations, including ones capable of releasing ligands detectable using the same seven sensors used in the above experiments, such as AVP and OXT^78–84^. MORSE probes were targeted to the PVH by visualizing the region after expressing a virus encoding yellow fluorescent protein (YFP) in local monoaminergic neurons (in *TH-Cre* mice) (**Figure 2G**). We used the same set of sensors for imaging in PVH as we did for imaging in dLGN, but this time we also assessed for graded signal release by washing on increasing concentrations of KCl in a stepwise fashion. KCl bath application here again activated the GRAB sensors for NE, DA, and 5-HT (**Figures 2H, top and S4E**). Similar to our observations in the dLGN, quantitative estimation of NE and DA concentrations along with unmixing of their combined effects at the DA sensor suggested that NE was solely responsible for activating both sensors in PVH (**Figure 2H, bottom**).

While we only bath applied saturating doses of NE and DA (but not 5-HT or other ligands) during these slice recordings, hierarchical clustering on all sensor time courses revealed a subpopulation of sensor cells responding to KCl but not to NE or DA (**Figure S4E**). When we performed dipping characterization of these probes following a subset of PVH slice experiments (n=2/6), we found that this third group of KCl responsive cells consistently aligned with 5-HT sensor cells. Furthermore, dipping directly into the KCl dilutions drove no responses in any sensor cells at any dosage, confirming that the KCl effects were due to evoked ligand release from the slice rather than direct KCl action at the sensors (**Figure S4F**). By the same definition of robust activation as above (peak-normalized ΔF/F_0_ sensor signal increase of at least 0.125), responses to KCl were detected for both NE and 5-HT in n=6/6 experiments (**Figure 2I**). While NE sensor cells were driven robustly by 25 mM KCl, 5-HT sensor cells consistently required a higher dose of KCl (50 mM) to be fully activated (**Figure 2J**). In response to 50 mM KCl, NE reached an average peak concentration of 13.4 µM (**Figures 2K and S4G**). As with the dLGN recordings, we also identified sizable populations of AVP, OXT, and HA sensor cells that were activated by dipping into the corresponding ligands but had no matching masks in the PVH slice recordings (**Figure S4E**). Thus, despite robust prior evidence of signaling at the PVH via the AVP and OXT peptides, we were unable to detect their release in response to KCl stimulation with MORSE^58^. This may be due to recordings at an incorrect depth or location in PVH, release of these molecules at concentrations below the detection limits of the sensors, insufficient quantities of these molecules reaching the probe to drive sensor responses due to their larger molecular weights and slower diffusion through brain tissue, and/or the presence of esterase and peptidase enzymes in the tissue.

Together, these findings demonstrate the ability of MORSE probes to detect endogenous signals from the brain. The results also highlight the viability of these extended, multi-step experiments, as most hydrogels remained stable through over an hour of recording on a slice (in n=27/29 probes), as well as a subsequent 1.5+ hours of post hoc dipping into wells containing ligands (in n=9/11 probes). Furthermore, these multi-sensor probes enabled us to quantitatively track several neuromodulators released from brain tissue simultaneously and unmix crosstalk at sensors variably responsive to multiple ligands. The NE concentration detected at the PVH in response to 50 mM KCl was over six-fold higher than at the dLGN, suggesting that the PVH may receive a greater density of noradrenergic inputs than the dLGN in mouse, consistent with anatomical findings in rats and macaque monkeys^85,86^.

### Real time tracking of CSF composition with MORSE *in vivo*

To begin to test MORSE’s ability to record signals in the awake brain, we focused on the cerebrospinal fluid (CSF). The CSF flows via the brain ventricles and the subarachnoid space, distributing both endogenous signals and pharmaceuticals^9,87–89^. AVP and OXT are two endogenous peptides implicated in a number of important central processes including sociability, feeding, and physiologic regulation^78–84^. Reduced CSF levels of AVP and OXT have been linked with certain symptoms of autism spectrum disorder, including anxiety and social deficits, and intranasal administration of both molecules for brain delivery has undergone trials for improvement of such symptoms^22,90–95^. Both molecules are also commonly delivered peripherally in the clinical setting for blood pressure support and induction of labor during pregnancy, and may require passage into the brain to achieve their full therapeutic effects^96^. However, the central bioavailability of peripherally administered AVP and OXT remains poorly understood. We prepared mice with a headpost and chronic implantation of cannulae in each of the two lateral ventricles (**Figure 3A**). We could observe and collect pristine cerebrospinal fluid from 2-24 weeks following the surgery and also confirmed cannula location histologically (**Figure S5A**). To record from the CSF, we acutely inserted a MORSE probe into one cannula of an awake mouse (both sides used interchangeably across mice). In a subset of experiments, we also introduced a catheter for ICV infusion via the contralateral ventricular cannula. We then performed 3D volumetric two-photon imaging of the probe as previously described while the mouse received sequential intraperitoneal (IP) drug deliveries, followed by intracerebroventricular (ICV) delivery through the second cannula when indicated per experiment (**Figure 3B; Supplementary Video 9**). Post hoc *ex vivo* dipping characterization of each probe was performed to identify sensor cell subpopulations (**Supplementary Video 10**).

**Figure 3.**
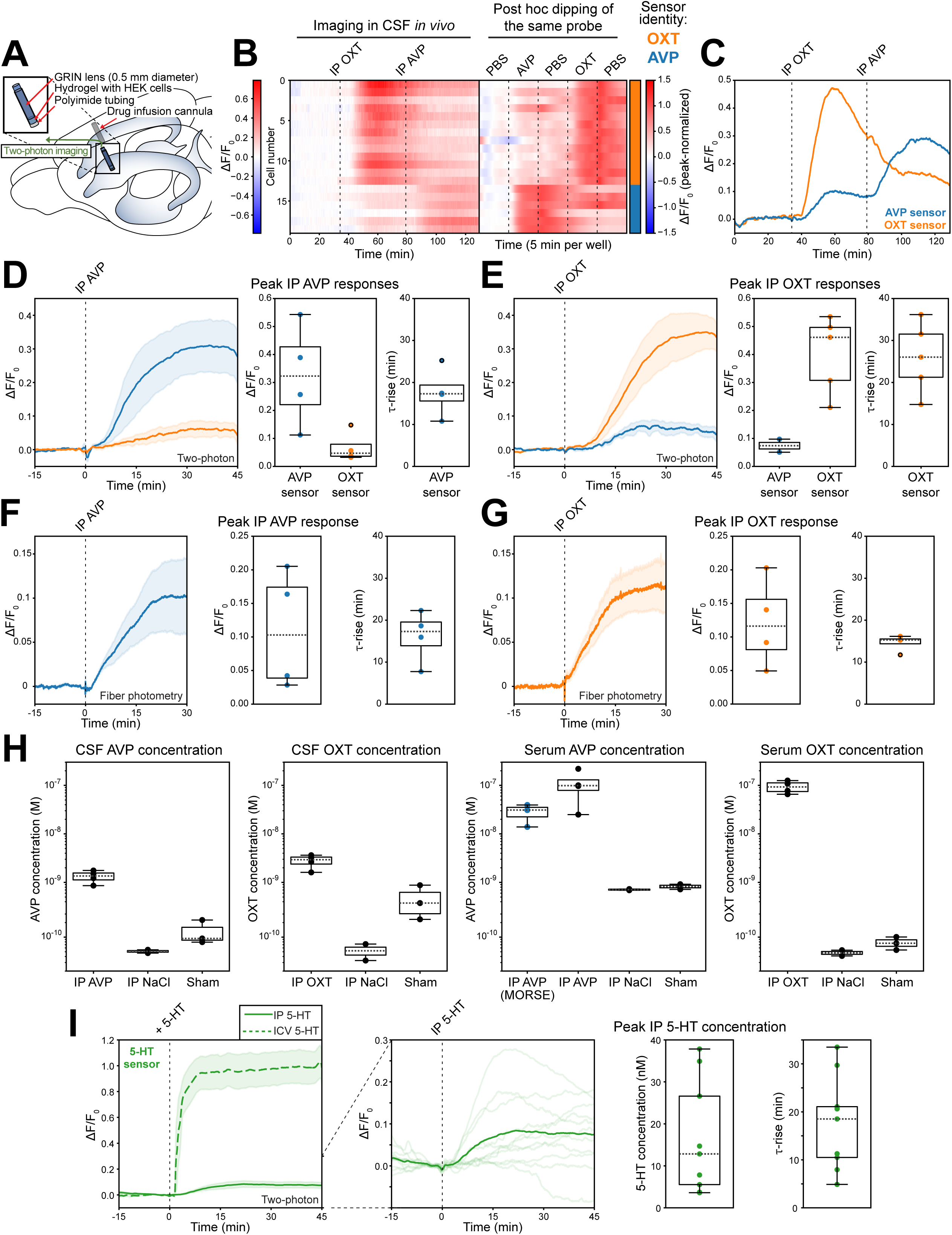
Three-sensor *in vivo* MORSE recordings in the lateral ventricle CSF of awake mice. (**A**) Schematic of the mouse lateral ventricle cannulation setup, demonstrating two chronically implanted cannulae at the anterior aspects of the lateral ventricles. Before a recording, the mouse was head-fixed on a running wheel and one of the cannulae was acutely opened to expose the CSF, at which point the MORSE probe was inserted hydrogel-first into the cannula and sealed into place with RTV silicone. Following several hours of imaging, the probe was removed from the animal for post hoc dipping characterization. (**B**) Single-sensor cell traces from one three-sensor MORSE probe (AVP, OXT, 5-HT – not shown) recorded in the mouse CSF (left heatmap, ΔF/F_0_ depicted as per left color bar) and subsequently dipped into standard solutions (right heatmap, peak-normalized ΔF/F_0_ depicted as per right color bar). OXT and AVP were delivered IP in 200 μL of PBS at a dose of 1 mg/kg at the indicated times. During dipping, the probe was dipped for 5 minutes at a time into 20 µL volumes of standard solutions of the three ligands corresponding to each sensor formulated in PBS (1 μM AVP, 1 μM OXT, 1 μM 5-HT – not shown), with interposed dips into wash wells containing PBS between them. The demonstrated sensor cells had their ROIs successfully matched between the *in vivo* recording and the dipping characterization. The sensor identity of each cell is shown with the color bar on the right as per legend. (**C**) Mean ΔF/F_0_ traces for each sensor type from the left heatmap in **B**, with IP injection times indicated and sensors color-coded as per legend. (**D**) Three-sensor (AVP, OXT, 5-HT) MORSE probes were recorded in mouse CSF during IP AVP administration. **Left:** mean ΔF/F_0_ responses in AVP sensor cells (n=4 mice) and OXT sensor cells (n=4 mice) in mouse CSF following IP injection of AVP. Shading depicts SEM across mice. **Middle:** peak ΔF/F_0_ signals attained in each recording summarized in **D, left**. **Right:** summary of 1-rise onset timings of the AVP sensor time courses summarized in **D, left**. (**E**) Three-sensor (AVP, OXT, 5-HT) MORSE probes were recorded in mouse CSF during IP OXT administration. **Left:** mean ΔF/F_0_ responses in AVP sensor cells (n=2 mice) and OXT sensor cells (n=5 mice) in mouse CSF following IP injection of OXT. Shading depicts SEM across mice. **Middle:** peak ΔF/F_0_ signals attained in each recording summarized in **E, left**. **Right:** summary of 1-rise onset timings of the OXT sensor time courses summarized in **E, left**. (**F**) Fiber photometry probes carrying cultured cells expressing AVP sensor only were recorded in mouse CSF during IP AVP administration, as shown in **Fig. S5A**. **Left:** mean ΔF/F_0_ responses from AVP sensor fiber photometry probes (n=4 mice) in mouse CSF following IP injection of AVP. Shading depicts SEM across mice. **Middle:** peak ΔF/F_0_ signals attained in each recording summarized in **F, left**. **Right:** summary of 1-rise onset timings of the AVP sensor photometry time courses summarized in **F, left**. (**G**) Fiber photometry probes carrying cultured cells expressing OXT sensor only were recorded in mouse CSF during IP OXT administration, as shown in **Fig. S5A**. **Left:** mean ΔF/F_0_ responses from OXT sensor fiber photometry probes (n=4 mice) in mouse CSF following IP injection of OXT. Shading depicts SEM across mice. **Middle:** peak ΔF/F_0_ signals attained in each recording summarized in **G, left**. **Right:** summary of 1-rise onset timings of the OXT sensor photometry time courses summarized in **G, left**. (**H**) ELISA quantifications of AVP and OXT levels in CSF and serum collected from naïve mice 30 minutes following IP injections of either AVP or OXT (1 mg/kg), saline controls, or sham injections, as shown in **Fig. S5B**. Each black dot represents an ELISA reading from a single mouse, with one CSF and one serum reading from each of n=18 animals. In the case of serum following IP AVP administration, dipping of a MORSE probe *in vitro* into the samples was additionally used to quantify AVP concentration in the serum. Results from n=3 such mice are plotted with blue dots, and raw ΔF/F_0_ plots are shown in **Fig. S5C**. Neither AVP nor OXT sensor activation was detected in CSF following IP administration of either ligand using MORSE dipping *in vitro*. OXT sensor activation was detected in serum following IP administration using MORSE dipping *in vitro*, but OXT concentrations in these samples could not be estimated from the collected ΔF/F_0_ signal due to incomplete post hoc dose-response calibration. (**I**) Ten-sensor (AVP, OXT, 5-HT, CCK, NE, DA, ACh, ATP, MT, HA) MORSE probes were recorded in mouse CSF during IP and ICV 5-HT administration. **Far left:** mean ΔF/F_0_ responses in 5-HT sensor cells in mouse CSF following ICV (5 nmol/µL in aCSF infused at 5 µL/min over 6 min, n=3 animals, 1 injection per animal) and IP (2 mg/kg, n=5 animals, 1-3 injections per animal for a total of n=11 injections) deliveries of 5-HT. Shading depicts SEM across injections of each type. **Center left:** ΔF/F_0_ responses in 5-HT sensor cells in mouse CSF during each of n=11 IP injections of 5-HT. Solid line depicts the mean response across all injections. **Center right:** quantification of the peak CSF 5-HT concentrations attained in the n=9/11 (from n=3 mice) IP injections of 5-HT demonstrated in **I, center left** for which post hoc dose-response calibration was completed. **Far right:** quantification of 1-rise onset timings of the n=9/11 IP injections of 5-HT demonstrated in **I, center left** for which post hoc dose-response calibration was completed.

We initially delivered AVP or OXT by IP injection (1 mg/kg, n=4 and n=5 experiments, respectively) and reliably observed a corresponding sensor response in the CSF within at most 10-15 minutes, followed by washout (**Figures 3B-C**). In a subset of these experiments (n=6/9 total), AVP and OXT were delivered sequentially in the same recording and each delivery activated different cells on the same probe with distinct, independent response time courses showing good sensor specificity (**Figures 3D-E**). To assess these molecules one at a time in CSF with a simpler fiber photometry setup, we replicated the analogous probe design on an optical fiber, utilizing one GRAB sensor at a time as reported recently (**Figure S5B**)^97^. This approach demonstrated similar kinetics of CSF AVP and OXT following IP injection as using the MORSE approach involving two-photon imaging (**Figures 3F-G**).

To confirm the physiological relevance of these recordings, we also performed analogous IP injections in naïve, unoperated mice, collected serum and CSF 30 minutes later, and performed ELISA quantification of AVP and OXT concentrations (**Figures 3H and S5C**). We found that IP injections of AVP and OXT led to elevated levels in both serum and CSF compared to IP saline control injections, with levels in CSF reaching the single nanomolar range. Notably, when we performed MORSE two-photon dipping experiments in these fluids, we could detect responses only from the serum but not from the CSF, likely due to ligand degradation between collection and *ex vivo* dipping, possibly via actions of endogenous peptidase enzymes (**Figures 3H and S5D-E**). These findings suggest that MORSE probes retain their ligand sensitivity and specificity over >4 hours of recording in the CSF of a live mouse, with sufficient probe stability to allow successful subsequent *ex vivo* dipping across several hours. Further, these probes can track pharmacologically relevant concentration changes in CSF over time while avoiding the risks of removing the fluid from the body, such as contamination during collection and rapid analyte breakdown.

To assess if we could multiplex more sensors simultaneously with MORSE *in vivo*, we further extended the CSF recordings to probes containing the full set of ten sensors used in **Figure 1E**. We sequentially delivered three IP injections of mixtures of four ligands at a time, with 5-HT present in every mixture, followed by a final saturating ICV injection of 5-HT alone (chosen to ensure repeatability and specificity of calibration at the 5-HT sensor cells). Across n=3 experiments, we were able to successfully recover signals from up to 9 sensors at the same time (median of 8 active sensors per probe, average of 5.1 cells per sensor per probe), with each sensor demonstrating distinct CSF time courses consistent with the IP injection time of its respective ligand (**Figure S5F**). In the case of 5-HT, we also used the ICV response in each recording to normalize the responses from the sensor and estimate quantitative time courses of CSF 5-HT concentration, demonstrating an average peak concentration of 16.4 nM in response to IP delivery (**Figure 3I**). This demonstrates how MORSE can facilitate *in vivo* studies of CSF pharmacodynamic interplay in combinatorial drug regimens, with a temporal resolution of seconds and the potential for quantitative concentration readouts.

### MORSE detects endogenous CSF signals at concentrations that impact downstream signaling

The MORSE probes we used in our IP AVP and OXT experiments always carried the 5-HT sensor as well. In a subset of these experiments (n=3/7), we observed a spontaneous elevation in CSF 5-HT that was evident in all 5-HT sensor cells (but none of the other sensor cells) (**Figures 4A-B**). These fluctuations were of similar magnitude or larger than the signals we observed in CSF following IP 5-HT delivery (c.f. **Figure 3I**) and slowly rose and decayed over tens of minutes (**Figures 4C-D**). Fluctuations in CSF 5-HT on a timescale of several hours have been described in humans, but their mechanisms and impacts are unknown^98^. One of the tissues that prominently contacts the CSF is the ChP epithelium, a sheet of cells within each ventricle that modulates CSF composition. The ChP epithelium is known to express high levels of 5-HT2C receptor, a serotonin G_αq_-coupled GPCR which drives an increase in cytosolic calcium and in secretory activity in these epithelial cells^99^. This receptor appears to be enriched on the highly elaborate microvillous apical surface of the ChP epithelium, which directly contacts the CSF in the ventricle^100^.

**Figure 4.**
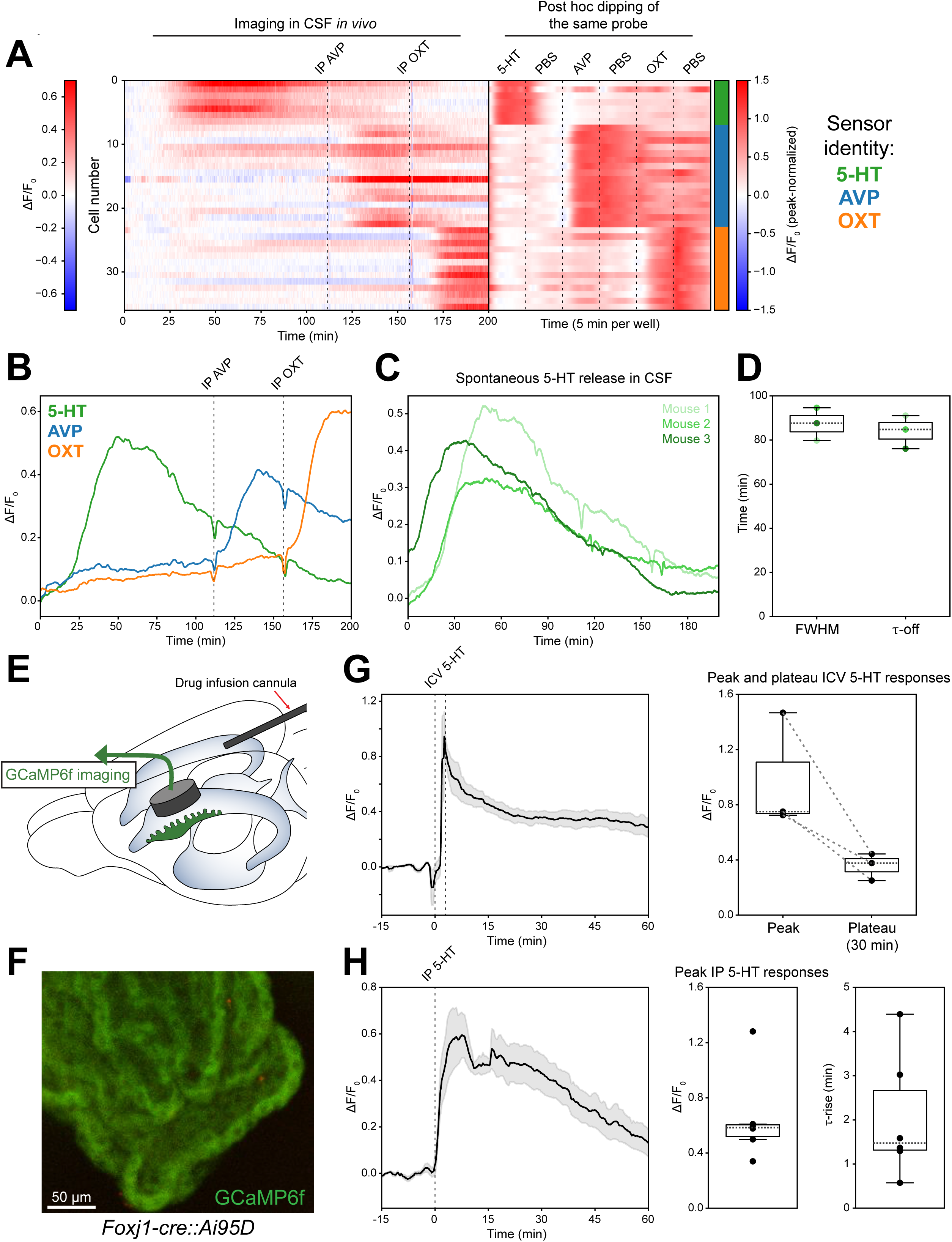
Spontaneous CSF 5-HT fluctuations identified by three-sensor *in vivo* MORSE recordings in the lateral ventricle CSF of awake mice. In a subset of n=3/7 three-sensor MORSE recordings summarized in Fig. 3A**-E**, what appear to be spontaneous CSF 5-HT fluctuations were identified. (**A**) Single-sensor cell traces from one three-sensor (AVP, OXT, 5-HT) MORSE probe recorded in the mouse CSF (left heatmap, ΔF/F_0_ depicted as per left color bar) and subsequently dipped into standard solutions (right heatmap, peak-normalized ΔF/F_0_ depicted as per right color bar). AVP and OXT were delivered IP in 200 μL of PBS at a dose of 1 mg/kg at the indicated times. During dipping, the probe was dipped into standard solutions of the three ligands corresponding to each sensor formulated in PBS (1 μM AVP, 1 μM OXT, 1 μM 5-HT), with interposed dips into wash wells containing PBS between them. The demonstrated sensor cells had their ROIs successfully matched between the *in vivo* recording and the dipping characterization. The sensor identity of each cell is shown with the color bar on the right as per legend. Spontaneous activation of 5-HT sensor is evident in the first 100 minutes of the recording. (**B**) Mean ΔF/F_0_ traces for each sensor type from the left heatmap in **A**, with IP injection times indicated and sensors color-coded as per legend in **A**. (**C**) Summary ΔF/F_0_ traces from 5-HT sensor cells in n=3 three-sensor MORSE recordings demonstrating spontaneous 5-HT fluctuations. (**D**) Quantification of dynamic parameters of the spontaneous 5-HT signals summarized in **C**. Full width at half maximum (FWHM) and 1-off offset timings following achievement of peak signal are shown. Color hues of the plotted dots match the hues of the corresponding traces in **C**. (**E**) Schematic of the mouse ChP epithelium GCaMP6f imaging setup, demonstrating a 3 mm diameter cranial window with a sealed circular glass front face chronically implanted over the large leaflet of the lateral ventricle choroid plexus within the lateral ventricle of a *FoxJ1-Cre::Ai95D* mouse, as well as a drug infusion cannula chronically implanted in the contralateral ventricle from the posterior occipital aspect. 3D two-photon imaging of the ChP epithelial cells was performed via the cranial window, with the contralateral cannula optionally used for ICV infusions during the recording. (**F**) Example still image of a ChP epithelium in an awake *FoxJ1-Cre::Ai95D* mouse imaged through the cranial window as depicted in **E**. Green fluorescence of GCaMP6f can be seen filling the cytosol of the epithelial cells. Scale bar: 50 µm (**G**) **Left:** mean ΔF/F_0_ GCaMP6f response to ICV 5-HT delivery (5 nmol/µL in aCSF infused at 5 µL/min over 3 min, n=3 mice) across representative areas of ChP epithelium in *FoxJ1-Cre::Ai95D* mice. **Right:** comparison of peak and plateau (30 min timepoint) ΔF/F_0_ GCaMP6f responses in the n=3 mice summarized in **G, left**. Values from the same recording are connected by dotted lines. (**H**) **Left:** mean ΔF/F_0_ GCaMP6f response to IP 5-HT delivery (2 mg/kg, n=6 mice) across representative areas of ChP epithelium in *FoxJ1-Cre::Ai95D* mice. **Middle:** peak ΔF/F_0_ signals attained in each recording summarized in **H, left**. **Right:** summary of 1-rise onset timings of the GCaMP6f time courses summarized in **H, left**.

We wondered whether CSF 5-HT concentrations of the magnitudes observed during spontaneous fluctuations and following IP injection might exert a meaningful impact on the adjacent ChP. To this end, we prepared mice expressing the cytosolic calcium indicator GCaMP6f in the ChP epithelial cells (in *FoxJ1-Cre::Ai95D* mice)^100^. In adult animals, we implanted an imaging cannula with a glass window into one lateral cerebral ventricle and an infusion cannula into the other (**Figures 4E-F**)^89,100,101^. After 4 weeks of recovery and habituation to head restraint, awake mice received either ICV (75-150 nmol or ∼0.5-1 mM) or IP (3 mg/kg) deliveries of 5-HT during 3D two-photon GCaMP imaging of the ChP epithelium (**Figures 4G-H; Supplementary Videos 11-12**). Both routes of 5-HT administration drove similar amplitudes of calcium activity in the ChP epithelial cells, and both responses persisted for at least 60 minutes following delivery. This confirmed that changes in either central or peripheral 5-HT can drive second messenger activity in the ChP epithelium, likely resulting in downstream effects on its secretory activity^100,101^. Notably, plasma 5-HT concentrations are known to exhibit substantial elevations in particular states such as viral illness, which may influence CSF composition through its action at the ChP^102^.

Molecular transport through the CSF is tightly regulated, and the CSF components that we recorded with MORSE must reach receptors at the ChP epithelium, other CSF-facing barriers, or the brain parenchyma in order to drive downstream signaling. Using AAV5 viral vectors, which exhibit a tropism for ChP epithelial cells in mice^103^, we expressed GRAB sensors directly in the ChP to confirm that MORSE recordings estimate CSF signals in a manner that reflects their levels and dynamics immediately adjacent to this tissue. We began by delivering viruses expressing the same AVP, OXT, and 5-HT sensors as used above (GRAB-AVP0.2, OXT1.7, 5-HT2h) to mouse CSF by ICV injection. One week later, we implanted the same lateral ventricle imaging windows and infusion cannulae as above (see **Figure 4E**). Subsequent two-photon imaging in awake animals weeks later demonstrated successful expression of each of the sensors on the membranes of the ChP epithelial cells (**Figure 5; Supplementary Video 13**). We additionally injected a Texas Red-labeled 70 kDa dextran in these mice IP to label the vasculature and confirmed GRAB sensor expression on the epithelial surfaces facing away from the vessels and towards the CSF. Notably, ChP epithelial cells also possess a basal membrane that faces a stromal space surrounding a fenestrated capillary bed, but the apical membrane makes up the vast majority of the plasma membrane area in these cells due to its highly elaborate microvilli^104^.

**Figure 5.**
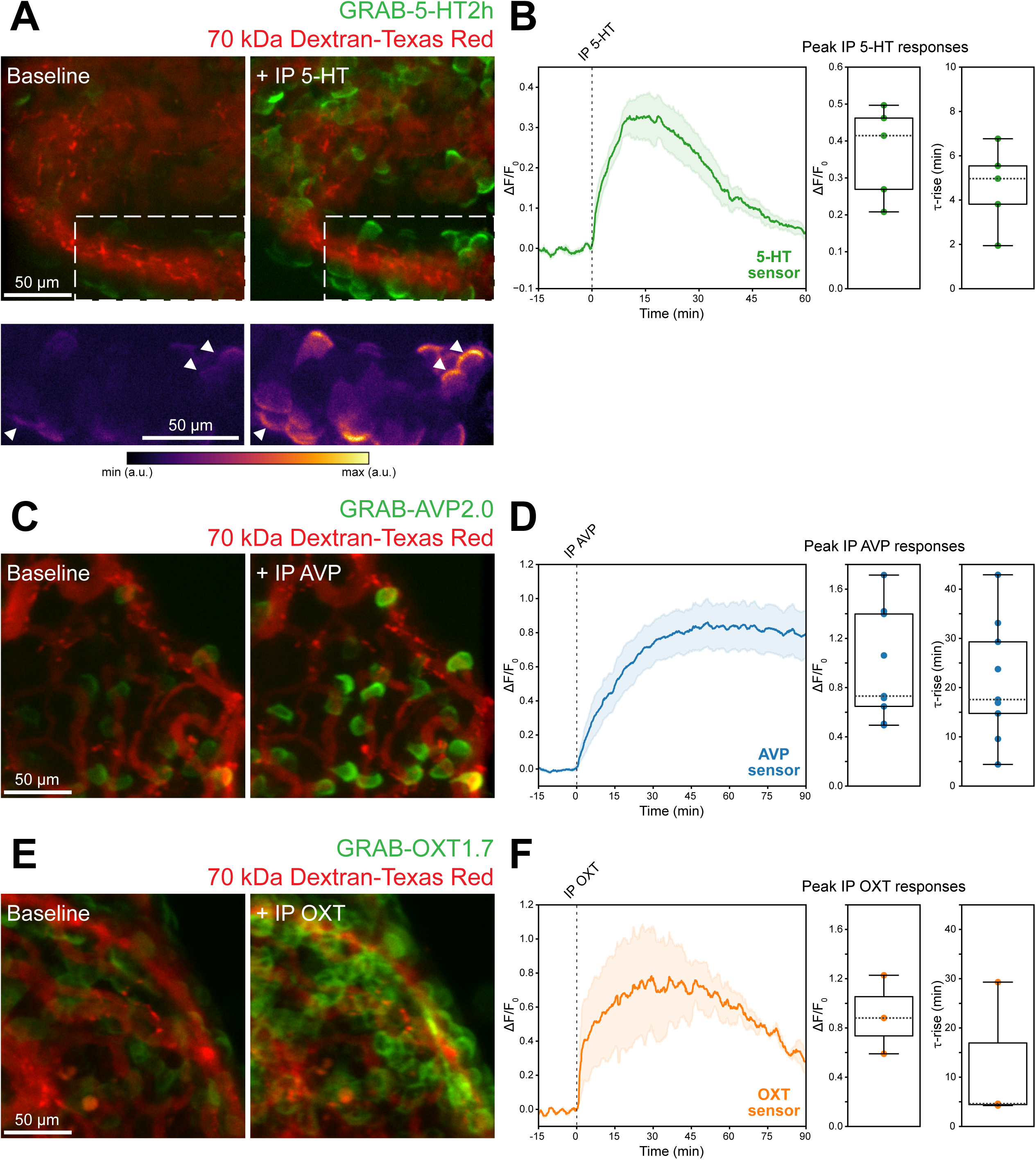
GRAB sensors expressed in the choroid plexus epithelium directly report chemical signals in awake mouse CSF. (**A**) **Top:** choroid plexus imaging in a mouse infected with AAV2/5 carrying GRAB-5-HT2h demonstrates robust expression of the sensor along the apical membranes of the epithelial cells facing away from the vasculature labeled with 70 kDa Dextran-Texas Red. Upon IP delivery of 5-HT (2 mg/kg), there is a robust activation of the sensor on the membranes of the epithelial cells. **Bottom:** closeup of the dashed area in the upper images, with the green GRAB-5-HT2h channel shown in false color. White arrowheads indicate sensor activation on the apical epithelial membranes facing the CSF. Scale bars: 50 μm. (**B**) **Left:** mean ΔF/F_0_ GRAB-5-HT2h response to IP 5-HT delivery (2 mg/kg, n=5 injections across n=4 mice) across representative areas of ChP epithelium expressing GRAB-5-HT2h. **Middle:** peak ΔF/F_0_ signals attained in each recording summarized in **B, left**. **Right:** summary of τ-rise onset timings of the GRAB-5-HT2h time courses summarized in **B, left**. (**C**) Choroid plexus imaging in a mouse infected with AAV2/5 carrying GRAB-AVP0.2 demonstrates robust expression of the sensor along the apical membranes of the epithelial cells. Upon IP delivery of AVP (1 mg/kg), there is a robust activation of the sensor on the membranes of the epithelial cells facing the CSF. Scale bar: 50 μm. (**D**) **Left:** mean ΔF/F_0_ GRAB-AVP0.2 response to IP AVP delivery (1 mg/kg, n=9 injections across n=4 mice) across representative areas of ChP epithelium expressing GRAB-AVP0.2. **Middle:** peak ΔF/F_0_ signals attained in each recording summarized in **D, left**. **Right:** summary of τ-rise onset timings of the GRAB-AVP0.2 time courses summarized in **D, left**. (**E**) Choroid plexus imaging in a mouse infected with AAV2/5 carrying GRAB-OXT1.7 demonstrates robust expression of the sensor along the apical membranes of the epithelial cells. Upon IP delivery of OXT (1 mg/kg), there is a robust activation of the sensor on the membranes of the epithelial cells facing the CSF. Scale bar: 50 μm. (**F**) **Left:** mean ΔF/F_0_ GRAB-OXT1.7 response to IP OXT delivery (1 mg/kg, n=3 injections across n=2 mice) across representative areas of ChP epithelium expressing GRAB-OXT1.7. **Middle:** peak ΔF/F_0_ signals attained in each recording summarized in **F, left**. **Right:** summary of τ-on onset timings of the GRAB-OXT1.7 time courses summarized in **F, left**.

IP deliveries of 5-HT, AVP, and OXT in separate experiments involving each associated sensor drove sensor signals on the apical membranes of the ChP epithelial cells with similar time courses as observed previously with MORSE (compare **Figures 5B** and **3I, 5D and 3D, 5F and 3E**). These experiments demonstrate a ChP imaging method for gold-standard, spatially resolved tracking of one signal in the CSF at a time. Furthermore, IP 5-HT reached the apical ChP membrane fast enough to explain the calcium sensor response it drove in the tissue (compare **Figures 5B** and **4H**). This suggests that CSF levels of 5-HT following IP delivery or during endogenous fluctuations were of sufficient magnitude to bind GPCRs at the ChP and drive calcium responses. Taken together, these experiments demonstrate that MORSE is a promising platform for high-dimensional tracking of the dynamic composition of brain fluids, with sufficient sensitivity to detect signals of likely biological relevance.

## Discussion

### Open questions and challenges in tracking extracellular chemical signals

The field of neuroscience has made tremendous strides in recording increasingly nuanced electrical signals from thousands of neurons simultaneously. In the meantime, the detection of neurotransmitters and neuromodulators – the molecules that brain cells use to communicate with each other – has lagged behind due to critical technical challenges. The state of the art in brain fluid composition studies still involves fluid samples collected at discrete times. In rodents, these studies are usually terminal, often require pooling of samples to overcome detection limits and minimum volume requirements for standard ELISA and mass spectrometry-based approaches, and likely involve analyte breakdown prior to *ex vivo* analysis. Developing new strategies for high-dimensional recordings of fluid composition is important: while pharmacological studies typically examine the effects of one ligand at a time at supraphysiologic doses, every neuron is sensitive to tens or possibly hundreds of signaling molecules that are dynamically changing in concentration and affecting its activity^11–15^. To better understand how these fluctuations in chemical signals shape brain states, we need tools capable of scalable tracking of many neuroactive molecules simultaneously and in real time.

Recent developments in sensors based on endogenous receptors, like the GRAB sensors, have achieved high signal-to-noise tracking of biologically relevant concentrations of many ligands recorded one at a time. Design principles gleaned from developing the first generation of GPCR-based sensors appear to be sufficiently general to allow efficient development of sensors for many new molecules over the coming decade. Beyond the challenge of producing sensors, a key hurdle involves the integration of many sensors into one probe. Successful scalable sensing platforms, such as multielectrode arrays commonly used in neuroscience, use a single detector architecture and modality in a massively parallelized fashion^105^. Yet, there are few examples of scalable biosensor probes due to the diversity of sensor strategies. One of the few previous attempts to read out many optical sensors simultaneously, MOSAIC, succeeded by printing viral particles carrying individual genetically encoded sensor constructs onto a flat substrate in islands spaced across an area of ∼10 mm^2^ ^106^. When cells were subsequently cultured on the substrate, they demonstrated spatially segregated expression patterns of the sensors. However, this approach was not designed for *in vivo* applications and the spatial footprint was too large to meaningfully capture regionalized signaling in the brain.

### Novel achievements and caveats of MORSE

To our knowledge, MORSE is the first demonstration of a probe that is both (i) capable of recording >10 optical sensors simultaneously and (ii) sufficiently compact (0.5 mm diameter, 0.2 mm^2^ sensing area) for *in vivo* chemical tracking in the brain of an awake, behaving animal. Key to this technology was the development and optimization of a thin and stable yet porous hydrogel interface that could adhere several hundred cultured cells to the front of a GRIN lens for many hours. Our approach is scalable due to the large number (>100) of sensor cells that we can image in 3D together with the ability to pair experiments with robotic dipping of the same probe into many standard solutions. This allows for demultiplexing of sensor signals based on 3D location of each sensor cell in the hydrogel and sets the stage for quantitative estimation of absolute signal concentrations in a given brain region. Importantly, simultaneous recording of sensors exhibiting ligand crosstalk allows for straightforward unmixing of these effects to determine individual signal dynamics. We expect that the number of signaling molecules that can be studied in this fashion will substantially expand as the repertoire of genetically encoded sensors continues to grow. This approach is also extendable to other cell-based sensors such as CNiFERs as well as cell-free soluble sensors immobilized on microbeads or other carriers^107,108^. Despite robust calibration and normalization, signal estimates remain susceptible to the usual downsides of intensity-based sensors, such as photobleaching. However, recent studies suggest that some GRAB sensors exhibit measurable changes in fluorescence lifetime upon ligand binding, which could be recorded in the current probe configuration and would allow quantitative tracking of absolute concentrations across days^47,109^. Furthermore, MORSE is compatible with real-time signal extraction from two-photon imaging, which would help enable high-dimensional closed-loop control of chemical environments^110–112^.

Our design possesses several limitations that could be improved in the future. First, we rely on a hydrogel to immobilize the sensor-expressing cells. A similar recent probe design, FOPECS, recorded a single GRAB sensor at a time *in vivo* using fiber photometry^97^. We optimized our fragile hydrogel significantly and combined several surface coatings at the lens-hydrogel interface to achieve reliable *in vivo* stability of ∼10-fold smaller gel volumes, which enhances sensor sensitivity and time resolution. Nevertheless, further improvements to probe sensitivity and speed will be useful. Another challenge with our approach is the reliance on cultured cells to express the sensors. We chose these because GRAB sensors, like many other receptor-based sensors, are transmembrane proteins that have been optimized and validated in cultured cells. While we did our best to impede cell motility and reduce immune response by treating animals with dexamethasone, future strategies could replace the use of cultured cells with sensor expression in liposomes or lipid bicelles^113,114^. These could potentially be purified and patterned directly on the lens face, resulting in a more stable probe-tissue interface that is essentially 2D^115^. Reducing immunogenic epitopes on the probes in this way, as well as potentially mixing immunosuppressant drugs directly into the hydrogel, would likely improve permeability at the probe-brain interface and increase the likelihood of their applicability in brain tissue in animal models and humans. While we did not assess the *in vivo* stability of our probes beyond several hours of imaging, previous hydrogel-based probes with similar recipes have been shown to remain functional for several days, though image quality may be compromised if the gel deforms or deteriorates over that time^97^.

### Implications and next steps

We chose the CSF as the proof of principle brain fluid to study with MORSE *in vivo* because it is a readily accessible bulk fluid with important functions in brain-wide signaling and a major conduit for pharmaceuticals intended to reach the brain. CSF composition and dynamics remain understudied due to technical limitations, with recent developments limited to fiber photometry tracking of transgenic or exogenous fluorescent proteins or tracers examined one at a time^2,116^. As such, general principles underlying the central trafficking of peripheral signals remain poorly understood. Remarkably, we found that most if not all of the ligands we delivered reached detectable concentrations at MORSE probes in CSF within 15 minutes of an IP injection at common doses used in the literature, suggesting that most molecules of sufficiently small molecular weight may cross from blood to CSF at appreciable quantities^89^. This is at odds with some previous reports suggesting that some of the molecules we tested – particularly the AVP and OXT peptides – “do not readily cross the blood-brain barrier^117,118^,” and underscores the role of the CSF in biodistribution of global signals to CSF-adjacent tissues (ChP and periventricular ependymal cells) and throughout the brain (though it is important to note that CSF and brain parenchymal concentrations are likely substantially decoupled). Our findings also suggest an important new avenue for pharmaceutical testing. In addition to AVP and OXT, the clinical relevance of which was described previously, other potent molecules that we tested include dopamine and melatonin, which have longstanding therapeutic uses in Parkinson’s disease and various sleep disorders, respectively.

We additionally detected previously unstudied spontaneous fluctuations in CSF 5-HT and demonstrated that they are likely sufficient to drive intracellular calcium activity in the apposing ChP epithelial cells, suggesting that the concentrations we observed are biologically meaningful and that at least certain CSF-contacting tissues are readily poised to respond to these signals^98^. This expands on previous studies that have established a role for the CSF in distributing signaling molecules mediating physiological transitions across circadian and diurnal rhythms, hunger and satiety, stress, and learning^2,5,119–121^. Future studies are needed to better understand how neurons sense these components in the CSF, either by sending projections directly into the fluid or by responding to molecules that have diffused into the interstitial fluid near the ventricle walls^122,123^. It also remains to be seen how all the barrier structures between blood and CSF – including the ChP as well as the circumventricular organs, meninges, blood-brain barrier of the neurovascular endothelium and others – work together to shape the bioavailability of each signaling factor^124^.

Multiplexed ligand tracking with MORSE also showed promise as a potential tool to study high-dimensional dynamics of interstitial fluid composition in the brain parenchyma. As we demonstrated in our brain slice experiments, the probe can detect evoked norepinephrine and serotonin release from brain slices in a quantitative manner that likely reflects the density of local neuromodulatory innervation. While previous studies using fast-scan cyclic voltammetry have performed similar quantitative measurements of monoamine release, they could not record all molecules simultaneously and did not allow for catecholamines NE and DA to be directly distinguished^125^. We were unable to detect evoked peptide release from AVP and OXT neurons in the PVH, likely in part because we were unable to position the sensors close enough to the sites of release. This effective distance can be improved in future versions of the probe by using a thinner hydrogel or shifting to 2D coating strategies as described above. We speculate that our ability to detect monoaminergic but not peptidergic signaling is not due to differences in sensor sensitivity but rather a reflection of differences in molecular size, which has a profound influence on diffusion speed and effective signaling distance in brain parenchyma.

Looking to the future, since there is already a precedent of successfully recording norepinephrine GRAB sensor signals in brain tissue using a hydrogel-based fiber photometry probe^97^, we expect that our slice findings will be replicable in brain parenchyma *in vivo*, at least for the monoamines and other small molecules. An essential yet unaddressed challenge in neuroscience is that these signals are often partially correlated but studied one at a time, even though various brain cell types integrate many of these signals simultaneously. Understanding the *dynamic patterns* of neuromodulatory concentrations across brain states is a new frontier, one that is critical to understanding how individual cells sense and adapt to changing brain states. Furthermore, by performing volumetric imaging through the GRIN lens, we should in principle be able to image beyond the ∼200 µm sensor layer and concurrently record the activity of nearby brain cells expressing genetically encoded sensors. This approach would allow for the combinatorial influences on neural activity of diffusible signals and of local connectivity to be disentangled in greater detail. Cannula and probe implantation in tissue does pose additional challenges in terms of inflammation followed by gliosis and scarring, which may restrict ligand diffusion beyond what we found in acute slices. Additional innovation in miniaturizing the MORSE probes and reducing their immunogenicity should further enhance the viability of this approach for diverse applications in brain tissue *in vivo*. Ultimately, these efforts will improve our understanding of how brain fluids change in composition across a range of timescales in a quantitative manner and allow for novel assessments of how sets of molecular signals act in concert to encode brain states and concurrently orchestrate the function of diverse populations of brain cells.

## Materials and methods

Experiments were performed in accordance with National Institutes of Health guidelines and were approved by the Institutional Animal Care and Use Committees and Institutional Biosafety Committees at Beth Israel Deaconess Medical Center and Boston Children’s Hospital. Male and female adult mice, 2-12 months old, were used in this study.

### MORSE manufacture

#### Cell culture

Human Embryonic Kidney-293T (HEK-293T) cells (ATCC) and Luciferase/tdTomato Dual-Reporter HEK-293 cells (Alstem) were propagated at 37°C, 95% humidity, and 5% CO_2_ in Denville 100 mm cell culture dishes (Thomas Scientific) stored in a standard tissue culture incubator. Cells were cultured in DMEM medium (high glucose GlutaMAX™; Thermo Fisher Scientific) supplemented with 10% fetal bovine serum (FBS; R&D Systems) and 1% penicillin/streptomycin (P/S, Sigma). Dishes were passaged after reaching 80% confluence (approximately every 5 days), and media was replaced every 2-3 days. After being thawed, cells used for experiments were maintained in culture for up to 8 passages (∼1.5 months) before being discarded.

#### Transfection

HEK-293T cells at ∼80% confluence were detached from the culture dish using mechanical force, resuspended at a density of 300,000-500,000 cells/mL, and seeded in each well of a poly-D-lysine-treated (Sigma) 24-well plate (Corning), either directly on the plate surface or on a 15 mm glass coverslip (Warner Instrument Corp), at 150,000-170,000 cells/well. Each well was transfected with a plasmid encoding a different G protein-coupled Receptor Activation-Based (GRAB) sensor (pDisplay-GRAB-ires-mCherry-CAAX, gifts from Yulong Li) using a standard EFFECTENE® transfection kit (Qiagen). The transfection mixture was optimized separately for each plasmid according to the instructions provided by the supplier. In the end, an EFFECTENE®/DNA ratio (in µL/µg) of 25 was used for NPY1.0, HA1m, AVP0.2, and CCK0.5, while one of 6.25 was used for DA2.9, NE2h, gACh4l, 5-HT2h, OXT1.7, ATP1.0-L, eCB2.0, VIP1.0, MT1.0, Ado2h, PTH1.0, and SST1.0. In a subset of experiments, we also transfected Luciferase/tdTomato Dual-Reporter HEK-293 cells with NE2h. All cells were incubated in the transfection mixture for 24 hours prior to direct imaging or MORSE probe preparation.

#### Cleaning and gluing of GRIN lenses and photometry fibers into tubing

To prepare a MORSE probe, a clean Gradient Refractive Index (GRIN) lens (singlet, 8.85 mm length, 500 μm diameter, GRINTECH NEM-050-25-10-860-S-2.0p) was carefully inserted into an 8 mm long piece of polyimide tubing (PI, internal diameter 0.56 mm, code 220-I, MicroLumen). All lenses were prepared on a hemocytometer under a dissection microscope to allow for exact measurements of probe dimensions (cat no. 3715211/3715298, Carl Zeiss Microscopy). Once the lens was at least 2 mm into the tubing, an applicator was used to add a drop of Kwik-Sil (World Precision Instruments) – an optically clear RTV silicone elastomer – to the lens-tubing interface. The tubing was then slowly moved along the lens, allowing the Kwik-Sil to gradually seep around the lens shaft until the tubing extended 0.3 mm beyond the front of the lens. The protruding tubing provided a small receptacle with a volume of ∼0.2 nL in front of the lens face surrounded by tubing. Lenses were then laid flat, and the Kwik-Sil was allowed to harden at room temperature for at least 1 hour. Once the Kwik-Sil had hardened, two lines were drawn around the circumference of the PI tubing, 7 mm from the end of the receptacle; this would ensure the probe was inserted into a mouse’s guide cannula (see below for details) to the correct depth. MORSE photometry probes were prepared in an analogous manner using 6.5 mm photometry fibers (400 µm diameter, Doric MFC_400/470-0.37_6.5mm_MR1.25_FLT) and 6 mm pieces of PI tubing (internal diameter 0.48 mm; code 190-I; MicroLumen).

We also developed a method to clean and re-use the GRIN lenses and optical fibers between successive experiments. First, probes were left in 100% acetone (VWR) for at least 24 hours. To begin to remove the PI tubing, Cohan-Vannas spring scissors (item No. 15000-02, Fine Science Tools) were used to cut the PI tubing along the length of the probe. The GRIN lens or fiber was then removed from the cut tubing using ceramic forceps (11 cm, cat no. 501209; World Precision Instruments) and dipped into a bath of 70% ethanol (Fisher). An applicator brush (cat no. S379; Parkell) dipped in ethanol was used to scrub the remaining Kwik-Sil from the sides of the lens or fiber. Once the sides were clean, the lens or fiber was mounted with one face up on a stainless steel micro V-clamp (cat. VK250, Thorlabs) under a dissection microscope. A thin piece of lens paper (Fisherbrand) dipped in ethanol was used to clear debris from each lens or fiber face. GRIN lenses and photometry fibers were reused and reimaged in this fashion up to 10+ times and retired when they accumulated excessive scrapes and blemishes.

#### Pre-coating of GRIN and photometry fiber hydrogel receptacles

After being cleaned and glued into PI tubing, GRIN lenses were placed in a polystyrene foam holder with the receptacle face-up and cooled to 4°C in a cold room. To fill any air gaps remaining between the PI tubing and the lens, the lens face was then coated with a thin layer (∼50 μm thick) of Kwik-Sil, which was carefully spread out and shaped into a curved meniscus above the lens face. This Kwik-Sil layer was allowed to harden at room temperature overnight. Next, to activate the elastomer surface and make it more hydrophilic for binding to the hydrogel, the receptacle was filled with a drop of 2.5% potassium hydroxide solution (Sigma) for 1 hour at 37°C in the humidified cell culture incubator, rinsed 3 times in phosphate buffered saline (PBS, Thermo Fisher), and allowed to dry for 30 minutes at 37°C in a dehumidified incubator. Pre-coated lenses were then cooled back to 4°C in a cold room to prepare for hydrogel application. Photometry fibers were prepared in an identical manner.

#### Preparation and application of cell-laden hydrogel

After a 24-hour transfection, expression of the sensors was confirmed under an epifluorescence microscope (Zeiss Axio Observer D1). Cells were then treated with colchicine (1 μM, Sigma) for one hour to inhibit cell movement and proliferation. Colchicine-treated cells were then detached using TrypLE (Gibco) and centrifuged and resuspended in PBS twice (first for 8 minutes at 500 g, then for 10 minutes at 800 g) to remove any colchicine, debris, and culture media. After the second centrifugation, the supernatant was aspirated and cells were encapsulated in a hydrogel containing ∼60% collagen I (stock concentration ∼10 mg/mL; Corning) and ∼40% MATRIGEL® (stock concentration ∼20 mg/mL; Corning) by volume at a concentration of 10,000-50,000 cells/uL (for details of why this mixture was optimal, see below). To prepare this hydrogel, 200,000-1,000,000 cells were first pelleted in a 2 mL microcentrifuge tube (Eppendorf), which we found achieved optimally tight and dehydrated pellets. When preparing hydrogels with multiple types of sensor cells, the number of cells of each type included in the pelleted mixture was the total number of cells divided by the number of sensor types, so each sensor was represented at equal proportions and the total number of cells was always consistent. In a subset of experiments, we additionally mixed in the tdTomato+ HEK-293 cells (see above) at a ratio of 1:10 to the total cell number. These cells were used as landmarks during two-photon acquisition and analysis (mCherry signal from GRAB plasmids was very dim at 980 nm excitation and thus did not present any confound to this approach). The pellet was then suspended in 11.5 μL of stock collagen I (kept on ice) and 7.5 uL of stock MATRIGEL® (thawed from −20°C on ice for 25-30 minutes), and the solution was thoroughly homogenized with a pipette. Because the polymerization of MATRIGEL® and collagen was temperature-sensitive, the cell/hydrogel solution was immediately placed on ice and transferred to a 4°C cold room.

In the cold room, the cell-laden hydrogel was drawn into a pulled glass capillary tube (Drummond 0.4 mm), and 100-200 nL of the mixture was applied to each pre-coated GRIN lens receptacle under a dissection microscope. The lens was then transferred to a humidifying chamber (a closed container lined with Kimwipes soaked in ddH_2_O) kept at 37°C in a tissue culture incubator for gel polymerization. Extracellular matrix polymerization was allowed to proceed for 10 minutes, forming a ∼200 μm thick hydrogel. Immediately after polymerization of the hydrogel and adherence to the distal end of the probe, each fully manufactured MORSE probe was immersed vertically in a separate 1.5 mL microcentrifuge tube (Eppendorf) containing the same media used to culture the cells (DMEM + 10% FBS + 1% P/S) for 1-2 hours. Finally, the probes were transferred to a PBS-filled 1.5 mL tube and allowed to soak for at least 15 min before being used in an experiment. MORSE probes were used to characterize the composition of biological fluids *in vitro* (i.e. dipping), *ex vivo* (i.e. brain slices), and *in vivo* (i.e. two-photon imaging in CSF). MORSE photometry probes were prepared in an identical manner, except that HEK-293T cells expressing just one sensor were used per probe. These were applied *in vivo* for fiber photometry recordings in CSF.

### Optimization of MORSE manufacture

We iteratively optimized each component of the MORSE manufacture protocol. Key steps in this optimization are detailed below.

#### HEK-293T cell transfection timeline

We compared the GRAB sensor expression in cells transfected for several durations between 12 and 48 hours. Cells transfected for longer than 24 hours accumulated increased quantities of bright green intracellular aggregates. We interpreted these inclusions as reflecting endoplasmic reticulum (ER) stress and excess GRAB sensor retention. This strong intracellular sensor expression, which is not exposed to extracellular fluid, overshadowed the signal from membrane-bound GRAB sensor expression, preventing these cells from meaningfully responding to extracellular ligands. Conversely, cells transfected for much less than 24 hours were difficult to visualize under a two-photon microscope. Thus, a 24-hour transfection was optimal.

#### Hydrogel composition

In optimizing our hydrogel’s composition, we had to consider several competing factors. First, we needed a gel that was rigid enough to not warp excessively, but also permeable enough to support rapid coupling of surrounding fluid composition changes to sensor response. Second, a gel’s composition affects its adhesive properties, so we needed to consider how each change that we made to prevent warping would affect the gel’s attachment to the lens. We screened many different combinations of conditions and extracellular matrix (ECM) components (including MATRIGEL®, collagen I, fibronectin, and hyaluronic acid), before arriving at a 3:2 volumetric ratio of collagen I (stock concentration ∼10 mg/mL): MATRIGEL® (stock concentration ∼20 mg/mL). We also found that HEK-293T cells formed notably more stable hydrogels compared to HEK-293 cells at the same cell density and ECM component composition, presumably reflecting differences in membrane receptors and structural proteins in these closely related cell lines. For this reason, we kept the fraction of tdTomato-expressing HEK-293 cells in our probes to at most 10%.

At the molecular level, collagen I is a fibrous protein that forms a triple helix structure. This creates a rigid, yet porous matrix that bolsters connective tissues throughout the body^68^. Therefore, collagen I is typically thought to confer a hydrogel with rigidity, strength, and inflexibility^126^. This ECM was crucial in maintaining the structural integrity of MORSE and preventing hydrogel warping during recordings. However, hydrogels made up of more than 60% collagen I would more frequently fall off the GRIN lens face during *in vitro* dipping experiments. We interpreted this as likely due to the gels being too rigid and not adherent enough to the lens. In dipping experiments, the hydrogel must repeatedly break through the air-fluid interface in each successive well. The forces associated with overcoming this surface tension are applied directly to the gel. If the gel is insufficiently elastic, the force will be transmitted to the weakest part of the probe – the hydrogel-Kwik-Sil interface – causing detachment of the gel.

MATRIGEL® is a highly adherent ECM mixture containing laminin, collagen IV, entactin, and heparin sulfate proteoglycans – common components of basement membranes at epithelial tissue boundaries^127^. This combination of ECM tends to create meshworks that are more elastic than collagen I matrices; a given concentration of MATRIGEL® has an elastic modulus that is ∼3x lower than the same concentration of collagen I^128^. Because it is derived from murine sarcoma cultures, MATRIGEL® also contains many globular proteins, including transforming growth factor and epidermal growth factor^129^. These proteins, along with the heavily glycosylated proteoglycans, make MATRIGEL® appreciably stickier than collagen. Notably, despite its increased flexibility, MATRIGEL® may hinder molecular diffusion to a greater extent than collagen I due to viscous forces, which is a further advantage of our majority-collagen recipe^130,131^. The addition of 40% MATRIGEL® provided needed flexibility to our hydrogel, allowing it to withstand repeated exposure to the air-fluid interface while remaining attached to the Kwik-Sil. However, gels with greater than 40% MATRIGEL® warped excessively. In sum, we carefully balanced structural rigidity with adhesiveness, elasticity and porosity to achieve an optimal mixture.

#### Pre-coating of GRIN lens and photometry fiber face

We initially deposited the cell-laden hydrogel directly onto the face of our lens after it was glued to the surrounding PI tubing. However, due to small gaps between the lens and the surrounding PI tubing, air often became trapped under the hydrogel. Then, while the still-liquid hydrogel was polymerizing in the incubator, the trapped air expanded and formed bubbles that forced their way upwards into the gel before it solidified, forming crescent-shaped gels that were difficult to use. We discovered that filling these gaps and sealing the lens face with a thin layer (∼50 μm) of Kwik-Sil completely eliminated this problem. We found that it is best to perform this treatment at 4°C to ensure that the elastomer remains liquid for as long as possible as it fills the gaps, avoiding an analogous trapping of air bubbles. The Kwik-Sil coating does not noticeably impair cell visibility or image quality through the GRIN lens.

Adding Kwik-Sil to the front of the lens created another challenge: this silicone elastomer surface is hydrophobic and does not readily interact with the charged and polar groups in the proteins that make up our hydrogel, resulting in gel detachment. We tried several strategies to make the silicone surface more hydrophilic, including coating it with poly-D lysine, alginate, hyaluronic acid, polyacrylamide, chitosan, and polyethylene glycol, as well as treating it with oxygen plasma. Eventually, we settled on a relatively simple but effective solution that maintained gel adhesion to the GRIN lens: we pre-treat the Kwik-Sil coating with potassium hydroxide, which we hypothesize reacts with the Kwik-Sil to deposit polar hydroxide groups on the surface and roughens it to increase contact surface area, better anchoring the hydrogel to the lens surface^69,70^.

#### Colchicine treatment

We began by treating cells with colchicine for 24 and 48 hours following transfection^67^. However, these cells began developing bright intracellular aggregates spanning across the green and red fluorescence emission spectrum, likely reflecting large-scale intracellular protein denaturation indicative of excess toxicity. We found that a short one-hour treatment after transfection was sufficient to immobilize the cells for several hours of subsequent imaging without affecting sensor expression patterns or sensor function.

#### Animals

All animal experiments were approved by the IACUC and Institutional Biosafety Committees at Boston Children’s Hospital and Beth Israel Deaconess Medical Center. Wildtype C57BL/6J male and female mice aged 8 weeks used for lateral ventricle cannulation surgeries (8 male and 14 female mice aged 8-40 weeks), ICV AAV-GRAB sensor injections (8 male and 3 female mice aged 8-24 weeks), and cranial window/contralateral side port implantation surgeries following ICV AAV were ordered from The Jackson Laboratory (JAX). Swiss Webster mice used for CSF and blood collections (due to their larger brain ventricles) were ordered from Charles River (18 male mice aged 8-12 weeks). *FoxJ1-Cre::Ai95D* mice were bred by crossing *FoxJ1-Cre* mice with GCaMP6f-expressing *Ai95D(RCL-GCaMP6f)-D* mice (JAX*#* 028865) ordered from JAX (6 male and 3 female mice aged 8-60 weeks, all in C57BL/6 background)^132^.

*B6.Cg-7630403G23RikTg(Th-cre)1Tmd/J* (or *TH-Cre*) mice (JAX#: 00860) were crossed with *B6.Cg-Gt(ROSA)26Sortm32(CAG-COP4*H134R/EYFP)Hze/J* (or *Ai32*) mice (JAX #: 024109) to generate mice with channelrhodopsin2-YFP expressed in tyrosine hydroxylase-expressing monoaminergic neurons. These mice were used for brain slice experiments in **Figures 2 and S4** (5 male and 3 female mice aged 8-24 weeks). *B6.Cg-Tg(Fev-cre)1Esd/J* (or *Pet1-Cre*) mice (JAX #: 012712), which express Cre recombinase in Pet1+ serotonergic neurons, were injected with rAAV1/Syn-Flex-ChrimsonR-tdTomato in the dorsal raphe nucleus (DRN) and used for brain slice experiments in **Figures 2 and S4** (3 male and 3 female mice aged 8-20 weeks). All mice were given free access to food and water in an animal facility maintained at 70 ± 3°F and 35-70% humidity on a 12-hour light/12-hour dark circadian cycle. All experiments were carried out in darkness during the light cycle.

#### ChP imaging surgeries

For all surgical procedures performed for ChP imaging, male and female mice aged 8 weeks were administered meloxicam (10 mg/kg, subcutaneous), dexamethasone (1 mg/mL, intramuscular), and 0.9% saline (1 mL, subcutaneous) pre-operatively and anesthetized using 1-4% isoflurane in 100% O_2_. They were then positioned in a stereotactic frame.

#### Intracerebroventricular (ICV) injection of AAV5-GRAB

To infect the lateral ventricle ChP epithelium with a GRAB sensor, we first made an incision in the scalp and drilled a craniotomy at the coordinates +0.9 mm lateral and −0.6 mm posterior to Bregma. A pulled microcapillary tube (cat. no. P0549, Sigma Aldrich) was slowly lowered to −2.75 mm ventral to Bregma and used to administer 1 µL of virus directly into the lateral ventricle. AAV5-CAG-GRAB-5-HT2h.D, AAV5-CAG-GRAB-AVP0.2, and AAV5-CAG-GRAB-OXT1.7 (University of Pennsylvania Vector Core) were administered at titers of 7.12*10^11^ gp/mL, 2.75*10^12^ gp/mL, and 1.28*10^12^ gp/mL, respectively (AAV5 has a known tropism to ChP epithelium when delivered ICV in mice)^103^. Mice were then sutured and given at least 1 week to recover. Cranial window and contralateral infusion cannula surgeries were then performed on these animals (see below).

#### Dorsal raphe nucleus (DRN) viral injection

To infect DRN serotonergic neurons with ChrimsonR and tdTomato, we used *Pet1-Cre* mice and first made an incision in the scalp and drilled a craniotomy at the coordinates Bregma: AP: −4.5 to −5.1 mm, ML: 0 mm. A pulled microcapillary tube (cat. no. P0549, Sigma Aldrich) was filled with rAAV1/Syn-Flex-ChrimsonR-tdTomato virus (University of Pennsylvania Vector Core), lowered to the injection coordinates and injected into the dorsal raphe nucleus (200 nL at three locations, Bregma: AP: −4.55/-4.8/5.05 mm, DV: −2.9/-3.1/-3.3 mm, ML: 0 mm). Mice were sutured and given at least 2 weeks to recover prior to usage in brain slice experiments.

#### Cranial window and contralateral infusion cannula surgeries

Cranial window surgeries using *FoxJ1-Cre::Ai95D* or AAV5-GRAB-injected mice were performed as described in Shipley *et al.* 2020^100^. Briefly, a 3 mm diameter craniotomy was drilled over the left hemisphere (2.0 mm lateral and 0.2 mm posterior to Bregma), and the brain regions over the lateral ventricle were carefully aspirated. After hemostasis was achieved, a stainless steel cranial window with a sealed circular glass front face of 3 mm diameter was implanted over the large leaflet of the lateral ventricle ChP within the lateral ventricle. A stainless steel headpost was then affixed to the skin with VetBond (3M), and the cavity within the headpost was filled with C&B Metabond (Parkell). Mice were also outfitted with a custom-printed 3D imaging well (outer diameter of the base, inner diameter, height: 20 mm, 10 mm, 4 mm, respectively).

Optionally, a contralateral infusion cannula (“side port”) was also placed using the contralateral trans-occipital approach described previously^101^. Briefly, the neck muscles attached to the suboccipital bone were gently removed, and a stainless steel cannula (0.7 mm diameter; 12 mm length; Amuza AtmosLM PEG-12) was inserted into the contralateral lateral ventricle through an occipital craniotomy. A dummy cannula was then inserted and secured in place by a screw-on cap (Amuza AtmosLM PED-12 and Cap Nut AC-5). Mice were administered meloxicam (5 mg/kg, subcutaneous) and dexamethasone (4 mg/mL, intraperitoneal) daily for three days following surgery to ensure optimal recovery and minimize scarring around the implant. These mice were given at least 4 weeks to recover, after which they were used to infuse pharmaceuticals directly into the CSF while recording fluorescent signal from the ChP epithelium.

#### MORSE CSF recording surgeries

For all surgical procedures performed for MORSE CSF recordings, male and female mice aged 8 weeks were administered long-acting meloxicam (0.5 mg/kg, subcutaneous), dexamethasone (4 mg/mL, intramuscular), and 0.9% saline (1 mL, subcutaneous) pre-operatively and anesthetized using 1-4% isoflurane in 100% O_2_. They were then positioned in a stereotactic frame.

#### Bilateral anterior lateral ventricle cannula implantation surgery

Bilateral craniotomies of ∼1 mm in diameter were performed in wildtype C57BL/6 mice above both lateral ventricles (stereotactic coordinates: +/-1 mm lateral and −0.45 mm posterior to Bregma). To create an initial cannula tract, a beveled 21-gauge needle (PrecisionGlide, Fisher) was lowered −2.05 mm ventral to Bregma within each craniotomy. Custom cut stainless steel cannulae (22XX gauge, 7 mm length, Microgroup) were implanted to a depth of −2 mm ventral to Bregma and secured with Metabond. A headpost was then attached to the skin with VetBond, and the surgical site was sealed with Metabond. 7 mm-long pieces of stainless steel wire (27XRND, Microgroup) were used as dummy cannulae and secured with Kwik-Cast (World Precision Instruments). Mice were administered dexamethasone (4 mg/mL, intraperitoneal) daily for three days following surgery, as well as for two days prior to every MORSE recording, to minimize scarring and inflammatory response to the probe.

### Sample collection

#### Terminal CSF and serum collection

Wild type Swiss Webster mice were injected with AVP (1 mg/kg IP, Sigma), OXT (1 mg/kg IP, Bachem) or a 0.9% sterile saline (Medline Industries Inc.) control (100 µL total volume), or received a sham needle poke without injection. 30 minutes later, the mice were anesthetized with 100-120 mg/mL ketamine

/ 10 mg/kg xylazine (IP). A pulled glass capillary tube (Drummond 0.4 mm) was then inserted into the cisterna magna and CSF was collected with gentle negative pressure. Next, a thoracotomy was performed, and blood was collected from the cardiac left ventricle using a 20-gauge needle attached to a syringe. The collected CSF and blood were stored in mini tubes (Micro sample tube Serum, cat no. 41.1392.105, Sarstedt). CSF was then centrifuged at 3,000 g for 10 minutes at 4°C to remove any cellular debris. The supernatant was aliquoted and processed for ELISA (see below) or stored at −80°C. Blood was allowed to coagulate on ice for 30 minutes and centrifuged at 2,000 g for 30 minutes at 4°C. The top two thirds of the serum volume, avoiding the coagulated pellet, was aspirated, aliquoted, and processed for ELISA (see below) or stored at −80°C.

#### Imaging experiments

##### Pre-infusion

Mice with bilateral lateral ventricle cannulae were pre-infused as described in Fame *et al.* 2023^2^. Briefly, two weeks after surgery, 15 μL of aCSF (cat no. 3525, Tocris) was infused into one cannula over three minutes. The contralateral cannula was then pre-infused two days later, and mice were given at least a week to recover before being used for experiments.

##### Habituation

Mice were habituated to being head-fixed on a 3D-printed running wheel for increasing periods of time starting from 30 min to 4-5 hours, every day for at least 1 week leading up to an experiment.

##### Two-photon microscopy for MORSE recordings

Volumetric imaging of the cell-laden hydrogel in MORSE was performed through the GRIN lens using a resonant-scanning two-photon microscope (Neurolabware, Optotune, Scanbox v.11.0, 15.63 frames per second; 1,154 pixels x 512 pixels) equipped with an Insight X3 laser (Spectra-Physics). Probes were imaged at 980 nm excitation (<30 mW power measured below the objective) using a 10x super achromatic air objective (0.5 NA, 7.77 mm WD, ∼1000×500 μm^2^ FOV, Thorlabs). Volumes with 30 z-planes (∼9 μm between planes) were acquired at a rate of ∼0.5 vol/s. Both red and green fluorescence emission images were collected.

##### Probe dipping experiments

MORSE probes were calibrated and tested *in vitro* by sequential dipping into wells of a 384-well plate (cat no. 89014-066, Nunc® Shallow Well Standard Height Plates, black, Thermo Scientific) containing experimental solutions or standards of known identity and concentration. The objective of these *in vitro* dipping experiments was to (i) spatially demultiplex sensor signals, (ii) establish quantitative dose-response curves for each ligand, and (iii) detect many molecules in synthetic and biological samples. To perform these recordings, the GRIN lens was placed in a stainless steel micro-V-clamp (cat. VK250, Thorlabs) attached to a rack and pinion stage (cat no. 36-347, Edmund Optics; for vertical alignment) via a clamp mounted on the microscope arm (see **Figure 1D; Supplementary Video 2**). This custom lens holder was secured in focus below the microscope objective with the probe aligned vertically and the hydrogel facing downward.

Motors on the microscope arm were programmed to move the front of the lens through sequential dips into the wells of the 384-well plate (20 μL of fluid per well, 3-5 min/well and 15 sec/transfer, up to 80 wells/recording). Calibration wells contained buffered saline, near-saturating concentrations of one of up to 16 sensor ligands individually, or one of 6 logarithmically spaced concentrations of 6-8 ligands mixed together. These calibrations were performed at room temperature, and dose-response patterns may differ slightly from sensor responses to similar concentrations of ligand in the heated slice chamber or *in vivo*.

##### Acute brain slice preparation

Brain slices were prepared by deeply anesthetizing mice (12-20 weeks old, ad libitum fed) with isoflurane, followed by transcardial perfusion with ice-cold, carbogen-saturated (95% O_2_, 5% CO_2_) aCSF consisting of (in mM): 126 NaCl, 27.4 NaHCO_3_, 2.5 KCl, 1.2 NaH_2_PO_4_, 1.2 MgCl_2_, 2.4 CaCl_2_, 10.5 glucose. The mice were then rapidly decapitated, and the brains were extracted. Upon removal, brains were immediately immersed in ice-cold, carbogen-saturated choline-based cutting solution consisting of (in mM): 92 choline chloride, 20 HEPES, 2.5 KCl, 1.25 NaH_2_PO_4_, 30 NaHCO_3_, 25 glucose, 10 MgSO_4_, 0.5 CaCl_2_, 2 thiourea, 5 sodium ascorbate, 3 sodium pyruvate. Osmolarity was measured at 310-320 mOsm/L and pH at 7.40. Next, 300 μm-thick coronal (for PVH) or sagittal (for dLGN) sections were cut with a vibratome (Campden 7000smz-2) and incubated in oxygenated cutting solution at 35°C for 15 min. Slices were subsequently transferred to oxygenated aCSF at 35°C for an additional 30 min. Slices were then kept at room temperature (RT, 20-24°C) for ∼30 min until use. For experiments with wash-ons of high KCl, a second aCSF solution was made with 52.5 mM KCl and 77 mM NaCl to maintain osmolarity. Before the experiment, a single slice was placed in the recording chamber under the two-photon microscope (Neurolabware), where it was continuously superfused with fresh oxygenated aCSF via a peristaltic perfusion pump (Cole Parmer Masterflex 77120-52 C/L) at a rate of ∼4 mL per min at RT. The target region on the slice was identified using live imaging with the two-photon microscope (locating YFP-positive axons in *TH-Cre::Ai32* slices to find the PVH and tdTomato-positive axons in *Pet1-Cre*/DRN AAV-DIO-ChrimsonR slices to find the dLGN). A dissection microscope (Olympus) was then set up to observe the slice from an oblique angle, and a MORSE probe was attached to the two-photon microscope under the objective with a custom lens holder (see above). The probe was observed with live two-photon imaging and secured with the hydrogel in the focal plane of the objective, and then the whole microscope arm was lowered to press the MORSE probe against the slice region of interest. Correct targeting was achieved by direct observation of the slice through the angled dissection microscope as the probe was lowered onto it, and z-control on the microscope arm was used to position the end of the probe to an indentation of 100-150 µm below the slice surface. Two-photon imaging of the cell-based sensors pressed against the slice was then performed while superfusing aCSF containing different ligands onto the slice. The flow in the recording chamber was tested to confirm fluid exchange timings: at 4 mL/min pump setting, the time to completely wash out a pigmented solution from the chamber and replace it with clean inflowing aCSF was approximately 5 min.

##### In vivo MORSE experiments

At least one week after pre-infusion (see above), mice with bilateral lateral ventricle cannulae were head-fixed for two-photon imaging. Both dummy cannulae were acutely removed and a ∼1 µL drop of CSF was allowed to accumulate at the top of each cannula. Only mice possessing optically clear CSF (indicating unobstructed cannula placement in a relatively healthy ventricle with minimal post-surgical bleeding) were used for recordings. A MORSE probe was removed from a PBS-filled 1.5 mL tube and carefully introduced into the cannula with better CSF flow, ensuring no air bubbles formed at the hydrogel-CSF interface. The lines drawn at 7 mm from the hydrogel end of the MORSE probe were used to guide the probe to the correct depth. The implanted probe was then sealed to the cannula with Kwik-Sil.

In a subset of experiments, a polyethylene catheter (inner diameter 0.28 mm, outer diameter 0.61 mm, cat no. BD 427401, Fisher Scientific) pre-filled with serotonin solution was introduced into the contralateral ventricular cannula for ICV infusions and sealed with Kwik-Cast. Two-photon imaging of the cell-based sensors was then performed through the GRIN lens while the mouse ran on a running wheel and was subjected to pharmacological (delivered IP and ICV) stimuli. Each session involved a 30 minute acclimation period, followed by 3-4 hours of two-photon imaging. Drugs were mixed in PBS (Thermo Fisher) for IP injections of 100-300 µL volumes performed with an insulin syringe (Fisher) and mixed in aCSF (Tocris) for ICV infusions in volumes of 15-30 µL at a rate of 5 μL/min using a syringe pump (cat no. 70-4504, Harvard Apparatus).

##### In vivo MORSE photometry experiments

At least one week after pre-infusion (see above), mice with bilateral lateral ventricle cannulae were head-fixed for photometry recordings. Both dummy cannulae were acutely removed and a ∼1 µL drop of CSF was allowed to accumulate at the top of each cannula. Mice lacking optically clear CSF were not used for recordings. A MORSE photometry probe was removed from a PBS-filled 1.5 mL tube and carefully introduced into the cannula with better CSF flow, ensuring no air bubbles formed at the hydrogel-CSF interface. The probe was pushed all the way into the cannula until the fiber ferrule was flush with the cannula opening. The implanted probe was then sealed to the cannula with Kwik-Sil. Zirconia sleeves (Precision Fiber Products) were used to couple the fiber to a patch cord (2 m length, NA 0.57, Doric MFP_400/430/1100-0.57_2m_FC-MF1.25_LAF) that had been photobleached overnight.

All experiments were carried out using the fiber photometry setup described in^133^. Briefly, a 465 nm LED (∼100 μW measured at the end of the patch cord, Plexon OPT/LED_Blue_TT_FC) excited the GRAB sensor for signal detection, and a 405 nm LED (∼100 μW measured at the end of the patch cord, Plexon OPT/LED_Ultraviolet3_TT_FC) was used to collect an isosbestic control signal. Excitation light pulses were delivered at 50 Hz and included 6 ms of 465 nm LED, 4 ms of darkness, 6 ms of 405 nm LED, then 4 nm of darkness, controlled by Plexon LED drivers (OPT/1ch_LED_Driver) via the Nanosec photometry-behavioral system^133^. Light from both LEDs converged into the patch cord via a four-port fluorescence mini cube (Doric, FMC4_IE(400-410)_E(460-490)_F(500-550)_S). The fluorescence intensity emitted from the GRAB sensors was detected by a femtowatt photoreceiver (Newport+Doric FC Adapter, NPM_2151_FOA_FC). The recordings were conducted in darkness, and black heat shrink tubing was placed around the fiber-patch cord coupling region to prevent the LED light from being seen by the mice.

Photometry recordings of the cell-based sensors were then performed through the optical fiber while the mouse ran on a running wheel and was subjected to pharmacological (delivered IP and ICV) stimuli. Each session involved a 30 minute acclimation period, followed by 2-3 hours of photometry recording. Drugs were mixed in PBS (Thermo Fisher) for IP injections (100-300 µL volume) performed with an insulin syringe (Fisher) and mixed in aCSF (Tocris) for ICV infusions in volumes of 15-30 µL at a rate of 5 μL/min using a syringe pump (cat no. 70-4504, Harvard Apparatus). The table below summarizes the pharmaceuticals injected IP and ICV in the two-photon and photometry MORSE experiments:

| Pharmaceutical | Dose(s) |
| --- | --- |
| 5-hydroxytryptamine (5-HT, Sigma) | 2-3 mg/kg (IP)<br>5 nmol/μL infused at 5 μL/min over 6 min (ICV) |
| Arginine vasopressin (AVP, Sigma) | 1 mg/kg (IP) |
| Oxytocin (OXT, Bachem) | 1 mg/kg (IP) |
| Norepinephrine (NE, Sigma) | 1 mg/kg (IP) |
| Dopamine (DA, Sigma) | 1 mg/kg (IP) |
| Cholecystokinin (CCK, Bachem) | 1 mg/kg (IP) |
| Acetylcholine (ACh, Sigma) | 10 mg/kg (IP) |
| Melatonin (MT, Sigma) | 10 mg/kg (IP) |
| Histamine (HA, Sigma) | 10 mg/kg (IP) |
| Adenosine triphosphate (ATP, Sigma) | 50 mg/kg (IP) |

##### Two-photon imaging of lateral ventricle ChP

Imaging of *FoxJ1-Cre::Ai95D* and ICV-AAV5-GRAB-injected mice was performed as described in Shipley *et al.* 2020^100^. Briefly, mice were habituated to being head-fixed on a running wheel for 30-240 min/day for at least 1 week. On the day of the experiment, mice were head-fixed on the running wheel directly under a resonant-scanning two-photon microscope (Olympus MPE-RS Multiphoton microscope, 18.75 frames/s, 512×512 pixels/frame). ICV-AAV5-GRAB-injected mice received an IP injection of 25 mg/kg of 70 kDa Texas Red™ Dextran (Thermo Fisher) 30 minutes before the recording to visualize the ChP vasculature. The imaging well on the mouse’s head was filled with ultrasound gel, and a 25x, 0.95 NA, 8 mm WD objective (Olympus XLSLPLN25XSVMP2) at 2.5x zoom (∼203 × 203 μm^2^) was used to view the lateral ventricle ChP. Volume scanning was achieved by using a piezo microscope stage (nPFocus250, nPoint) moving axially in a sawtooth pattern.

*FoxJ1-Cre::Ai95D* and ICV-AAV5-GRAB were imaged at 940 nm excitation (Mai Tai DeepSee laser, Spectra Physics) and 3D volumes (81 z-planes, 5 μm/plane, stack volume 203 × 203 × 400 μm^3^) were collected at a rate of 1 volume/4.32 seconds for 2-3 hours. The power under the objective was measured to be between 34-45 mW. An IP injection of a ligand/antagonist was administered around 60-90 minutes into the recording. Optionally, in mice receiving ICV injections, a polyethylene catheter (inner diameter 0.28 mm, outer diameter 0.61 mm; cat no. BD 427401, Fisher Scientific) pre-loaded with a ligand was threaded 12 mm into the side port and secured using Kwik-Cast. 15 μL of the ligand diluted in aCSF was infused at a rate of 5 μL/min using a syringe pump (cat no. 70-4504, Harvard Apparatus). The table below summarizes the pharmaceuticals injected IP and ICV in two-photon ChP imaging experiments:

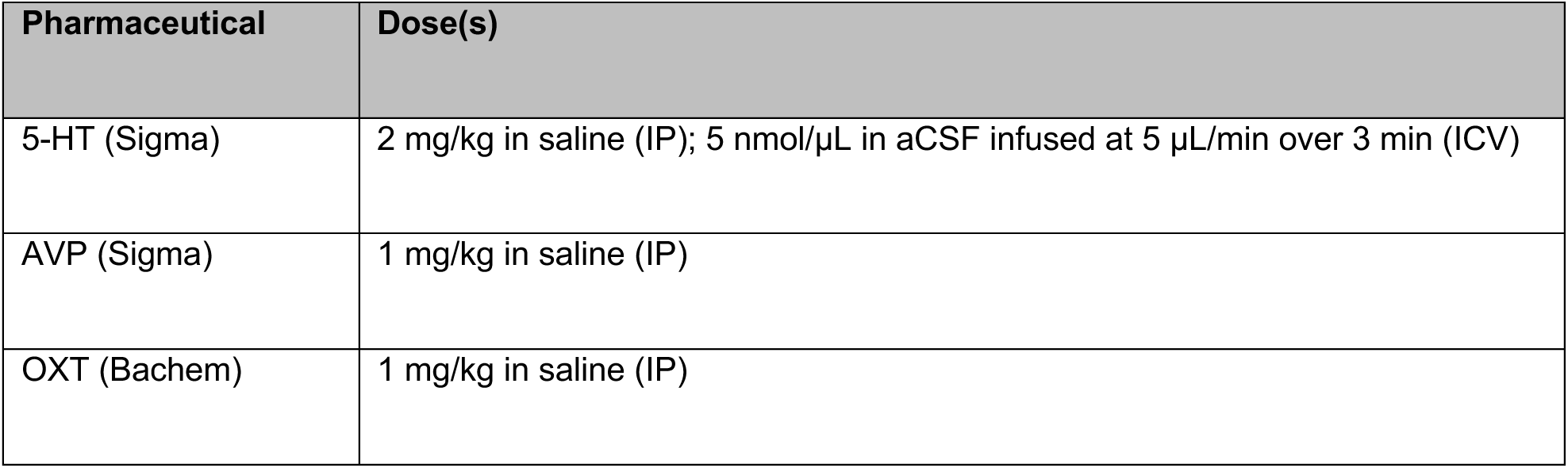

### Post hoc tissue and fluid analysis

#### CSF and blood ELISA

Blood and serum samples were collected as described above. The samples were processed further according to the manufacturer’s protocols: DetectX® Arg8-Vasopressin Enzyme Immunoassay Kit, cat no. K049-H1, Arbor Assays for AVP quantification and DetectX®, Oxytocin Enzyme Immunoassay Kit, cat no. K048-H1, Arbor Assays for OXT quantification.

#### Perfusion and histology

Mice were deeply anesthetized with ketamine (100-120 mg/mL IP) and xylazine (10 mg/kg IP) or with Avertin (tribromoethanol, 125 mg/kg IP). A thoracotomy was performed, and the cardiac left ventricle was perfused with 10 mL of PBS followed by 10 mL of 4% paraformaldehyde or 10% formalin. Brains were extracted, allowed to fix in 4% paraformaldehyde or 10% formalin for 24 hours, and moved to 20% sucrose for 24 hours. 60 μm-thick coronal sections were then prepared and stored in PBS.

#### Immunostaining and image acquisition

Histology was performed on a representative subset of mice used for *in vivo* experiments to confirm cannula placement and evaluate inflammation. Slices were blocked in 1% normal donkey serum, 0.05% sodium azide, and 0.05% Triton-X on a shaker at room temperature for 2 hours. Rat anti-GFAP (1:200, cat no. 13-0300, Invitrogen) antibodies diluted in the blocking solution were then used to stain the tissue, which was kept on a shaker at 4°C overnight. The tissue was then washed three times with PBS and stained with AlexaFluor594 donkey anti-rat (1:500, cat no. A-21209, Invitrogen) for 2 hours on a shaker at room temperature. Stained slices were mounted on slides using VectaShield® HardSet™ with DAPI (cat no. H-1500-10, Vector Labs). Slides were imaged using a VS120 slide scanner microscope (Olympus).

### Two-photon image analysis

#### MORSE hydrogel deformation quantification and image stabilization

This process of MORSE hydrogel deformation quantification is diagrammatically outlined in **Figure S1A, left**. Due to the relatively slow (tens of seconds) timescales of the ligand calibration responses *in vitro* and *in vivo*, as well as the relatively slow progress of hydrogel warping, we began by creating temporally-downsampled (TDS) MORSE recordings by taking means of every 31-52 volumes (down to 1 volume every ∼60-100 sec). For a limited subset of image planes from these TDS movies, we annotated individual cells by hand and trained a neural network model for 3D cell segmentation using Cellpose2.0^60^. We then applied this model to both the green (GRAB sensor) and red (tdTomato, in a subset of recordings) color channels of every TDS MORSE movie at every timepoint to generate probability maps estimating how likely each pixel is to be part of a cell (**Figure S1B**). Next, we performed nonlinear optical flow-based registration on these probability maps derived either from the green or red channel to a single reference timepoint, usually the middle timepoint of the movie (compute time ∼5 sec/volume with GPU acceleration on Nvidia RTX3090 using the imregdemons function in MATLAB2023b, 4 Pyramid Levels, Accumulated Field Smoothing 2.0) (**Figure S1C**). To stabilize the recordings using these calculated hydrogel deformations, we linearly interpolated the resulting displacement fields up to the sampling frequency of the original raw movies (∼0.5 vol/sec), applied the transformations to the raw movies, and corrected high-frequency intervolume translations by registering sets of 31-52 adjacent volumes using cross-correlation of the 3D Fourier transform to create aligned raw movies (compute time ∼150 sec/stabilized volume using the imwarp function in MATLAB2021a on Intel Xeon E5-2650)^100,134^. Stabilization success was quantified by reapplying the Cellpose2.0 model to temporally-downsampled versions of the aligned raw movies and calculating the cross-correlations of the GRAB signal probability maps at each timepoint to the first timepoint, restricted to the voxels identified as having at least a 50% likelihood of being part of a cell at the first timepoint (**Figure S2C**).

#### MORSE image segmentation and signal extraction

This process of MORSE single-cell trace extraction is diagrammatically outlined in **Figure S2A**. To estimate signals from single-cell regions of interest (ROIs), we generated temporally-downsampled aligned (TDSA) MORSE recordings by taking means of every 31-52 volumes of the stabilized raw movies from above and created Cellpose probability maps for these using the same neural network model as above. This time, we created maximum intensity projections of the probability maps over time and binarized them by identifying pixels that had greater than 50% likelihood of being part of a cell in at least one volume over the course of the TDSA movie. We masked out these pixels in the TDSA movie, spatially-downsampled the remaining pixels by a factor of 16 in the x and y dimensions, and performed 3D thin plate spline interpolation across the whole volume at each timepoint (python function scipy.interpolate.RBFInterpolator with degree=1 and smoothing=100), followed by Gaussian filtering in the x and y dimensions (σ = 1.2 pixels) and upsampling back to the original spatial dimensions of the TDSA movie. We then subtracted this interpolated background from each TDSA movie to create the temporally-downsampled background-corrected aligned (TDSBA) movies.

For a preliminary estimation of cell masks, we applied the Caiman CNMF (constrained nonnegative matrix factorization) algorithm for fluorescence signal extraction from cells, which performs spatial and temporal unmixing of distinct signal time courses in local regions, to the TDSBA movies (p=0 autoregressive order, gnb=0 global background components, rf=60 pix patch half-size corresponding to roughly 60*60 µm^2^ patches in the x-y dimension across the full z-thickness of the recording, K=30 cell components per patch, gSig=[10,10,1.25] pix half-widths of expected cells across the x-y-z dimensions corresponding to [10,10,11.25] µm, method_init=’greedy_roi’ initialization method, ssub=tsub=1 indicating no spatial subsampling during initialization). Each movie was thus separated into 30 3D spatial bins of equal size evenly tiling the volume, with 30 putative cell signal ROIs extracted per bin for 900 total ROIs (compute time ∼20 min/movie parallelized over 30 CPU cores on Intel Xeon E5-2650). ROIs possessing a minimum signal above a reasonable threshold (calculated as 1.96 times the standard deviation of the distribution of ROI baseline signals with negative values of background-corrected fluorescence signal, assumed to be a half-normal distribution) as well as high signal-to-noise ratio (calculated as the ratio of the maximal baseline-corrected signal at the ROI over time divided by the standard deviation of the signal around a Savitzky-Golay-smoothened version of the signal, with ROIs possessing ratio values above the 20-25^th^ percentile of all ROIs included) were retained for downstream analysis. The ∼450-500 ROIs that passed this step were then manually curated to distinguish signals from individual cells using a custom GUI built in napari (masks were discarded or retained and possibly merged into groups, manual time ∼2 hrs/movie)^73^. Additional quality control was subsequently performed using another custom napari GUI to ensure that any previously discarded ROIs corresponding to true cells were retained (manual time ∼1 hr/movie). The curated ROIs were then refitted to the TDSBA movies using Caiman to yield estimated cell masks and time courses (compute time ∼2 hrs/movie on Intel Xeon E5-2650).

Finally, to apply these calculated cell masks back to the aligned raw movies, we first linearly interpolated the estimated background signals up to the sampling frequency of the aligned raw movies and subtracted them to create aligned raw background-corrected movies. We then linearly interpolated the estimated time courses for each cell mask in the TDSBA movies up to the raw sampling frequency and refitted them to the aligned raw background-corrected movies using Caiman, resulting in a final set of single-cell masks with time courses at native temporal resolution and corrected for background (compute time ∼3 hrs/movie on Intel Xeon E5-2650).

#### MORSE cell matching across recordings

Many of the MORSE probes were imaged several times in multiple contexts *in vitro* and *in vivo*. We performed cell mask matching between different recordings of the same probe to track cell identities and activity patterns across recordings. Since the probes were moved and imaged in different orientations, we began by calculating transformations between recordings. We extracted the single reference volume used for registration of each TDS movie of a given probe (see above) and chose one of them as the target for cross-movie alignment. Using the BigStitcher plugin in ImageJ, we then manually rotated and translated each of the other reference volumes to best align with this alignment target volume (manual time ∼10 min/probe)^75^. For most probes, this was not enough for satisfactory alignment, as the recordings were performed with slight variations in vertical alignment of the probe relative to the microscope objective, and the hydrogels also exhibited some deformation in between imaging sessions. To address this, we further used the BigWarp plugin in ImageJ to manually match 30-40 anchor points between every pair of reference volumes aligned previously and produced a thin plate spline deformation field to achieve fine-scale alignment (manual time ∼20 min/pair of volumes) (**Figure S2D**)^74^. We then applied these deformation fields to the set of masks from the corresponding recordings of the probe and overlaid the outputs with the masks from the alignment target movie, successfully overlapping many masks with their correct partners (**Figure S2E**). Finally, we manually curated these overlapped mask matches to confirm correct cell identities using another custom GUI in napari (manual time ∼2 hrs/pair of movies) (**Figure S2F**).

#### MORSE signal processing

We aimed to calculate dose-response parameters for each sensor to estimate quantitative concentrations of the ligands and unmix the dopamine-norepinephrine sensor crosstalk. To do this, we assumed that sensor-ligand binding kinetics were sufficiently rapid to model the process as a classic equilibrium reaction, with S representing the free sensor, L representing the free ligand, S•L the bound sensor-ligand complex, and K_d_ the dissociation constant between them^135^:

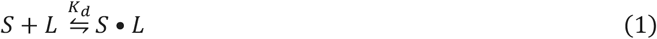

In this case, the standard equilibrium expression, where [S], [L], and [S•L] are the concentrations of free sensor, free ligand, and bound sensor-ligand complex, respectively, holds:

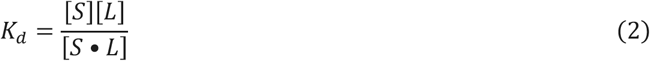

If we assume that the S and S•L forms of the sensor have stable fluorescence brightnesses α and β, respectively, and that the recording also includes a ligand-independent background signal ε (all three dependent on imaging parameters like excitation light intensity and detector gain), then we can express the fluorescent signal F as follows:

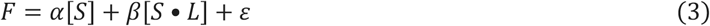

If we assume that the total concentration of the sensor [S]_0_ = [S]+[S•L] is constant, we can define fractional sensor occupancies as follows:

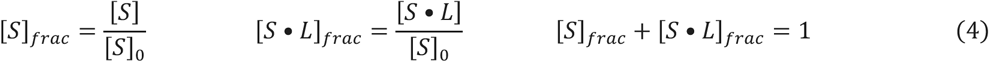

Finally, if we define baseline ligand-free signal F_0_ = α[S]_0_+ε and normalized brightness constant β_norm_ = [S]_0_(β-α)/F_0_, we can express the fractional change in fluorescence as:

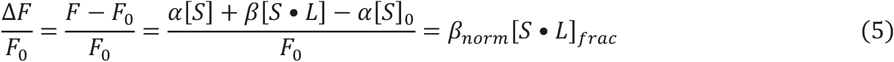

with β_norm_ equaling the peak ΔF/F_0_ response to a saturating dose of L that brings [S•L]_frac_ asymptotically close to 1. This means that peak-normalizing the ΔF/F_0_ is equivalent to dividing by β_norm_:

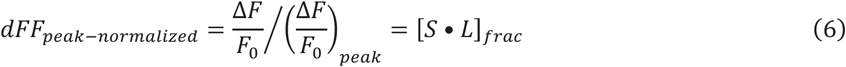

Writing out the equilibrium reaction in terms of fractional occupancies then yields:

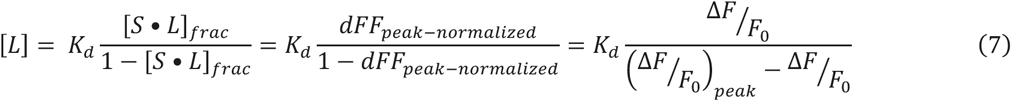

This expression can be used to calculate K_d_ for each sensor by fitting a linear regression to known concentrations [L] and corresponding sensor responses dFF_peak-normalized_. For completeness, the applied concentration of L, [L]_0_, is the sum of free and bound L, i.e. [L]_0_ = [L]+[S•L] = [L] + [S]_0_[S•L]_frac_ so for [L] to be a faithful reflection of [L]_0_, we must assume conditions of little buffering of free ligand by the sensor, i.e. [S]_0_ << K_d_.

In the case of dopamine-norepinephrine crosstalk, the model would include two ligand-bound states with different equilibrium constants:

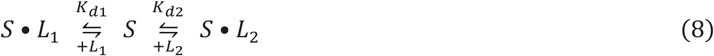

In that case, the fluorescent signal is:

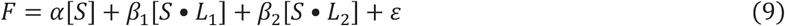

With analogous definitions of fractional occupancy and brightness, we then have:

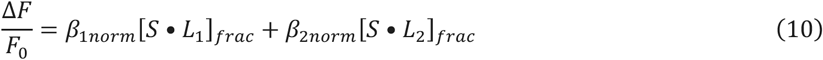

Normalizing ΔF/F_0_ by the saturating response to L_2_ and defining γ = β_1norm_/β_2norm_ yields:

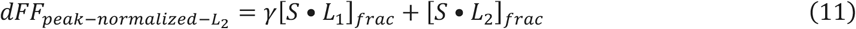

At this point, we can derive all the sensor response parameters as above by applying escalating doses of one ligand at a time and performing an analogous regression, e.g.

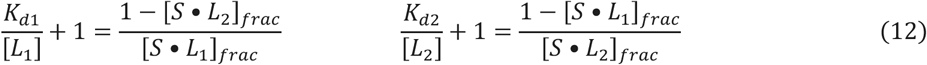

For any subsequent recording using the probe, we would then collect two sensor signals (e.g. DA2.9 and NE2h) with two unknowns (e.g. [DA] and [NE]) and have all the parameters necessary to solve the resulting system of equations.

On top of that, because we perform identification and dose-response dipping recordings on all ligands at the same time, we can in principle determine the ratios between their ΔF/F_0_ peaks (i.e. their β_norm_ values), and thus in future recordings with this probe we would likely only need one ligand to be applied at saturating dose to estimate the expected peak responses of all the other sensors.

Importantly, all of the above derivations hold for the sensor at steady state, once the concentration throughout the hydrogel has equilibrated to the concentration in the outside fluid. However, in order to reach the sensor protein, each ligand must diffuse through the thin hydrogel layer. Modeling this diffusion process would allow us to obtain the transfer function of the hydrogel, which we could deconvolve from the observed signal to obtain a better estimate of ligand concentration in the surrounding fluid. It should be true that if the probe is placed into a relatively large volume of fluid with the ligand at a given concentration (as it happens during our dipping characterization), the ligand concentration inside the hydrogel will eventually plateau to that in the fluid. This diffusion process is expected to follow Fick’s second law. If we define [L](d,t) to be the ligand concentration at a sensor positioned at a depth d in the hydrogel over time, h as the total depth of the hydrogel, and [L]_f_ as the constant concentration of the fluid that the probe is placed into at t=0, the solution to Fick’s second law for a cylindrical hydrogel with one open end is the following^136^:

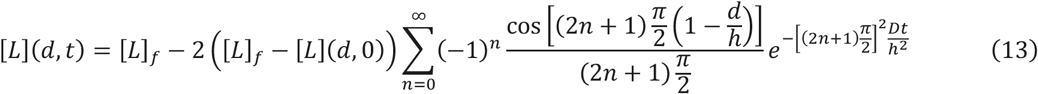

Rearranging, we get

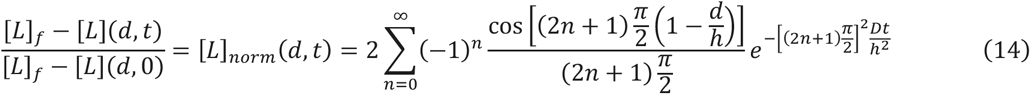

This is an infinite sum of exponentially decaying terms, with the n=0 term having the slowest decay and smallest factor in the denominator (although higher n components may have greater multiplicative weights depending on d and the factor in the numerator). In this case, fitting an exponential decay to this relationship will result in decay constant

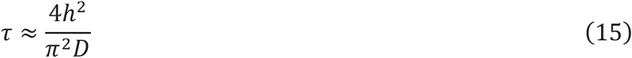

The exact value will vary with d, but will always be independent of surrounding fluid concentration [L]_f_. It is critical to note that this τ calculated on [L]_norm_ is not equivalent to an exponential fit on any linearly normalized version of the fluorescence signal since [L] is nonlinearly related to dFF_peak-normalized_ as described above. When L is applied on its own, ln([L]) is approximately linearly related to fluorescence around the K_d_ of the sensor:

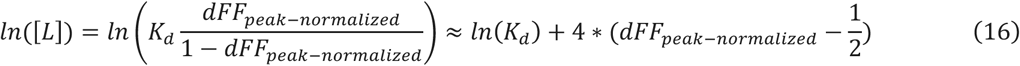

in which case [L] can be estimated by taking the exponential of both sides of the equation, and that result can be used to assess diffusion:

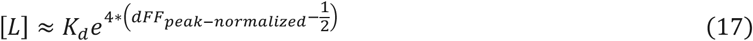

However, this is not true where [L] is substantially smaller or larger than K_d_ – in those cases, exponential fits on the result will be *underestimates* of diffusion speed (i.e. overestimates of τ) at low [L] and *overestimates* of diffusion speed (i.e. underestimates of τ) at high [L]. In particular, this means that calculating probe response kinetics using saturating concentrations of [L] will *underestimate* the magnitude by which the hydrogel slows down responsiveness if the nonlinearity is not correctly accounted for.

Additionally, these relationships depend on the assumption that ligand binding kinetics at the sensor are much faster than diffusion through the hydrogel, which we assumed to be the case for most of the GRAB sensors we used based on published findings^39,43,45,58,137^.

Now, given this concentration-independent delay constant τ relating the concentration measured at the sensor to the external concentration in the fluid immediately surrounding the probe, we can estimate the hydrogel’s transfer function. Importantly, this depends on the assumption that the external concentration [L]_e_(t) (distinct from [L]_f_ above because it is now varying over time) is uniform throughout the fluid facing the probe (likely true in CSF). In that case, we can model the hydrogel akin to a classic RC circuit, with the external concentration [L]_e_(t) analogous to the input voltage and the sensor concentration [L](t) analogous to the voltage across the capacitor. This would imply that in the frequency domain,

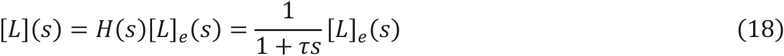

with transfer function H(s) = 1/(1+τs) and corresponding impulse response

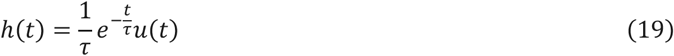

with u(t) the Heaviside step function. Deconvolving this impulse response from [L](t) (possibly with an appropriate Savitzky-Golay or Wiener filtering step to reduce noise) will then estimate [L]_e_(t) in situations that satisfy the above assumptions.

#### ChP image registration

Registration of the volumetric ChP images was performed as described in Shipley *et al.* 2020^100^. Briefly, due to rapid movement of the mouse, it was necessary in some recordings to account for intra-volume shifts of the tissue. This was achieved by aligning each plane of the volume to the brightest plane, which acted as an anchor, using Fourier cross-correlation. Next, volumes were registered to each other using cross-correlation of the 3D Fourier transform^100,134^.

#### ChP signal processing

Following registration, a representative 3D subvolume of the recording was extracted by choosing x, y, and z dimension ranges that lacked any voxels cropped during registration. A mean projection of this volume across the z dimension was then produced, yielding a two-dimensional timelapse movie. A representative square region of interest spanning several ChP epithelial cells expressing the sensor of interest (GCaMP6f or a GRAB sensor) was selected and the mean green signal trace over the region across time was obtained. In recordings demonstrating slow signal decay prior to ligand administration, an exponential fit to the signal preceding ligand administration was obtained and subtracted from the trace to obtain the estimated change in fluorescence (ΔF) which was divided by the mean raw signal over the first 5 minutes of the recording to estimate the ΔF/F_0_.

#### Quantification and statistical analysis

In the case of two-photon recordings, processed single cell fluorescence signals from MORSE or tissue signals from ChP imaging were normalized to a baseline corresponding either to the mean signal over the 15-minute period preceding ligand administration (in the case of IP or ICV injections) or to the mean signal of the lowest 10% of readings over the entire recording for each individual trace as follows:

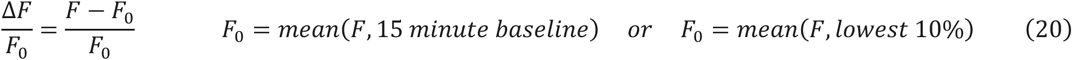

In the case of dipping recordings, processed fluorescence signals from MORSE that were normalized as above to obtain a ΔF/F_0_ signal often demonstrated a sharp upward jump in the signal at the start of every well, likely due to forces applied to the hydrogel as it crossed the air-fluid interface. This artifact was fitted with an exponential decay scaled by the jump in the ΔF/F_0_ signal between adjacent wells, which exhibited a consistent decay constant of ∼30 sec (see **Supplementary Video 3** for an example of such jumps). The artifact-corrected signal was then either fitted with an exponentially decaying background correction over tens of minutes of signal preceding sensor activation (single-dose sensor identification dipping as in **Figure 1E**, with activation identified by a spike in the sensor’s estimated first derivative – see below) or processed with a Butterworth filter with a passband in the 1/6000 to 1/50 Hz range to accentuate responses on the 3-5 minute timescale of the dipping process while discarding slowly changing background signal (slice and *in vivo* dipping characterizations in **Figures 2-4 and S4**, except for KCl dose-response dipping in **Figures S4E-F** which was not baseline corrected). Dipping characterization signals were subsequently reported in peak-normalized form (except for KCl dose-response dipping in **Figures S4E-F**), which was calculated as follows:

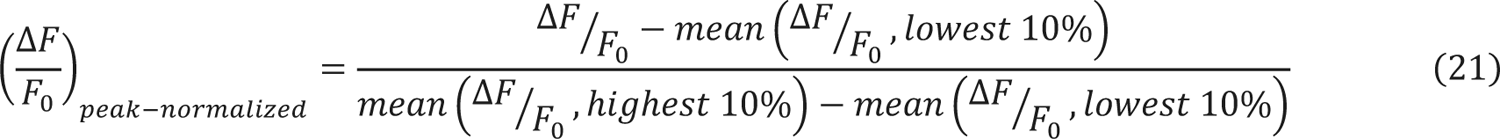

We found that the most effective approach to clustering cells by sensor identity was to focus on the onset timings of each standard ligand administration (either dipping into an identification well or washing on a saturating ligand dose onto the slice), since different sensors and ligands exhibited variable wash-off kinetics. To do this, the peak-normalized ΔF/F_0_ signal calculated as above (with or without background correction or Butterworth filter) was processed with a Savitzky-Golay filter with a 1-minute window and polynomial order 1, which estimated the first derivative. This resulted in an estimate of sensor activation time, as the filtered signal had large values at the times when the signal was rising the fastest (**Figures 1E, bottom and S3A-C**). We subsequently performed hierarchical density-based spatial clustering of applications with noise (HDBSCAN) on the derivative time courses^138^. This yielded high-quality clusters which were automatically matched to corresponding ligand identities by calculating the cross-correlation of the derivative time courses with the time courses of ligand administration. Calculating the means of the derivative time courses across each cell cluster and cross-corelating them with each other consistently demonstrated that only the DA2.9 dopamine and the NE2h norepinephrine sensors, as well as to a lesser extent the AVP0.2 vasopressin and OXT1.7 oxytocin sensors, exhibited crosstalk in their ligand responses (**Figures 1F and S3D**)^43^. In the slice and *in vivo* recordings, we performed similar HDBSCAN clustering on the ΔF/F_0_ signals or peak-normalized ΔF/F_0_ signals to cluster the cell responses.

For the assessment of sensor K_d_ (i.e. EC_50_) and saturation, ΔF/F_0_ signal from each cell from the dose-response dipping recording that was matched to a sensor identity from an identification dipping recording of the same probe was first corrected with an exponentially decaying background fit and then peak-normalized as above. Next, the mean of the sensor signal from the third minute of recording in a given well with a given concentration of the ligand was taken as the sensor’s plateaued signal at that concentration. These values were fitted with a dissociation equation as follows:

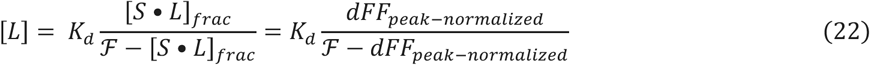

where K_d_ is the dissociation constant and ℱ is the predicted maximal dFF_peak-normalized_ signal in the sensor (should be close to 1 if the sensor was subjected to a full dose-response and peak-normalized to a near-saturating concentration signal).

In a subset of slice and *in vivo* MORSE recordings, these sensor parameters were subsequently used to obtain quantitative estimates of molecular concentrations. To achieve this, the signals were first peak-normalized. In the case of the slice recordings, in which the saturating wash-on concentrations of NE, DA, and 5-HT were all 50 µM, the peak-normalization was performed to this reference concentration – in other words, the calculated K_d_ and ℱ parameters for each sensor were used to estimate the dFF_peak-normalized_ signal expected for a 50 µM administration of the ligand individually, and the signals were then scaled for their post-wash-on peaks to match this value. For *in vivo* 5-HT quantification, the delivery of 15-30 µL of 5 mM stock 5-HT in aCSF into a ∼36 µL volume of endogenous mouse CSF translates to a final 5-HT concentration of at least 1 mM, which should be more than sufficient to completely saturate the sensor^139^. Therefore, in this case the signals were peak-normalized directly to the ℱ parameter of the 5-HT sensor.

Concentration estimates of NE and 5-HT over time were then obtained by applying Eq. 22 to the normalized time course of each sensor. In the case of the DA sensor, which demonstrated strong crosstalk with the NE ligand, a modified expression based on the sensor’s combined response to both ligands (Eq. 11-12) was used to unmix the contribution of NE:

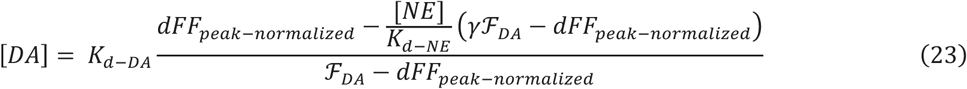

where γ is the ratio of the DA sensor’s saturating responses to NE and DA ligands individually, which can be estimated as

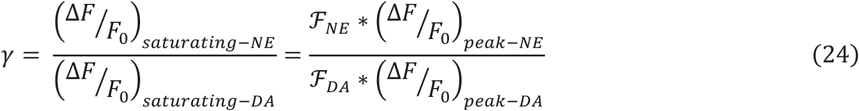

Here, dFF_peak-normalized_ is obtained by scaling the DA sensor signal relative to the 50 µM DA response as described above, K_d-DA_ is the estimated dissociation constant between DA sensor and DA ligand, K_d-NE_ is the estimated dissociation constant between DA sensor and NE ligand, [NE] is the concentration of NE ligand (in this case derived from applying Eq. 22 to the NE sensor, where no crosstalk is assumed in the range of DA doses that we used for calibration), ℱ_NE_ and ℱ_DA_ correspond to the ℱ parameters estimated when fitting Eq. 22 to either NE or DA doses applied to the DA sensor individually, and (ΔF/F_0_)_peak-NE_ and (ΔF/F_0_)_peak-DA_ represent the peak raw ΔF/F_0_ values obtained from the DA sensor for the highest concentration of the respective ligand applied during dipping characterization (these are also the values which the raw signals were normalized by to obtain the inputs for the single-ligand fits using Eq. 22). One remaining caveat of this approach is that a slight amount of crosstalk at the NE sensor is also evident, as can be seen in the NE sensor’s response to 50 µM DA wash-on in **Figures 2B-C,H and S4E-F**; however, this effect was too weak in the range of DA concentrations applied during dipping calibration, so we were unable to calculate dissociation and brightness parameters for the NE sensor in response to DA. With that information, a system of two equations analogous to Eq. 23 could be solved to better estimate both concentrations and dissociate the effects of high-dose NE and DA ligands at the two sensors.

We further used the dipping recordings to assess hydrogel diffusion parameters. To do this, we fitted an exponential rise curve to the first three minutes of each individual sensor cell’s time course in each well of the recording and defined its time constant as the hydrogel diffusion time constant 1-on for that sensor cell at the ligand concentration in the given well (**Figure S3E, bottom**). We performed this analysis both on the raw ΔF/F_0_ signals for each cell as well as the estimated quantitative concentration time course obtained by applying Eq. 22. We subsequently used these 1-on values to perform deconvolution as described above (**Figures S4B-D,G**).

In the case of fiber photometry measurements, the original signals were collected at 50 Hz frequency as described above. A median filter with a 2 second window (every 100 readings) was applied, and the signal was then fitted with an exponentially decaying background correction over tens of minutes of signal preceding sensor activation. Photometry signals were reported in ΔF/F_0_ format normalized to the 15 minutes preceding each IP injection, as in Eq. 20.

Outliers in box plots used throughout the paper are defined as values lying further than 1.5x the interquartile range from the box. All error bars and shading represent ±SEM as defined in figure legends.

## Acknowledgements

We thank all members of the Lehtinen and Andermann labs for fruitful discussions; A. Jang for pilot cell line experiments; C. Sadegh, O. Alturkistani, E. Klein and C. Moore for mouse surgery development and support; R. Fame for CSF collection support; H. Kucukdereli for electronics support; F. Shipley for image analysis support; P. Prasad, M. Webb and B. Grant for mouse colony support; J. Edelhaus, J. Baker, Z. Stolberg, L. Byer and H. Choh for histology support; J. DeBolt and J. Shapiro for administrative support; C. Saper, B. Sabatini, C. Gu, D. Lin, A. Cohen, E. Brown, and G. Yellen for feedback; C. Chen, H. Umemori, C.-H. Chien and the BCH IDDRC Cellular Imaging Core; and M.J. Holtzman for sharing the *FoxJ1-Cre* mouse line. This work was supported by NIH F30 DK131642, T32 HL007901, T32 GM007753, and T32 GM144273 (P.N.K.); an Ellen R. and Melvin J. Gordon Center for the Cure and Treatment of Paralysis Fellowship (C.I.M.); a Walter Benjamin Fellowship of the Deutsche Forschungsgemeinschaft (M.P.); a Lefler Fellowship, Charles A. King Trust Fellowship, and NIH K99 DK134853-02 (S.X.Z.); NIH R01 NS129823-02S1 (J.C.B.); the Bill and Melinda Gates Millennium Scholarship (T.E.L.); a Howard Hughes Medical Institute James H. Gilliam Fellowship and a National Science Foundation Graduate Research Fellowship (Y.C.); NIH T32 NS007473 and F32 NS134588 (A.H.); NIH F32 DK112589 and ZIA DK075169 (A.L.); NIH R01 NS088566, R01 NS129823, RF1 DA048790, a Harvard Brain Science Initiative Bipolar Seed Grant and the New York Stem Cell Foundation (M.K.L.); NIH DP1 AT010971, R01 EY032749, R21 EY035436, Simons Foundation Pilot Award ID 976096, Pew Charitable Trusts Innovation Fund, and a Charles Robert Broderick III Phytocannabinoid Research Grant (M.L.A.); and Boston Children’s Hospital Intellectual and Developmental Disabilities Research Center NIH U54 HD090255, P50 HD105351, and S10 OD016453 to the IDDRC Cellular Imaging Core. The content is solely the responsibility of the authors and does not necessarily represent the official views of the National Institutes of Health.

## Code availability

The custom analysis codes in this work are publicly available at GitHub (https://github.com/pkalugin/kalugin-et-al-morse).

---

**Figure S1.**
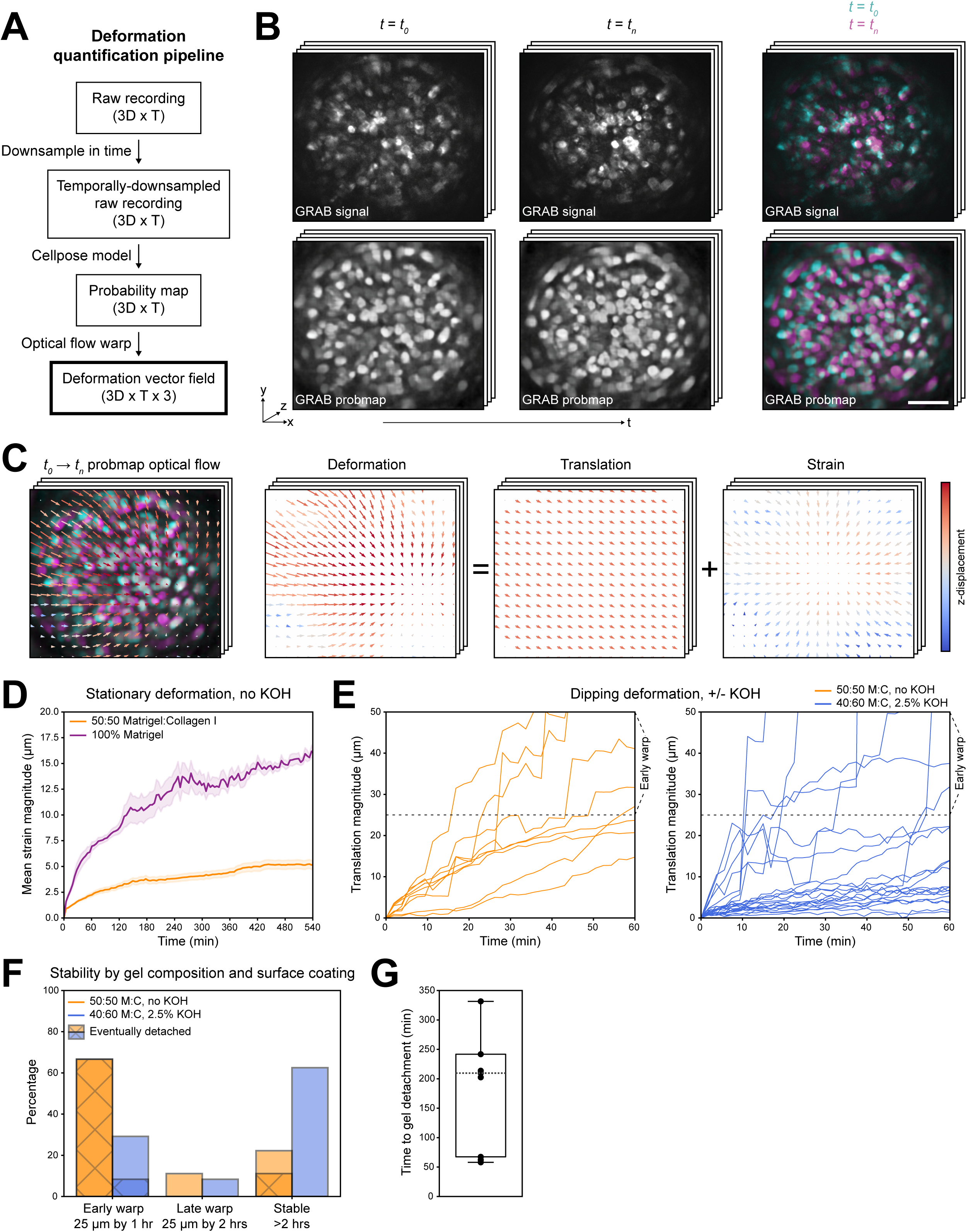
MORSE probe optimization. (**A**) MORSE hydrogel deformation was quantified by applying a Cellpose2.0 model to a temporally-downsampled version of the 3D raw fluorescence recording. The resulting 3D cell probability maps (“probmaps”) estimating each voxel’s likelihood to be part of a cell at each timepoint were registered to a single reference timepoint by calculating optical flow using the imregdemons function in Matlab. (**B**) GRAB sensor fluorescence from an example MORSE recording is shown at two timepoints, *t_0_* and *t_n_*. Cellpose2.0 yields 3D cell probmaps from 3D volumes of fluorescence signal. **Right:** overlays of fluorescence images and probmaps at *t_0_* and *t_n_* highlight hydrogel deformation between the two timepoints, with the probmaps demonstrating notably more uniform signals and better signal-to-noise across the face of the GRIN lens compared to the fluorescence. Scale bar: 100 µm. (**C**) The optical flow pattern calculated on the probmaps from *t_0_* to *t_n_* estimates the pattern of hydrogel deformation between the two timepoints. This deformation vector field can be separated into a 3D translation component (the mean of the vector field across the volume of the hydrogel in the captured volume) and a 3D strain component reflecting swelling/compression, rotation, and other local warping of the hydrogel. The vectors at each point reflect the local deformation estimate, with z-deformation reflected by arrow color; x-y deformation is scaled up by a factor of 2 relative to the dimensions of the image for easier visualization. (**D**) Quantification of warping in static hydrogels of different recipes. MORSE probes were prepared without lens surface coating using hydrogels consisting of pure Matrigel and of a 50:50 mixture of Matrigel and collagen I (n=6 probes per recipe). Probes were secured vertically in PBS, sealed to minimize fluid evaporation, and imaged continuously overnight for 9 hours. Deformation is reported as local strain vector magnitude averaged across the hydrogel volume. Shading depicts SEM across probes. (**E**) Quantification of detachment in dipping hydrogels of different recipes. MORSE probes were prepared with or without 2.5% KOH lens surface coating using hydrogels consisting of 50:50 Matrigel:collagen I (n=9) and 40:60 Matrigel:collagen I (n=24), respectively. Probes were dipped for 5 min at a time over 2-7 hours into sequential wells of ligand solutions formulated in PBS, as described in Figure 1D. Detachment is reported as mean 3D translation of the hydrogel volume. Horizontal lines at 25 µm translation indicate the major warp threshold, corresponding to around one cell diameter’s displacement. (**F**) Summary of the detachment time courses shown in **E**. Probes are subdivided into early warp (>25 µm translation in the first hour of recording), late warp (>25 µm translation in the second hour), and stable (<25 µm translation in the first two hours). Of the 50:50 Matrigel:collagen I probes prepared without KOH surface treatment, n=7/9 eventually demonstrated complete hydrogel detachment, with n=6/7 of those also falling into the early warp category. Only n=2/9 (22%) were stable over 2 hours, and one of those hydrogels eventually detached. Of the 40:60 Matrigel:collagen I probes prepared with 2.5% KOH surface treatment, n=7/24 fell into the early warp category, with n=2/7 of those detaching completely. Meanwhile, n=15/24 (63%) were stable over 2 hours. (**G**) Summary of the times to hydrogel detachment in the n=9 detached probes of both recipes shown in **E-F**.

**Figure S2.**
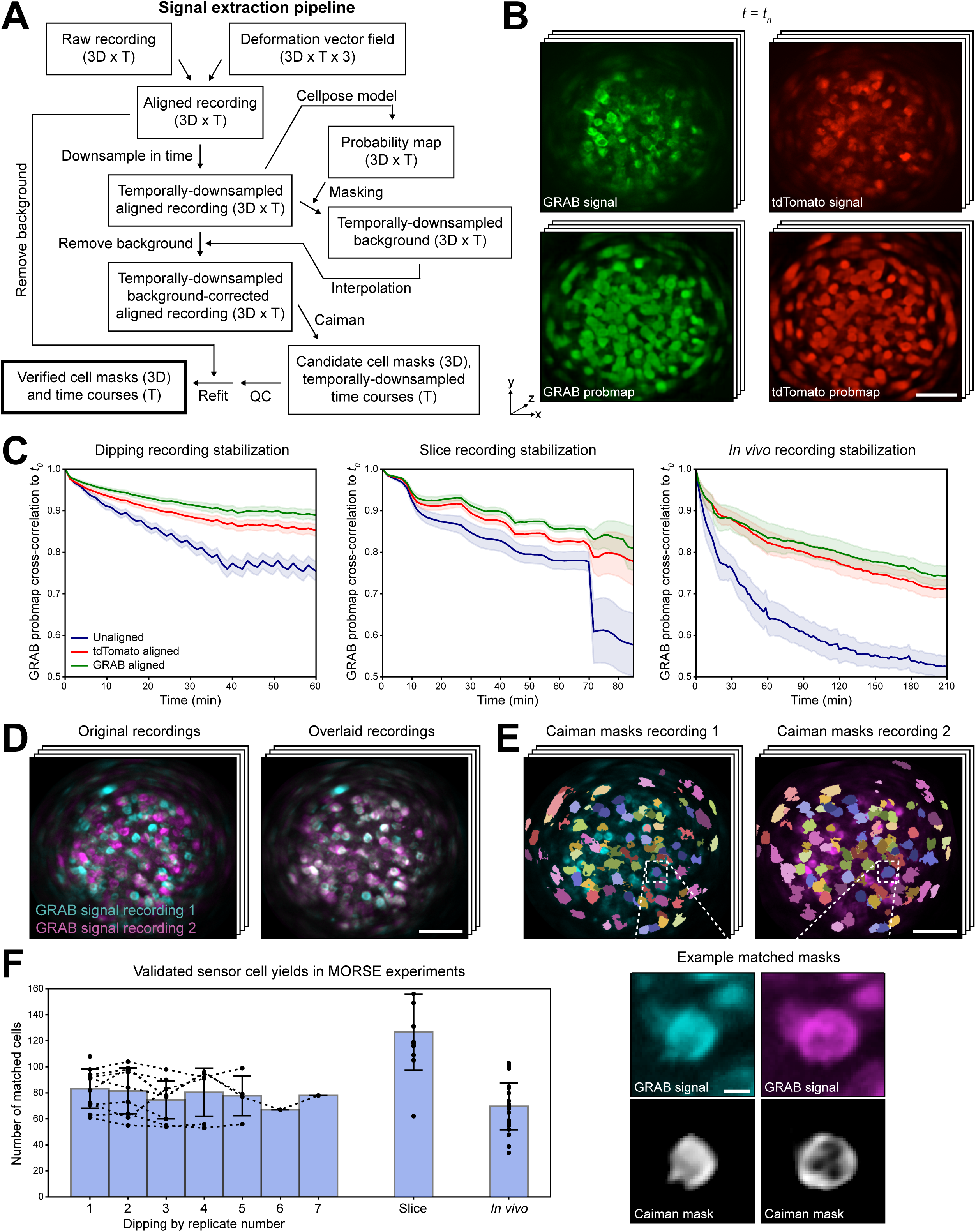
MORSE 3D alignment and signal extraction. (**A**) Residual hydrogel deformation in MORSE probes of optimal recipe was stabilized by applying the deformation vector field calculated as in **Figures S1A-C** to the raw fluorescence recording. This aligned fluorescence signal was temporally downsampled and background fluorescence across the hydrogel was estimated by recalculating Cellpose2.0 probmaps on this signal, using the probmaps to mask out voxels belonging to cells, and spatially interpolating across the remaining non-cell voxels. The background-corrected temporally-downsampled fluorescence was then processed with the Caiman CNMF algorithm to generate candidate 3D single cell masks and associated signal time courses. These candidate masks underwent manual quality control (QC) and were subsequently refitted on the original aligned fluorescence signal at full sampling frequency, yielding final verified cell masks and time courses. (**B**) A subset of MORSE probes was prepared with 10% of the sensor cells expressing cytosolic tdTomato, a signal that is inert to extracellular ligand concentration changes, in addition to the GRAB sensors. Two-color imaging of these probes was performed, and Cellpose2.0 was used to produce one set of probmaps from each color channel, which were subsequently used for image alignment. (**C**) Quantification of image stabilization in dipping (n=36 recordings across n=11 probes), slice (n=11), and *in vivo* CSF (n=11) MORSE recordings using tdTomato and GRAB sensor fluorescence-derived probmaps for deformation calculation. Image stability is reported as normalized cross-correlation of the GRAB fluorescence probmap at each timepoint compared to the start of the recording. Shading depicts SEM across recordings. Both GRAB and tdTomato-based alignments substantially improve image stability in all three types of experiments, with the GRAB signal performing as well as or slightly better than the tdTomato. This is likely due to better spatial coverage achieved by the larger numbers of cells expressing GRAB sensors on the probes – up to 50% of cells depending on transfection efficiency (not shown), compared to 10% of cells for tdTomato. Notably, n=5/11 of the slice recordings involved the lifting of the probe off the slice at around the 70 min timepoint, causing a substantial deformation in the gel that alignment using both color channels was able to correct. (**D**) Representative GRAB fluorescence signal volumes from two dipping recordings of the same MORSE probe (**left**) are aligned using manually defined landmarks (**right**). Scale bar: 100 µm. (**E**) Following alignment of representative fluorescence volumes, cell masks calculated on the two recordings of the same MORSE probe are overlaid and manually curated to ensure correct matches. **Top:** masks matched across two dipping recordings shown in **D** are overlaid on their respective representative fluorescence volumes, with color-coding reflecting matches. Scale bar: 100 µm. **Bottom:** inset shows the GRAB signal distributions and corresponding masks for a single example cell matched across both recordings. Note how the masks reflect the changing shape of the cell over time, with mask weights preferentially distributed on the cell membrane, the site of GRAB sensor expression. Scale bar: 10 µm. (**F**) Summary of matched cell mask numbers across dipping (n=50 recordings across n=10 probes), slice (n=11 recordings with paired dipping characterizations), and *in vivo* CSF (n=22 recordings with paired dipping characterizations) MORSE recordings. In the case of the dipping recordings, each probe was dipped up to six additional times for 30-90 min at a time following the initial recording and paired dipping characterization; all additional recordings were matched to the initial recording, and data points corresponding to an individual probe are connected across replicates by dotted lines. In the case of slice recordings, the yield is likely higher due to a combination of high hydrogel stability, allowing for better recovery of cells with low signal-to-noise, and the fact that these probes expressed seven robustly active GRAB sensors, while the dipping probes expressed 16 sensors, only ten of which were reliably active and recovered by the Caiman algorithm. In case of the *in vivo* probes, the yield is likely lower than in slice due to accumulated deformation across long recordings of 3-4 hours in the mouse, as shown in **C**, prior to dipping characterization. Error bars: ±SEM across probes.

**Figure S3.**
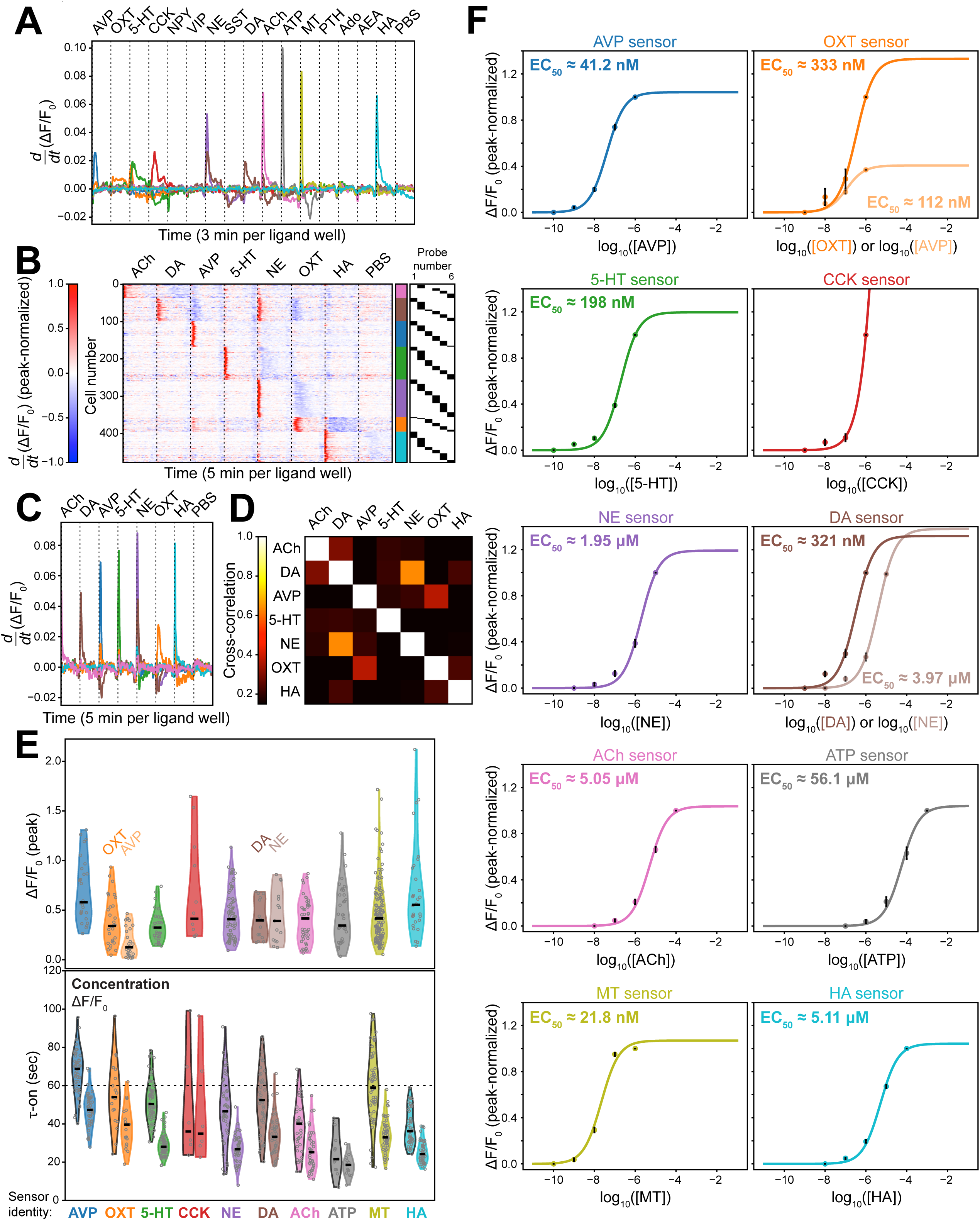
MORSE cell clustering, hydrogel diffusion assessment, and dose-response relationships. (**A**) Mean time-derivative traces for each sensor group in Fig. 1E (**bottom**). Plotted colors correspond to the sensor names labeled below in **E**. (**B**) 478 single-cell ROIs were recovered from n=6 MORSE probes (independent set from those depicted in Fig. 1E) carrying 7 GRAB sensor cell types and dipped into high concentrations of their 7 respective ligands: acetylcholine (ACh, 1 mM), dopamine (DA, 1 μM), vasopressin (AVP, 1 μM), serotonin (5-HT, 1 μM), norepinephrine (NE, 10 μM), oxytocin (OXT, 1 μM), and histamine (HA, 1 mM), followed by a wash well of phosphate-buffered saline (PBS). Each probe was dipped into a 20 μL volume of each ligand for 5 minutes prior to movement into the next well. Heatmap depicts peak-normalized time-derivatives of the single-cell signals. The identity of the probe that each cell is derived from is depicted in the black raster plot on the far right, and the sensor identity of the cell is shown with the color bar corresponding to the sensor names in **E**. The cells from all probes are listed in order of activation during the recording, demonstrating robust activation of up to all 7/7 sensors by their respective ligands (median of 7 active sensors per probe, interquartile range 6-7; average of 12.0 cells per sensor per probe). (**C**) Mean time-derivative traces for each sensor group in **B**. (**D**) Cross-correlation of the mean time-derivative traces for each sensor group in **C**. Robust crosstalk between the NE and DA sensors, along with weaker crosstalk between AVP and OXT sensors, is redemonstrated as in Fig. 1E**-F**. (**E**) Summary of sensor response characteristics for the ten GRAB sensors demonstrating robust activation in the initial set of ten MORSE probes depicted in Fig. 1E**-F**. **Top:** summary of peak ΔF/F_0_ signals for each cell expressing each of the ten activated sensors in response to high concentrations of their respective ligands: 1 μM AVP, 1 μM OXT, 1 μM 5-HT, 1 μM CCK, 10 μM NE, 1 μM DA, 1 mM ACh, 1 mM ATP, 10 μM MT, 1 mM HA. Each cell’s peak ΔF/F_0_ signal is plotted as a gray point, with their distribution shown in the violin plot and a solid black line depicting the median value for each sensor type. Sensor types are color-coded below. For OXT and DA sensors, both their on-target (dark violin plot) and off-target activation (pale violin plot) by AVP and NE, respectively, are shown. **Bottom:** each MORSE probe was then dipped into two sets of wells containing increasing concentrations of 8 ligands at a time extending in 10-fold increments across a dynamic range of 100,000-fold. The time course of each sensor’s ΔF/F_0_ response to dipping into the lowest concentration of its corresponding ligand above the sensor’s estimated EC_50_ value was analyzed. For each sensor type, the τ-on value for each cell’s ΔF/F_0_ response to this ligand concentration is plotted on the right and summarized with the violin plot with a gray border. Additionally, using the determined sensor sensitivity characteristics, the time course of each sensor’s ligand concentration estimate was calculated from the ΔF/F_0_ response. The τ-on value for each cell’s concentration estimate to this ligand concentration is plotted on the left and summarized with the violin plot with a black border. Due to the nonlinear transformation from ΔF/F_0_ signal to concentration estimate and the choice of a concentration above the EC_50_ for each sensor, all τ-on values for the concentration estimates are larger than their respective ΔF/F_0_ response τ-on values. Almost all sensors demonstrate a τ-on under 1 minute, reflecting rapid diffusion of all ten ligands through the hydrogel to reach their sensors. (**F**) Dose-response curves of each of ten GRAB sensors responding to their corresponding ligands at 10-fold concentration increments. Nine of the ten sensors (all except CCK) exhibited sufficient trends toward a saturation plateau by the highest concentration applied. EC_50_ values for these successfully characterized sensors ranged from the tens of nanomolar to the tens of micromolar. In the case of the OXT and DA sensors, which demonstrated strong crosstalk in the tested concentration ranges from the AVP and NE ligands, respectively, these off-target EC_50_ values are also quantified. Plots demonstrate ΔF/F_0_ signals peak-normalized to the signal achieved in each cell by the maximal administered ligand concentration, averaged across all cells corresponding to each sensor at each ligand concentration. Error bars depict ±SEM across sensor cells and solid lines are the estimated sigmoid response curves.

**Figure S4.**
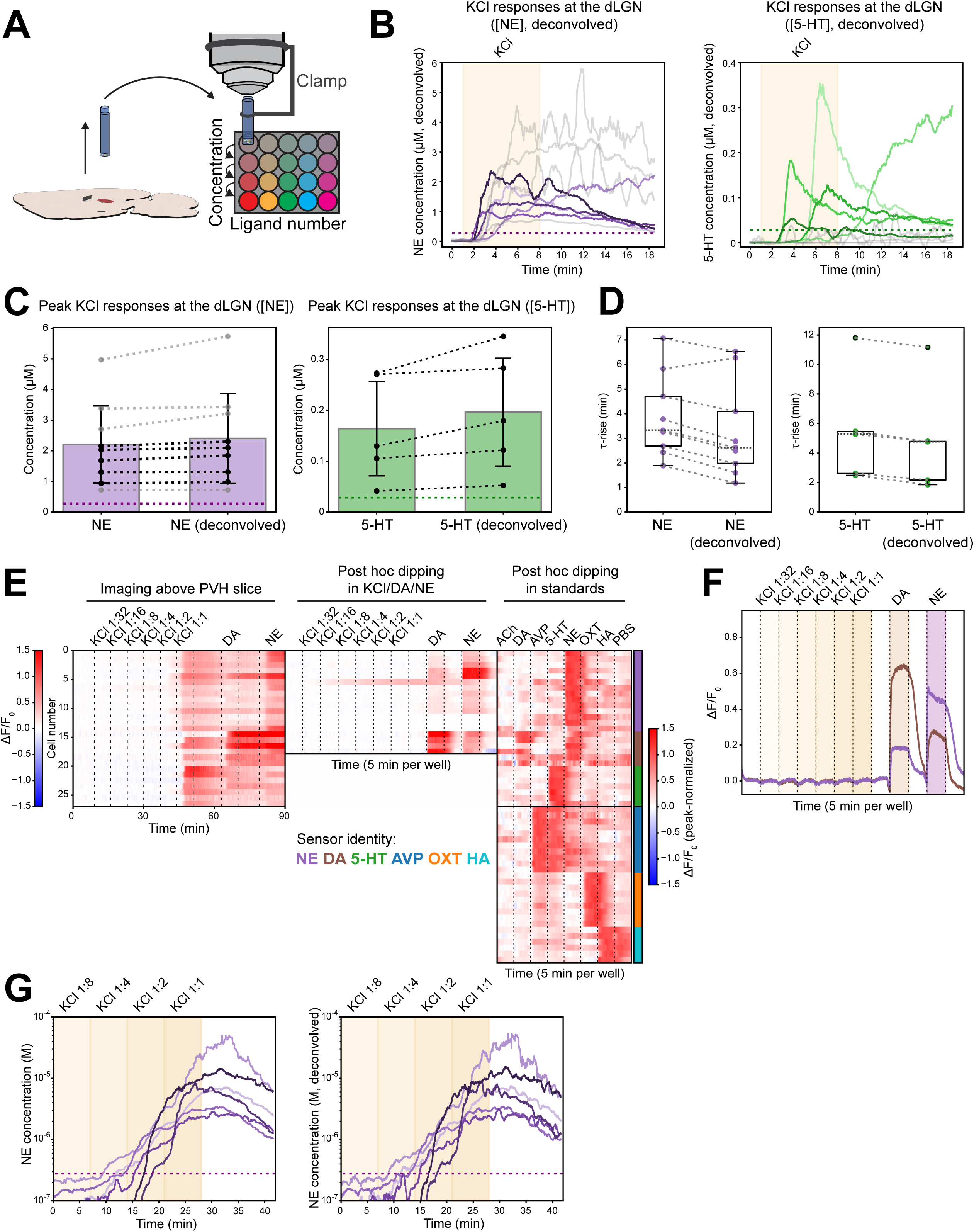
Additional quantifications of MORSE recordings from live brain slices. (**A**) Schematic of MORSE probe liftoff from the dLGN slice and subsequent dipping characterization. (**B**) NE (**left**) and 5-HT (**right**) concentrations measured by n=9 MORSE probes positioned over the dLGN during KCl administration, deconvolved using the hydrogel transfer functions for each sensor with τ values as summarized in **Fig. S3E** (original traces in Fig. 2D). As in Fig. 2D, a threshold peak-normalized ΔF/F_0_ signal of 0.125 was set to distinguish strong responses, which were present in both sensors in n=5/9 recordings. These are plotted with matching hue depths for comparison (e.g. the darkest purple NE curve corresponds to the darkest green 5-HT curve), while the remaining n=4/9 recordings with only robust NE responses plotted in gray. Colored dotted lines on both plots demonstrate the peak-normalized ΔF/F_0_ = 0.125 thresholds (corresponding to 0.278 µM for NE and 28.3 nM for 5-HT). (**C**) Comparison of the peak NE (**left**) and 5-HT (**left**) concentrations elicited by KCl administration in **Figs. 2D, S4B** as calculated with and without deconvolution of the hydrogel transfer functions. Values from the same recording and for the same ligand calculated with each of the two methods are connected by dotted lines, and the n=4/9 recordings demonstrating robust release of NE but not 5-HT are plotted in gray. Dotted lines on both sets of colored bars correspond to the peak-normalized ΔF/F_0_ = 0.125 thresholds (0.278 µM for NE and 28.3 nM for 5-HT). Deconvolution increases the average peak NE concentration by 193 nM/8.7% and the average peak 5-HT concentration by 32.3 nM/19.7%. Error bars: ±SEM across recordings. (**D**) Comparison of the τ-rise onset timings of the NE (**left**) and DA (**right**) concentration time courses elicited by KCl administration in **Figs. 2D, S4B** as calculated with and without deconvolution of the hydrogel transfer functions. Values from the same recording and for the same ligand calculated with each of the two methods are connected by dotted lines. Deconvolution reduces the average NE concentration τ-rise by 35.4 sec/15.2% and the average 5-HT concentration τ-rise by 34.9 sec/10.5%. (**E**) Single-sensor cell traces from one seven-sensor (ACh, AVP, OXT, 5-HT, NE, DA, HA) MORSE probe recorded in the PVH (left heatmap, ΔF/F_0_ depicted as per left color bar) and subsequently dipped first into a dose range of KCl concentrations followed by saturating doses of DA and NE (middle heatmap, ΔF/F_0_ depicted as per left color bar) and then separately into standard solutions (right heatmap, peak-normalized ΔF/F_0_ depicted as per right color bar). The slice was continuously perfused with warm aCSF, which was periodically replaced with aCSF containing a dilution range of KCl concentrations (1:32, 1:16, 1:8, 1:4, and 1:2 dilutions of a 50 mM stock of KCl in aCSF as well as the full stock concentration labeled 1:1), DA (50 μM), and NE (50 μM) at the indicated times. During the first dipping recording (middle heatmap), the probe was dipped into these same solutions, with blank spaces indicating wells containing aCSF. During the second dipping recording (right heatmap), the probe was dipped into standard solutions of the seven ligands corresponding to each sensor formulated in PBS (1 mM ACh, 1 μM DA, 1 μM AVP, 1 μM 5-HT, 10 μM NE, 1 μM OXT, 1 mM HA). Cell matches are performed across all three recordings, with blank rows in the middle heatmap indicating cells that were only matched between the first and third recordings but not identified in the second. The upper half of the right heatmap demonstrates the sensor cells with ROIs successfully matched between the slice recording and the second dipping recording, and the lower half shows additional cells that were inactive during the slice recording as well as the first dipping recording but responded in the second dipping recording. The sensor identity of each cell is shown with the color bar on the right as per legend. (**F**) Mean ΔF/F_0_ traces for NE and DA sensors from the middle heatmap in **D**, with dipping timings indicated by vertical dotted lines and colored shading, and sensors color-coded as per legend in **D**. Neither sensor is driven by dipping into any KCl solution, indicating that the KCl responses on the left heatmap in **D** correspond to KCl-induced ligand release from the PVH slice rather than direct sensor response to KCl. On the other hand, both sensors continue to demonstrate responses to their respective ligands (as well as crosstalk at both sensors, with both ligands administered at very high concentrations of 50 µM). (**G**) NE concentrations measured by n=6 MORSE probes at the PVH during administration of escalating doses of KCl, corresponding to the ΔF/F_0_ signals in Fig. 2I**, left**, before (**top**) and after (**bottom**) deconvolution using the hydrogel transfer function for the sensor. Colored dotted lines on both plots correspond to the peak-normalized ΔF/F_0_ = 0.125 threshold (0.278 µM for NE).

**Figure S5.**
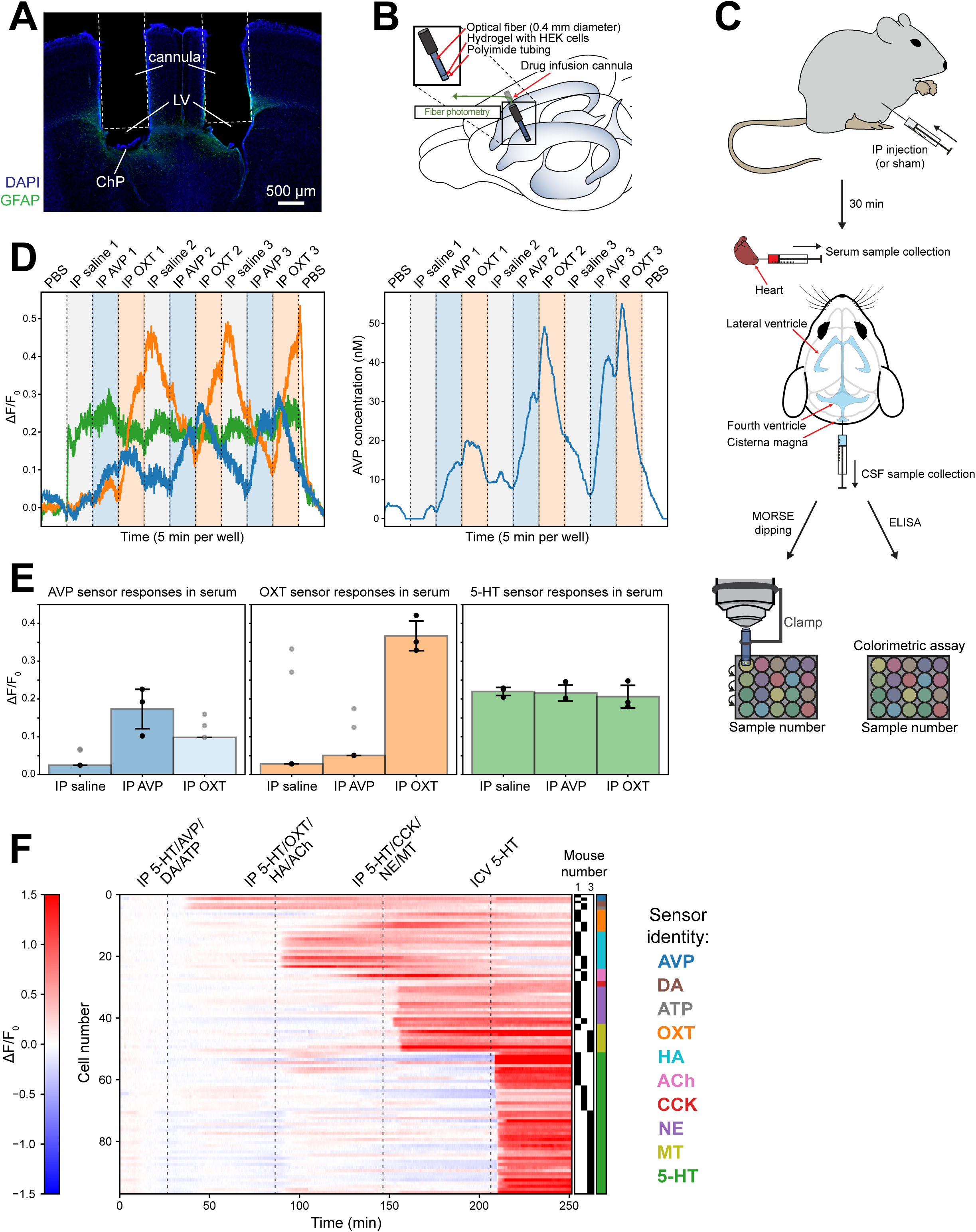
Ten-sensor MORSE recordings from mouse serum *in vitro* following IP injections of AVP and OXT and *in vivo* in the lateral ventricle CSF of awake mice. (**A**) Histology from a representative mouse with bilateral anterior lateral ventricle cannulae, stained for glial fibrillary acidic protein (GFAP, green) to visualize gliosis. White dotted lines delineate the cannula tracks inserted into the lateral ventricles (LV), containing choroid plexus tissue (ChP). No GFAP signal asymmetry between left and right hemispheres induced by MORSE probe implantation into one of the cannulae was noted in any animals across the span of the ventricular system. (**B**) Schematic of the mouse lateral ventricle cannulation setup as in Fig. 3A, this time used to record from fiber photometry probes. These probes were prepared analogously to the MORSE probes, except that a 400 µm diameter optical fiber was used instead of a 500 µm diameter GRIN lens, and HEK293T cells were prepared with a single GRAB sensor type. Before a recording, the mouse was head-fixed on a running wheel and one of the cannulae was acutely opened to expose the CSF, at which point the fiber photometry probe was inserted hydrogel-first into the cannula and sealed into place with RTV silicone. Following several hours of imaging, the probe was removed from the animal. (**C**) Schematic of the *in vitro* CSF analysis experiments. **Top:** naïve mice received IP injections of AVP, OXT (both at 1 mg/kg in 200 µL of normal saline), or normal saline (200 µL), or sham abdominal needle pokes without injection. **Middle:** 30 minutes following these treatments, the mice were sacrificed. Transcardiac serum collections (above) and cisterna magna CSF collections (below) were performed. **Bottom:** the fresh CSF and serum samples were then split apart for concurrent measurement of AVP, OXT, and 5-HT composition by both ELISA colorimetric assay and *in vitro* dipping of MORSE probes into the samples with two-photon imaging. (**D**) A MORSE probe carrying ten GRAB sensors (AVP, OXT, 5-HT, CCK, NE, DA, ACh, ATP, MT, HA) was sequentially dipped *in vitro* into 20 µL serum samples from mice that received IP saline, AVP, or OXT injections (n=3 mice each), as well as wash wells containing plain PBS at the start and end of the recording. **Left:** ΔF/F_0_ responses in AVP, OXT, and 5-HT sensor cells on this probe as it was dipped sequentially into the samples labeled above. Sensors are color-coded as per legend in **E**. **Right:** estimated AVP concentration trace calculated from the ΔF/F_0_ trace on the left, color-coded in the same fashion. (**E**) Peak ΔF/F_0_ readings attained for each sensor in each sample dip demonstrated in **A, left**. Each individual sample reading is plotted with a dot, and mean readings across samples are shown with colored bars. Due to incomplete ligand washout between some adjacent wells, there is some signal contamination. Readings likely contaminated by an incompletely washed out signal from the immediately preceding sample are plotted in gray and not included in the mean ΔF/F_0_ readings. In particular, AVP sensor readings from IP OXT serum samples are contaminated in all three evaluated samples, so the entire mean bar has pale shading. Error bars depict SEM across samples of each type. Overall, it is evident that high AVP is only detected in serum from mice that received IP AVP, high OXT is only detected in serum from mice that received IP OXT, and 5-HT is detected in all serum samples (but not the PBS wash wells), reflecting baseline endogenous serotonin levels in serum. (**F**) MORSE recordings were performed from awake mouse CSF with the setup as depicted in Fig. 3A. Mice were outfitted with MORSE probes carrying ten GRAB sensors (AVP, OXT, 5-HT, CCK, NE, DA, ACh, ATP, MT, HA) and received three IP injections containing mixtures of four ligands at a time (5-HT/AVP/DA/ATP at doses of 3 mg/kg, 1 mg/kg, 1 mg/kg, and 50 mg/kg respectively, 5-HT/OXT/HA/ACh at doses of 3 mg/kg, 1 mg/kg, 10 mg/kg, and 10 mg/kg respectively, and 5-HT/CCK/NE/MT at doses of 3 mg/kg, 1 mg/kg, 1 mg/kg, and 10 mg/kg respectively), followed by an ICV injection of 5-HT (5 nmol/µL in aCSF infused at 5 µL/min over 6 min). The heatmap depicts ΔF/F_0_ signals for each individual cell from such recordings in n=3 different mice. The identity of the mouse that each cell is derived from is depicted in the black raster plot on the far right, and the sensor identity of the cell is shown with the color bar corresponding to the sensor names in the legend on the right. Up to 9/10 sensors were successfully recorded in a single mouse (median of 8 active sensors per probe, average of 5.1 cells per sensor per probe). All ligands reached the CSF within minutes of their respective IP injections, with clear differences in kinetics (for example, much more rapid appearance of HA in the CSF compared to concurrently delivered OXT and ACh at the second IP injection).

## Supplementary Video Legends

All Supplementary Videos can be downloaded in a single zipped folder here: https://www.dropbox.com/scl/fi/4ainp9mptgpokoya6fpe7/allvideos.zip?rlkey=nz5dvo4dvs3hmfgy03iemdkt4&st=6g08X305&dl=0

**Supplementary Video 1. MORSE hydrogel optimization.** https://www.dropbox.com/scl/fi/uaujqowl3loweunqj3lps/sv1.avi?rlkey=52vba8bp97xvpvk3jug72fea1&st=upv7j3no&dl=0

Continuous volumetric two-photon imaging in a single well of PBS from MORSE probes prepared with a hydrogel composed of pure Matrigel (**left**) or a 50-50 mixture of Matrigel and collagen I (**right**), containing sensor cells expressing various GRAB sensors. The raw GRAB sensor fluorescence (**top**) and the associated Cellpose2.0 probability map (**bottom**) are shown. Images at each timepoint demonstrate the mean raw signal (no normalization) of ten consecutive imaging volumes (1.9 sec/vol) collected every 4.7 min, averaged over the full thickness of the gel (20 z-planes, ∼180 µm thickness). Time is shown as hh:mm:ss, scale bar: 100 µm. See **Figure S1D** for additional details.

**Supplementary Video 2. MORSE robotic dipping.** https://www.dropbox.com/scl/fi/s3w3ctke5r3whvuncbymr/sv2.avi?rlkey=vk8z3zs5odhsv5rmuv4q530m7&st=pilcrbyd&dl=0

A MORSE probe was secured on the two-photon microscope with a custom lens clamp, centered under the objective and parallel to the objective axis, with the hydrogel facing downward and the distance of the probe from the objective fixed to place the middle layers of the hydrogel in the image focus. The probe was then positioned above a 384-well plate carrying ∼20 µL wells of solutions carrying various biological fluid samples or ligands at different concentrations. Using the robotic controls of the microscope, the probe was subsequently dipped for several minutes at a time into one well of the plate after another, with 3D volumetric two-photon imaging of the hydrogel performed throughout using optotune focus tuning. The dwell time in each well is shortened in this video for demonstration purposes, but the probe transfer time between wells of ∼22 sec is consistent for all dipping recordings. Time is shown as hh:mm:ss. See Figure 1D for additional details.

**Supplementary Video 3. MORSE silicone surface coating optimization.** https://www.dropbox.com/scl/fi/oz6nl6bau90nbiy9zrcvl/sv3.avi?rlkey=gfbuwztxdk2qg3u4X8hgh1u25&st=w4d561jb&dl=0

MORSE probes carrying six GRAB sensor cell types were prepared either without RTV silicone surface coating or with 2.5% KOH coating for 1 hour, and with either 50:50 (**left**), 60:40 (**middle**), or 40:60 (**right**) ratios of Matrigel and collagen I as indicated. Dipping characterization was performed, with probes dipped sequentially into up to 38 wells (5 minutes per well) containing PBS or near-saturating sensor ligand concentrations in PBS: serotonin (5-HT, 1 µM), vasopressin (AVP, 1 µM), oxytocin (OXT, 1 µM), cholecystokinin (CCK, 1 µM), neuropeptide Y (NPY, 1 µM), vasoactive intestinal peptide (VIP, 1 µM). Images at each timepoint demonstrate the mean raw GRAB sensor fluorescence signal (no normalization) of ten consecutive imaging volumes (1.9 sec/vol) averaged over the full thickness of the gel (20 z-planes, ∼180 µm thickness). As can be appreciated at several ligand transition points in this video, dipping fluorescence signals often demonstrated a sharp upward jump at the start of a new well of fluid. The magnitude of this jump appears to depend on the starting brightness of the cell and likely reflects the effects of forces applied to the hydrogel as it crosses the air-fluid interface. This artifact was fitted with an exponential decay (which exhibited a consistent decay constant of ∼30 sec), scaled by the jump magnitude between adjacent wells for subsequent analyses,. Composition of the solution in each well is shown in the lower left. Time is shown as hh:mm:ss, scale bar: 100 µm. See **Figures S1E-G** for additional details.

**Supplementary Video 4. Stabilization of a dipping 16-sensor MORSE probe.** https://www.dropbox.com/scl/fi/cuhjkrb4fknmqltw66p32/sv4.avi?rlkey=5f0k5kt7pzcafqewbjp9twk3c&st=03runeq9&dl=0

A representative MORSE probe carrying 16 GRAB sensor cell types is shown deforming slowly over tens of minutes as it is dipped into various ligand solutions in PBS (**left**). A Cellpose2.0 model was applied to the raw GRAB sensor or inert tdTomato (not shown) signals to calculate cell probability maps over time, and these probability maps were used to quantify the hydrogel deformation (see **Figures S1A-C**). Image stabilization achieved using the tdTomato signal probability maps (**middle**) and the GRAB signal probability maps (**right**) is shown. This probe was first dipped into two sets of mixtures, each containing increasing concentrations of 8 ligands in PBS. The doses constituted several dilutions of an initial stock solution containing a combined mixture of ligands as follows:

Mix 1 undiluted stock: vasopressin (AVP, 1 µM), oxytocin (OXT, 1 µM), serotonin (5-HT, 1 µM), cholecystokinin (CCK, 1 µM), neuropeptide Y (NPY, 1 µM), vasoactive intestinal peptide (VIP, 1 µM), norepinephrine (NE, 10 µM), and somatostatin (SST, 1 µM) Mix 2 undiluted stock: dopamine (DA, 1 µM), acetylcholine (ACh, 1 mM), adenosine triphosphate (ATP, 1 mM), melatonin (MT, 10 µM), parathyroid hormone (PTH, 1 µM), adenosine (Ado, 1 mM), anandamide (AEA, 1 mM), and histamine (HA, 1 mM)

The dilutions for each mixture were applied in increasing order spaced by multiples of ten: 10^5^-, 10^4^-, 10^3^-, 10^2^-, and 10-fold dilutions of the stock followed by the undiluted stock. Next, the probe was sequentially dipped into mixtures containing one of three sets of ligands (5-HT/NE/DA, VIP/AVP/Ado/HA, ATP/SST/AEA/PTH) combined in combinations of various concentrations. Each dip lasted 3 minutes. Images at each timepoint demonstrate the mean raw GRAB sensor fluorescence signal (no normalization) of three consecutive imaging volumes (1.9 sec/vol) averaged over six adjacent image planes spanning a total of ∼55 µm in gel thickness (∼30% of total thickness containing sensor cells). Composition of the solution in each well is shown in the lower left. Time is shown as hh:mm:ss, scale bar: 100 µm. See **Figures S2A-C** for additional details.

**Supplementary Video 5. Identification dipping of a 16-sensor MORSE probe.** https://www.dropbox.com/scl/fi/g4dxmjr9701ojli1v6h8i/sv5.avi?rlkey=9a17wrionhlurv7vsb7gh1412&st=vplijule&dl=0

A representative MORSE probe carrying 16 GRAB sensor cell types is shown being dipped into solutions containing near-saturating concentrations of the 16 respective ligands in PBS: vasopressin (AVP, 1 µM), oxytocin (OXT, 1 µM), serotonin (5-HT, 1 µM), cholecystokinin (CCK, 1 µM), neuropeptide Y (NPY, 1 µM), vasoactive intestinal peptide (VIP, 1 µM), norepinephrine (NE, 10 µM), somatostatin (SST, 1 µM), dopamine (DA, 1 µM), acetylcholine (ACh, 1 mM), adenosine triphosphate (ATP, 1 mM), melatonin (MT, 10 µM), parathyroid hormone (PTH, 1 µM), adenosine (Ado, 1 mM), anandamide (AEA, 1 mM), and histamine (HA, 1 mM). Each dip lasted 3 minutes. Images at each timepoint demonstrate the mean raw GRAB sensor fluorescence signal (no normalization) of three consecutive imaging volumes (1.9 sec/vol) averaged over three adjacent image planes spanning a total of ∼30 µm in gel thickness (∼15% of total thickness containing sensor cells) (**left**) and the corresponding ΔF/F_0_ signal (**right**). Composition of the solution in each well is shown in the lower left. Time is shown as hh:mm:ss, scale bar: 100 µm. See Figures 1E-F **and S3A** for additional details. The same MORSE probe is shown dipping into escalating ligand doses in **Supplementary Video 6**.

**Supplementary Video 6. Dose-response dipping of a 16-sensor MORSE probe.** https://www.dropbox.com/scl/fi/hx7X0wa1bwdvvkzpkqz8g/sv6.avi?rlkey=jc5p9avczwlifu0bbj96wkf6p&st=xxyx52pd&dl=0

The same MORSE probe as in **Supplementary Video 5**, with the image rotated and warped for alignment, is shown being dipped into two sets of mixtures, each containing increasing concentrations of 8 ligands in PBS. The doses constituted several dilutions of an initial stock solution containing a combined mixture of ligands as follows:

Mix 1 undiluted stock: vasopressin (AVP, 1 µM), oxytocin (OXT, 1 µM), serotonin (5-HT, 1 µM), cholecystokinin (CCK, 1 µM), neuropeptide Y (NPY, 1 µM), vasoactive intestinal peptide (VIP, 1 µM), norepinephrine (NE, 10 µM), and somatostatin (SST, 1 µM) Mix 2 undiluted stock: dopamine (DA, 1 µM), acetylcholine (ACh, 1 mM), adenosine triphosphate (ATP, 1 mM), melatonin (MT, 10 µM), parathyroid hormone (PTH, 1 µM), adenosine (Ado, 1 mM), anandamide (AEA, 1 mM), and histamine (HA, 1 mM)

The dilutions for each mixture were applied in increasing order spaced by multiples of ten: 10^5^-, 10^4^-, 10^3^-, 10^2^-, and 10-fold dilutions of the stock followed by the undiluted stock. Each dip lasted 3 minutes. Images at each timepoint demonstrate the mean raw GRAB sensor fluorescence signal (no normalization) of three consecutive imaging volumes (1.9 sec/vol) averaged over three adjacent image planes spanning a total of ∼30 µm in gel thickness (∼15% of total thickness containing sensor cells) (**left**) and the corresponding ΔF/F_0_ signal (**right**). Composition of the solution in each well is shown in the lower left. Time is shown as hh:mm:ss, scale bar: 100 µm. See Figures 1G **and S3E-F** for additional details.

**Supplementary Video 7. MORSE recording from the dorsolateral geniculate nucleus (dLGN) of the thalamus in a live brain slice with a seven-sensor probe.** https://www.dropbox.com/scl/fi/goyxk7duok38xoarwqiab/sv7.avi?rlkey=xxvu1991k4h0xzktuajdabqhy&st=jpdn1xy9&dl=0

A representative MORSE probe carrying seven GRAB sensor cell types is recorded while pressed against the dLGN region of a live brain slice. The slice was continuously perfused with warm aCSF, which was periodically replaced with aCSF containing KCl (50 mM), DA (50 µM), 5-HT (50 µM), and NE (50 µM). Images at each timepoint demonstrate the mean raw GRAB sensor fluorescence signal (no normalization) of three consecutive imaging volumes (1.9 sec/vol) averaged over three adjacent image planes spanning a total of ∼30 µm in gel thickness (∼15% of total thickness containing sensor cells) (**left**) and the corresponding ΔF/F_0_ signal (**right**). White arrowheads indicate example cells carrying DA, 5-HT, and NE sensors (same cells as in **Supplementary Video 8**), and individual active cells from this recording are plotted in Figure 2B**, left heatmap**. Composition of the wash-on solution is shown in the lower left. Time is shown as hh:mm:ss, scale bar: 100 µm. See Figures 2 **and S4** for additional details. The same MORSE probe is shown undergoing post hoc dipping characterization in **Supplementary Video 8**.

**Supplementary Video 8. Post hoc dipping of a seven-sensor MORSE probe following a slice recording over the dLGN.** https://www.dropbox.com/scl/fi/10kx3ia5m0bbn5lvbhtoi/sv8.avi?rlkey=5f3f0ueggt46vh7wrao18ob6d&st=hq6n8b4s&dl=0

The same MORSE probe as in **Supplementary Video 7**, with the image rotated and warped for alignment, is shown being dipped into standard solutions of seven ligands. The first 8 wells contain the same solutions as washed onto the slice – dips into aCSF alone, interspersed with dips into aCSF containing KCl (50 mM), DA (50 µM), 5-HT (50 µM), or NE (50 µM). Next, the probe was dipped into solutions containing near-saturating concentrations of seven ligands in PBS: acetylcholine (ACh, 1 mM), dopamine (DA, 1 µM), vasopressin (AVP, 1 µM), serotonin (5-HT, 1 µM), norepinephrine (NE, 10 µM), oxytocin (OXT, 1 µM), and histamine (HA, 1 mM). Each dip lasted 5 minutes. Images at each timepoint demonstrate the mean raw GRAB sensor fluorescence signal (no normalization) of three consecutive imaging volumes (1.9 sec/vol) averaged over three adjacent image planes spanning a total of ∼30 µm in gel thickness (∼15% of total thickness containing sensor cells) (**left**) and the corresponding ΔF/F_0_ signal (**right**). White arrowheads indicate example cells carrying DA, 5-HT, and NE sensors (same cells as in **Supplementary Video 7**), and individual active cells from this recording are plotted in Figure 2B**, right heatmap**. Composition of the solution in each well is shown in the lower left. Time is shown as hh:mm:ss, scale bar: 100 µm. See Figures 2 **and S4** for additional details.

**Supplementary Video 9. MORSE recording from the lateral ventricle CSF of an awake mouse with a four-sensor probe.** https://www.dropbox.com/scl/fi/sbtu3d9wikpor04vv8sqd/sv9.avi?rlkey=rwlt8nulhr5nhu2hlopwqjpkk&st=ujvms36v&dl=0

A representative MORSE probe carrying four GRAB sensor cell types is recorded while implanted in the lateral ventricle CSF of an awake mouse. The mouse received IP injections of OXT and AVP (1 mg/kg each in 200 µL PBS) followed by an ICV injection of 5-HT (150 nmol in 30 µL aCSF, via the contralateral cannula). Images at each timepoint demonstrate the mean raw GRAB sensor fluorescence signal (no normalization) of three consecutive imaging volumes (1.9 sec/vol) averaged over three adjacent image planes spanning a total of ∼30 µm in gel thickness (∼15% of total thickness containing sensor cells) (**left**) and the corresponding ΔF/F_0_ signal (**right**). White arrowheads indicate example cells carrying OXT, AVP, and 5-HT sensors (same cells as in **Supplementary Video 10**), and a subset of individual active cells from this recording is plotted in Figure 3B**, left heatmap**. Note that 5-HT sensor signal begins to increase before ICV 5-HT delivery, possibly reflecting endogenous serotonin release into the CSF (demonstrated further in Figures 4A-D). Injection times are shown in the lower left. Time is shown as hh:mm:ss, scale bar: 100 µm. See Figures 3-4 for additional details. The same MORSE probe is shown undergoing post hoc dipping characterization in **Supplementary Video 10**.

**Supplementary Video 10. Post hoc dipping of a four-sensor MORSE probe following an *in vivo* recording in live mouse CSF.** https://www.dropbox.com/scl/fi/zg16vnd1rtrkm11kmtdad/sv10.avi?rlkey=k1rswu8v250a6fp7vgk3pjcfk&st=fe977q9u&dl=0

The same MORSE probe as in **Supplementary Video 9**, with the image rotated and warped for alignment, is shown being dipped into solutions containing near-saturating concentrations of four ligands in PBS: vasopressin (AVP, 1 µM), oxytocin (OXT, 1 µM), cholecystokinin (CCK, 1 µM), and serotonin (5-HT, 1 µM). Each dip lasted 5 minutes. Images at each timepoint demonstrate the mean raw GRAB sensor fluorescence signal (no normalization) of three consecutive imaging volumes (1.9 sec/vol) averaged over three adjacent image planes spanning a total of ∼30 µm in gel thickness (∼15% of total thickness containing sensor cells) (**left**) and the corresponding ΔF/F_0_ signal (**right**). White arrowheads indicate example cells carrying OXT, AVP, and, 5-HT sensors (same cells as in **Supplementary Video 9**), and a subset of individual active cells from this recording is plotted in Figure 3B**, right heatmap**.. Composition of the solution in each well is shown in the lower left. Time is shown as hh:mm:ss, scale bar: 100 µm. See Figures 3-4 for additional details.

**Supplementary Video 11. GCaMP6f recording from choroid plexus epithelium during ICV injection of serotonin.** https://www.dropbox.com/scl/fi/xh655iqanpw97sli5602o/sv11.avi?rlkey=cnu1dg2ygq0cditewfxd4rpfg&st=bk4u9ena&dl=0

A representative recording of the choroid plexus epithelium in an awake *FoxJ1-Cre::Ai95D* mouse is shown. The mouse received an ICV injection of 5-HT (75 nmol in 15 µL aCSF over 3 min, via the contralateral cannula). Images at each timepoint demonstrate the mean raw GCaMP6f sensor fluorescence signal (no normalization) of ten consecutive imaging volumes (4.3 sec/vol) maximum-projected through the full thickness of the epithelial sheet (∼100 µm total thickness including two layers of epithelium with stroma and vasculature in between). Injection time is shown in the lower left. Time is shown as hh:mm:ss, scale bar: 50 µm. See Figure 4G for additional details.

**Supplementary Video 12. GCaMP6f recording from choroid plexus epithelium during IP injection of serotonin.** https://www.dropbox.com/scl/fi/h6tvtxqbjj5zyyj2ff0zx/sv12.avi?rlkey=mzgioauznm50xgil7tlkeuh5b&st=4tzokmb9&dl=0

A representative recording of GCaMP6f calcium signals restricted to the choroid plexus epithelium in an awake *FoxJ1-Cre::Ai95D* mouse is shown. The mouse received an IP injection of 5-HT (2 mg/kg in 200 µL saline). Images at each timepoint demonstrate the mean raw GCaMP6f sensor fluorescence signal (no normalization) of ten consecutive imaging volumes (4.3 sec/vol) maximum-projected through the full thickness of the epithelial sheet (∼100 µm total thickness including two layers of epithelium with stroma and vasculature in between). Injection time is shown in the lower left. Time is shown as hh:mm:ss, scale bar: 25 µm. See Figure 4H for additional details.

**Supplementary Video 13. GRAB-5-HT2h recording from choroid plexus epithelium during IP injection of serotonin.** https://www.dropbox.com/scl/fi/62j9wkwyk08h47o1czxi0/sv13.avi?rlkey=jcu8c4326oz271ca5ntev80vs&st=4h47fb7a&dl=0

A representative recording of the choroid plexus epithelium in an awake wild-type mouse, 7 weeks after ICV injection of AAV2/5 carrying the GRAB-5-HT2h construct, is shown. The mouse received an IP injection of 5-HT (2 mg/kg in 200 µL PBS). Images at each timepoint demonstrate the mean raw GRAB-5-HT2h fluorescence signal (no normalization) of ten consecutive imaging volumes (4.3 sec/vol) maximum-projected through the full thickness of the epithelial sheet (∼100 µm total thickness including two layers of epithelium with stroma and vasculature in between). Note that increases in fluorescence are evident on the apical (CSF-facing) side of the epithelial cells, where stronger baseline sensor expression is observed. Injection time is shown in the lower left. Time is shown as hh:mm:ss., scale bar: 25 µm. See Figure 5 for additional details.

